# Downregulating α-synuclein in iPSC-derived dopaminergic neurons mimics electrophysiological phenotype of the A53T mutation

**DOI:** 10.1101/2022.03.31.486582

**Authors:** Philipp Hornauer, Gustavo Prack, Nadia Anastasi, Silvia Ronchi, Taehoon Kim, Christian Donner, Michele Fiscella, Karsten Borgwardt, Verdon Taylor, Ravi Jagasia, Damian Roqueiro, Andreas Hierlemann, Manuel Schröter

## Abstract

Parkinson’s disease (PD) is a common debilitating neurodegenerative disorder, characterized by a progressive loss of dopaminergic (DA) neurons. Mutations, gene dosage increase, and single nucleotide polymorphisms in the α-synuclein-encoding gene *SNCA* either cause or increase the risk for PD. However, neither the function of α-synuclein in health and disease, nor its role throughout development is fully understood. Here, we introduce *DeePhys*, a new tool that allows for data-driven functional phenotyping of neuronal cell lines by combining electrophysiological features inferred from high-density microelectrode array (HD-MEA) recordings with a robust machine learning workflow. We apply *DeePhys* to human induced pluripotent stem cell (iPSC)-derived DA neuron-astrocyte co-cultures harboring the prominent *SNCA* mutation A53T and an isogenic control line. Moreover, we demonstrate how *DeePhys* can facilitate the assessment of cellular and network-level electrophysiological features to build functional phenotypes and to evaluate potential treatment interventions. We find that electrophysiological features across all scales proved to be highly specific for the A53T phenotype, enabled to predict the genotype and age of individual cultures with high accuracy, and revealed a mutant-like phenotype after downregulation of α-synuclein.

## Introduction

Neurological disorders are a leading cause of disability in today’s aging societies. One of the fastest growing neurological diseases is Parkinson’s disease (PD) (Bloem et al., 2021). PD patients suffer from a progressive loss of dopaminergic (DA) neurons within the substantia nigra *pars compacta* (SNc), which results in the development of tremor, shaking, rigidity and slowness at initiating movement (Jankovic, 2008). PD is currently not curable, hence, there is an urgent need to better understand the precise pathophysiology of PD, and how potential treatments could rescue neuronal physiology.

While the causes of PD are still debated (Dickson, 2018), a number of genetic mutations have been associated with the disease (Klein & Westenberger, 2012). Neuropathological investigations have found α-synuclein (α-syn) protein aggregates to be a key feature of the disease, and excessive levels of α-syn oligomers have been implicated in both sporadic and familial forms of PD (Mezey et al., 1998; Polymeropoulos et al., 1997; Spillantini et al., 1997). Several mutations modulating the aggregation rate of the α-syn protein have been identified in the α-syn gene (*SNCA*) (Flagmeier et al., 2016), and toxic gain-of-function mutations of the resulting oligomers have been associated with mitochondrial defects (Plotegher et al., 2014), membrane damage (Chaudhary et al., 2016), and synaptic dysfunction (Choi et al., 2013; Kouroupi et al., 2017). Among these mutations, the A53T variant has one of the strongest effects on the initiation and spreading of α-syn aggregations (Conway et al., 2000; Flagmeier et al., 2016) and results in an autosomal dominant form of familial PD (Polymeropoulos et al., 1997). Previous studies have identified distinct molecular phenotypes in human neurons harboring this mutation, including impaired bioenergetics (Zambon et al., 2019, Ryan et al., 2013) and altered synaptic connectivity (Kouroupi et al., 2017). However, these results were mainly based on immunohistochemical and transcriptomic data, while a more systematic functional characterization of A53T mutant α-syn expressing neuronal cultures is still lacking.

Despite considerable progress in our understanding of the physiological properties of α-syn, such as its predominantly presynaptic localization (Galvin et al., 2001; Withers et al., 1997) and its interactions with highly curved membranes and synaptic proteins (Burré et al., 2010; Sun et al., 2019), the impact of α-syn on synaptic function is still not fully understood and warrants further investigation (Burré, 2015). Similarly, the precise pathomechanisms by which α-syn accumulation affects the likelihood to develop PD remain largely unknown. As a result of this knowledge gap, the success of therapeutic strategies, such as the downregulation of α-syn expression (Dehay et al., 2015; Fields et al., 2019), is difficult to predict, as the latter might be accompanied by severe side effects caused by the loss of physiological α-syn function.

The advent of induced pluripotent stem cells (iPSCs) has provided unprecedented access to human neuronal cells and means to study specific mechanisms and developmental pathways that give rise to a range of neurological disorders *in vitro* (Hu et al., 2020; Wu et al., 2019). Previous studies have used human iPSCs-derived DA neurons to study the effect of PD-relevant mutations on cellular integrity, survival, morphology and biochemical or metabolic alterations (Byers et al., 2011; Hu et al., 2020; Laperle et al., 2020; Nguyen et al., 2011; Woodard et al., 2014). However, due to the documented role of α-syn in synaptic physiology (Bellani et al., 2010; Cheng et al., 2011; Sulzer & Edwards, 2019) more detailed electrophysiological analyses of human DA neuronal networks harboring pathogenetic *SNCA* mutations could provide important additional insights regarding its complex functional role and the interaction of altered synaptic activity with other PD-associated disease mechanisms.

High-density microelectrode arrays (HD-MEAs) are a technology platform that is suitable to probe the development of disease-specific electrophysiological phenotypes *in vitro*: they allow to capture the electrical activity of several hundreds of neurons simultaneously, thus providing the high yield and throughput needed for high-content screenings (Abbott et al., 2020; Berdondini et al., 2009; Eversmann et al., 2003; Müller et al., 2015; Tsai et al., 2017; Yuan et al., 2020). In addition, HD-MEAs allow for the electrophysiological characterization of neurons across scales, that is, at the subcellular, cellular, and network-level (Obien et al., 2014; Spira & Hai, 2013). Despite being commercially available for several years now, the rich details provided by HD-MEAs have often remained untapped (Battaglia et al., 2020; Hiramatsu et al., 2021; Schenke et al., 2021). While generic readouts, such as multi-unit activity can undoubtedly give some insight into culture development and viability, more comprehensive longitudinal readouts that provide detailed physiological insights into the multifaceted mechanisms of neurodegeneration and disease-relevant functional aspects are needed. Detailed investigations could include, for example, the study of ion-channel dependent changes in the action potential (AP) waveform of neurons, which have been identified to be a common pathognomonic feature of neurological diseases (Brenes et al., 2015; Kopach et al., 2021).

Systematic analysis of single-neuron waveform features and network-related measures requires spike sorting of the recorded data (Rey et al., 2015). Spike sorting is a computationally expensive process that has, however, been significantly facilitated by an increasing number of well-documented algorithms (Diggelmann et al., 2018; Stringer et al., 2019; Yger et al., 2018) and helper packages such as SpikeInterface (Buccino et al., 2020) and SpikeForest (Magland et al., 2020). While the barrier of entry has been lowered considerably for researchers from outside the electrophysiology domain, there is still a lack of easy-to-use software that provides an integrated approach to infer, quantify and compare single-neuron and network features.

Here, we introduce *DeePhys*, a novel modular analysis tool that we use in a proof-of-concept application to quantitatively infer the functional phenotypes of isogenic wild-type (WT) and A53T mutant α-syn expressing human iPSC-derived DA neuronal cultures across scales (single-cell and network), during development, and under specific treatment conditions. We demonstrate that *DeePhys* provides intuitive insights into a set of specific features that are reliable predictors for functional phenotypes and their development during culture maturation. Using over 30 extracellular electrophysiological features, inferred from over 130.000 spike-sorted neurons/units across 5 weeks *in vitro*, we thoroughly characterized DA neurons and found consistent differences across all feature groups, i.e., differences in waveform, single-cell activity, and network activity of neurons in health and disease. Finally, we applied *DeePhys* to probe the impact of locked nucleic acid (LNA)-mediated downregulation of α-syn expression on electrophysiological properties and evaluated the potential of this treatment approach to correct the phenotype of A53T mutant α-syn expressing DA neurons.

## Results

Inferring comprehensive and reproducible functional phenotypes from human induced pluripotent stem cells (iPSC)-derived cellular models of neurological disorders is fundamental in order to effectively develop and evaluate potential pharmacological interventions in the pre-clinical setting. In the present study, we used previously established protocols to culture (Ronchi et al., 2021; see **Methods *Cell culture and plating***) and electrophysiologically characterize human dopaminergic (DA) neurons on high-density microelectrode arrays (HD-MEAs; see **Methods *High-density microelectrode array recordings***), to demonstrate the potential of this combination for systematic functional phenotyping. The main dataset consisted of healthy (WT; N=18) and A53T mutant DA (N=19) neuronal cultures, which were pooled over two experiments. A third dataset consisting of 8 WT and 8 A53T cultures was used to study the effects of downregulating α-synuclein (α-syn) by chronic treatment with locked nucleic acid probes (LNAs) from day *in vitro* (DIV) 1 onwards (see **Methods *Locked nucleic acid administration***; **Supplemental Table 1**). An overview of the experiment is shown in **Figure 1**.

**Figure 1.**
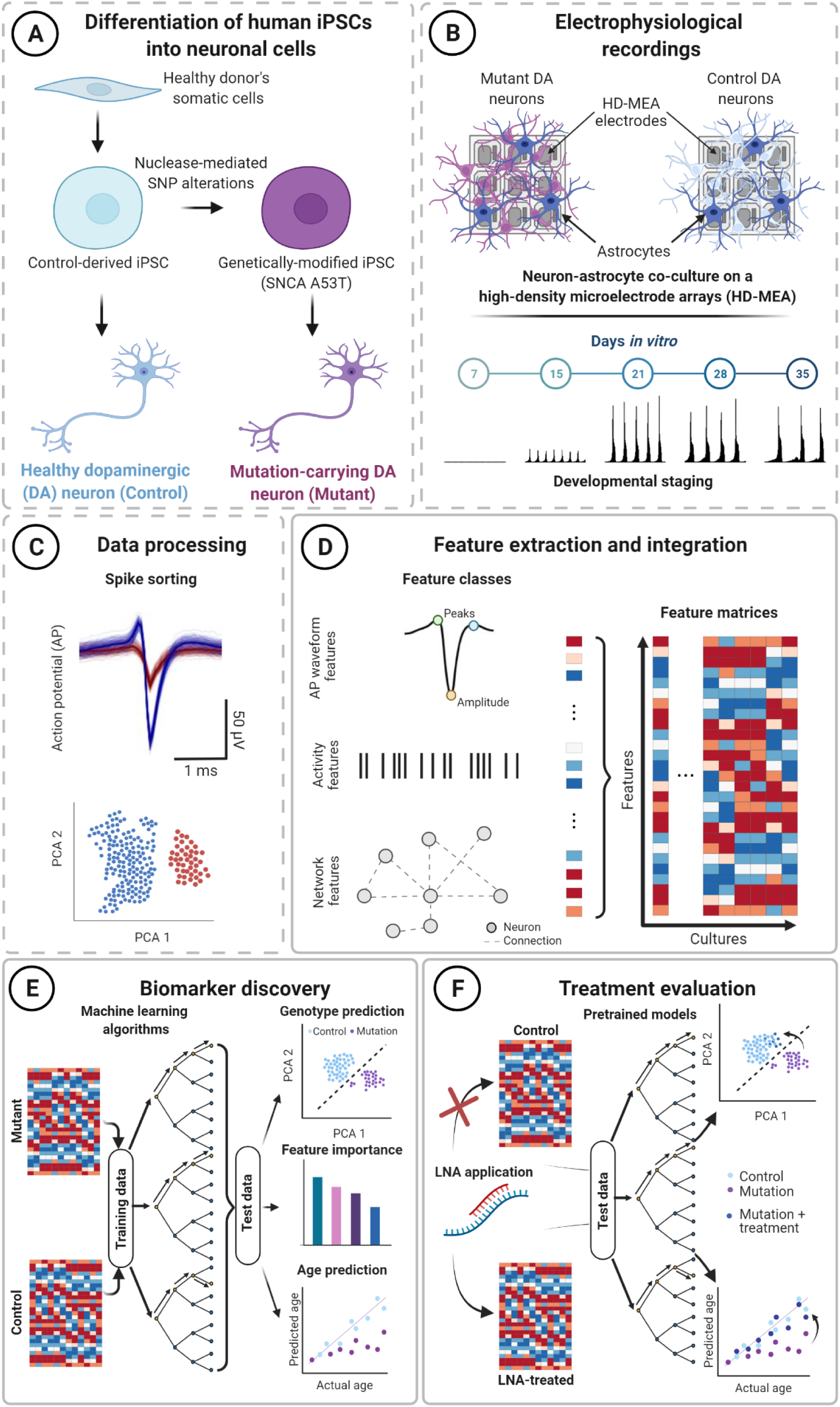
Schematic of the *DeePhys* analysis pipeline. **(A)** Human iPSC-derived dopaminergic (DA) neurons and isogenic DA mutant neurons (*SNCA*-A53T) were plated on high-density microelectrode arrays (HD-MEAs) for electrophysiological characterization. (**B**) Control and mutant DA neurons were co-cultured with human control astrocytes and electrophysiologically tracked for up to 35 days *in vitro* (DIV). **(C)** The *DeePhys* pipeline begins after the HD-MEA network data underwent spike sorting, here performed with SpyKING Circus (Yger et al., 2018), see **Figure 1D-F**. Spike sorting is required to demix the HD-MEA data and to assign the recorded activity to single neurons/units. **(D)** Spike-sorted data enable the inference of detailed electrophysiological metrics, including temporal and spatial aspects of neuronal activity at the single-unit and network level. In *DeePhys*, this rich data is combined and used to build feature vectors/matrices to systematically assess mutant and control cultures functionally. **(E)** The inferred features can be used for data-driven classification analyses (see upper inset: *Genotype prediction*), to find particularly informative features (see middle inset: *Feature importance*), and to perform age regression analysis (see lower inset: *Age prediction*) to quantify alterations in the developmental trajectories of mutant and control DA neuron cultures. **(F)** Pre-trained models can be used to evaluate treatment effects on the functional phenotype. In the present study, for example, we studied the effects of reducing the expression of α-syn through the application of a locked nucleic acid (LNA) for phenotype rescue. Panels with dashed outlines represent steps that precede the analysis with *DeePhys* (**Figure 1A-C**), while a solid outline indicates the core steps of the *DeePhys* pipeline (**Figure 1D-F**).

In order to obtain robust functional phenotypes from HD-MEA recordings, and to probe the effects of treatment interventions, we developed *DeePhys. DeePhys* consists of a MATLAB-based analysis pipeline that includes three main modules: The analysis pipeline starts with a processing module that extracts extra-cellular features from spike-sorted HD-MEA data (**Figure 1D;** see **Methods *Feature Extraction*** for a full list). The inferred properties consist of action potential (AP) waveform metrics, single neuron firing statistics, and metrics that quantify the regularity and synchronicity of population activity. The second module aggregates these features into functional phenotypes and aims at providing an analytical workflow to address key questions that are relevant for many iPSC-disease modeling studies (**Figure 1E**): Is there a functional phenotype that allows for discriminating different cell lines (e.g., according to their genotypes)? What are the metrics that are most discriminative? How do phenotypes develop and when are they most apparent? Finally, the last *DeePhys* module allows quantification of the treatment success following a pharmacological intervention (see **Figure 1F**). The MATLAB functions for each module are described in **Supplemental Table 2**.

### Immunocytochemical and electrophysiological characterization of human DA neuron cultures

We used a previously established protocol to plate differentiated, highly-enriched, human iPSC-derived DA neurons carrying a heterozygous A53T mutation and an isogenic wild type (WT) control line on HD-MEAs and cover slips (control experiments; Ronchi et al., 2021). DA neurons were co-cultured with human control astrocytes and plated on HD-MEAs at a ratio of 5:1 (DA neurons:astrocytes, Figure 1A-B). We confirmed the presence and development of DA neurons and astrocytes in control experiments by immunocytochemical stainings for tyrosine hydroxylase (TH) expression, microtubule-associated protein 2 (MAP2) and glial fibrillary acidic protein (GFAP) at DIV 21 (Figure 2A, Figure 2 - figure supplement 1). Both WT and A53T DA neuron-astrocyte co-cultures showed robust outgrowth and neuritic arborization of TH^+^ neurons, and a wide coverage of GFAP^+^ astrocytic processes. Despite this apparent similarity, quantifications of TH^+^ and MAP2^+^ nuclei revealed differences in the composition of both lines, such as a decreased number of TH^+^ neurons per imaged field (WT: 122.8 ± 66.9; A53T: 80 ± 40.7; mean ± SD) and a decrease in the TH^+^/MAP2^+^ ratio (WT: 49.2 ± 14.0%; A53T: 43.9 ± 11.3%) (for statistical details see **Supplemental Table 3-4**).

**Figure 2.**
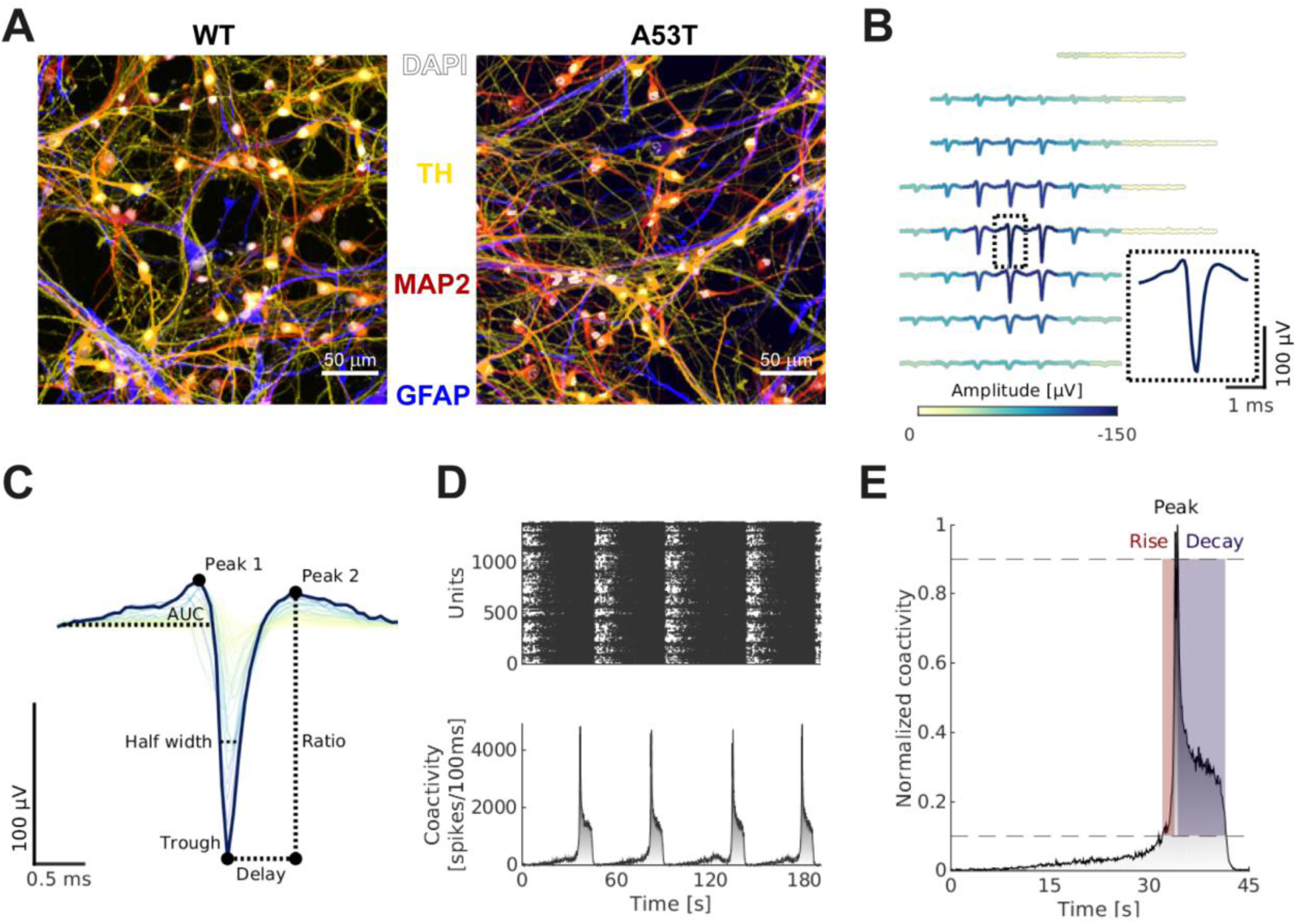
Growing human iPSC-derived DA neurons on high-density micro-electrode arrays. (**A**) Control (left panel) and mutant (right panel) human iPSC-derived DA neurons, co-cultured with human astrocytes, showed robust TH expression at days *in vitro* (DIV) 21 (DAPI: gray, TH: yellow, MAP2: red, GFAP: blue). Cultures of both genotypes did not display any clear morphological differences at this time (Figure 2 - figure supplement 1). (**B**) Example electrical footprint (EF), inferred by spike-triggered averaging using the spike-sorted activity. The largest negative amplitude waveform of an EF, its ‘reference waveform’, was used to infer various AP waveform features. Coloring of EF waveforms indicates their respective maximum amplitudes. (**C**) The EF depicted in (**B**) in the temporal domain and some features that were inferred from the reference waveform (in dark blue), including the half-width and the trough-to-peak delay. Coloring of the waveforms is according to the respective trough amplitudes. (**D**) HD-MEA network recordings allowed for systematic characterization of DA population activity. The upper panel shows a representative spike raster plot with activity recorded from a more mature DA WT culture (DIV 35). Dots represent spike-sorted action potentials. The lower panel displays the corresponding binned co-activity of all neurons (i.e., number of spikes/100ms bin). Overall, both WT and A53T DA neuronal cultures developed robust network burst activity. (**E**) Zoom in on a single network burst. As part of *DeePhys*, the shape and dynamics of individual bursts were further characterized, for example, by determining the duration of bursts, their rise and decay times and the interburst intervals.

HD-MEA recordings of spontaneous neuronal activity started one week after plating and continued for up to 5 weeks (Figure 1B). Longitudinal HD-MEA recordings consisted of 15 minutes of spontaneous network activity, following an array-wide Activity Scan to select the most active neurons on the array (see **Methods *High-density microelectrode array recordings*** for details). Spike-sorting was performed using SpyKING Circus (Yger et al., 2018), a density-based clustering and template matching algorithm, with parameter settings adapted to the data at hand (see **Supplemental Table 5**). It is noteworthy, that despite robust overall activity levels at later stages of development (e.g., an average mean firing rate of 1.04Hz at DIV 28), iPSC-derived DA neurons showed relatively small spike amplitudes (e.g., an average spike amplitude of 36.2μV at DIV 28), which made spike sorting challenging. Following spike sorting, we restricted our analysis to units/neurons that showed minimal refractory period violations (<2%) and robust baseline activity (0.1Hz, see **Methods *Spike sorting*)**.

For each unit, we inferred 17 features that describe the neuronal activity at the cellular level (see **Methods *Feature extraction*** and Figure 2B-C). *Single-cell waveform features* were derived from the electrical footprints (EFs) of spike-sorted units, which represent the extracellular electrical potential landscape of a specific unit as recorded by the HD-MEA electrodes (Figure 2B). Here, we used the spike-triggered multi-electrode waveform signal (template), as generated during the spike sorting step (Figure 2D). Templates were generated from the highpass filtered (300Hz) signal, using a minimum of 90 and up to a maximum of 500 randomly sampled spikes. Single-cell waveform features were then extracted from the signal of the “reference electrode” of the EF, i.e., from the electrode that recorded the largest negative waveform amplitude (Figure 2C). *Activity-based single-cell features* were inferred from the spike times of individual units, describing the temporal pattern and total extent of DA neuron activity. Additionally, we inferred 14 *network features* that describe the population activity over all spike-sorted units of a culture (see **Methods *Feature extraction*** and Figure 2D-E), resulting in a total of 31 features.

Overall, we found that the number of spike-sorted neuronal units and their respective firing rates increased significantly during development from week 1 (number of neurons across both WT and A53T cultures at week 1: 326 ± 111; firing rate: 0.38 ± 0.28 Hz) until week 3/4 (number of neurons across both WT and A53T cultures at week 3/4: 647 ± 143, 1.04 ± 0.52 Hz; both p<0.001, linear mixed-effect model (LMM)), but did not differ significantly between WT and A53T mutant cell lines (genotype effect: p=0.609, p=0.496; genotype-time interaction: p=0.163, p=0.803, LMM; for details see **Supplemental table 1**).

### *DeePhys* allows for highly accurate predictions of DA neuron culture type

To probe whether individual single-cell and network electrophysiological properties differed between WT and A53T DA neuron cultures, we trained random forest (RF) classifiers for each metric (Figure 3). The relative importance of each time point for these predictions was measured by the permutation predictor importance (Breiman, 2001; Figure 3A). Differences in the development of the electrophysiological properties of each cell line were further assessed statistically using LMMs (Figure 3; **Supplemental Table 6**). A comparison of RF classifier performance to other classification algorithms is provided in the supplement (Figure 3 - figure supplement 1).

**Figure 3.**
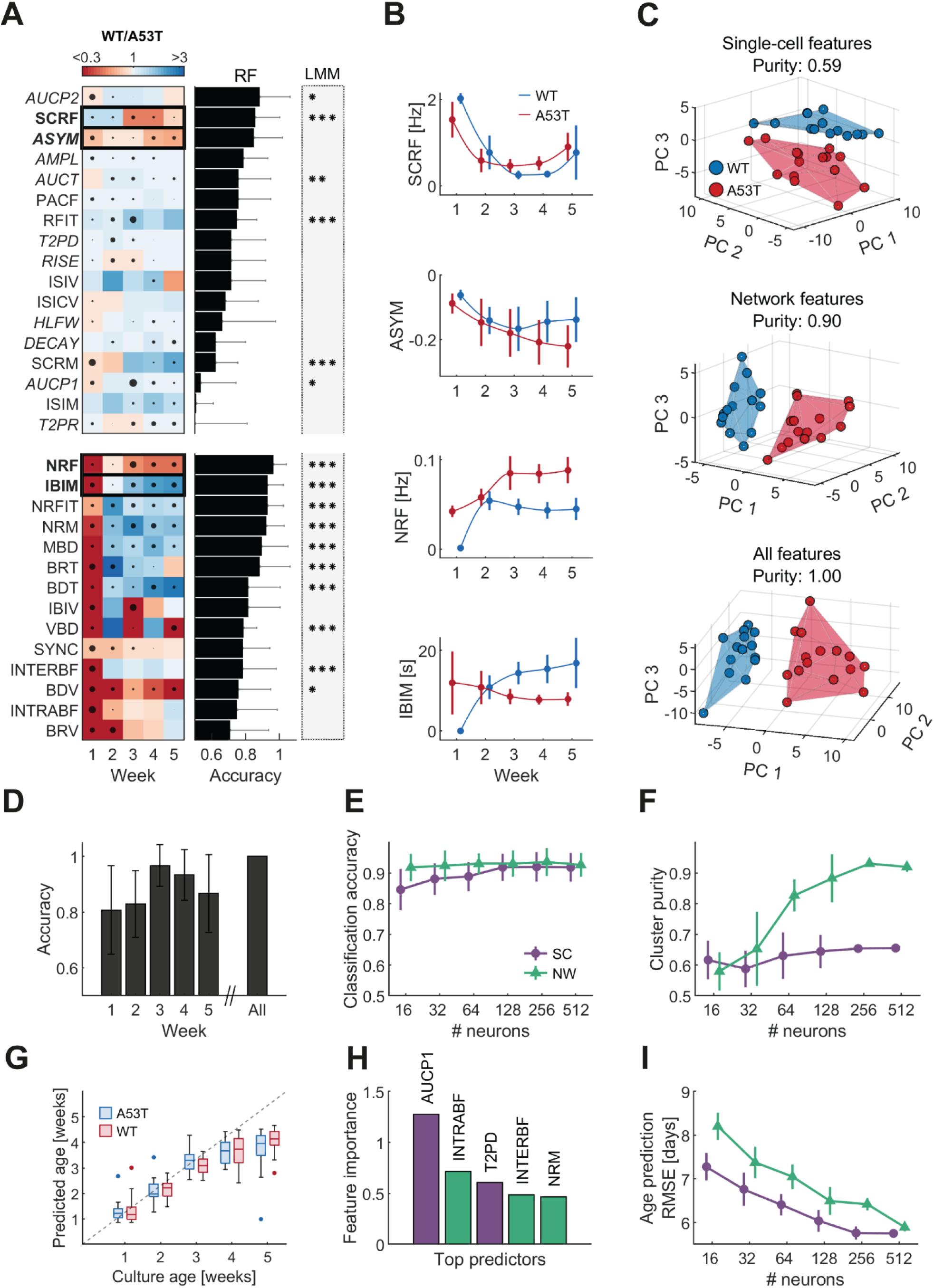
A53T DA neuron cultures exhibit age-dependent alterations at the single-cell and network level. **(A)** Electrophysiological single-cell and network feature differences in the developmental trajectories of both genotypes across five weeks of development *in vitro* (N_WT_=18, N_A53T_=19 cultures), as inferred with the *DeePhys* pipeline. Heatmaps indicate differences between genotypes at a given recording time point (weeks 1-5; ratio WT/A53T: blue indicates a higher value in WT cultures, red indicates a higher feature value in A53T cultures). The horizontal bar plots display the accuracies of Random Forest (RF) classification models, trained with the respective features as input (mean ± SD from 5-fold cross-validation (CV)). The size of the black dots within each panel of the heatmap indicates the relative *predictor importance* (Breiman, 2001) at the respective time point. Additionally, we computed linear mixed-effect models (LMM) to probe if features display a significant genotype difference or genotype-time interaction (see gray-shaded column for LMM results; asterisks indicate significance: *p<0.05, **p<0.01, ***p<0.001, after Bonferroni correction; see **Supplemental Table 6** for details). Overall, network-level features yielded more pronounced differences between phenotypes than single-cell features.This was also underlined by a larger number of features that yielded high prediction accuracy. **(B)** Example trajectories of four highly predictive features, including single-cell regularity frequency (SCRF), asymmetry (ASYM) and mean inter-burst-interval (IBIM) (error bars represent mean ± SD values; N_WT_=18, N_A53T_=19 cultures; colors: blue = WT, red= A53T DA neuronal cultures). **(C)** Principal component analysis (PCA) results demonstrate that WT/A53T DA neuronal cultures can be grouped based on inferred single-cell (upper panel) and network features (middle panel). For visualization, we depict here the first three PCs. The lower plot displays PCA results from a combination of single- and network-level features (Colors: blue = WT, red= A53T DA neuronal cultures). We also report the *cluster purity* of the k-means clustering (k=2) for each PCA result. **(D)** Phenotypic differences across development were assessed by training RF classification models. Classification accuracy peaked at week 3 and decreased thereafter (mean ± SD from 5-fold CV). Combining feature values across all weeks resulted in a perfect classification accuracy (bar on the right). A comparison of RF classification performance to other classifiers is provided in Figure 3 - figure supplement 1. **(E)** Between phenotype RF classification results from a subsampling analysis, i.e., analyses with single-cell or network-level features from a smaller subset of neurons. We find a small drop in prediction accuracy for single-cell (SC) features, while the prediction accuracy for network (NW) features remains largely unchanged (colors: purple = network features, green = single-cell features). **(F)** Subsampling results for cluster purity (see also panel **C**): cluster purity for NW features decreased at lower neuron numbers but remained relatively stable for SC features. **(G)** RF regression analysis was used to predict culture age (including all feature values from one recording time point). Until week 4, the regression analysis remained accurate (all root mean square error (RMSE) < 4.5 days), but culture age was underestimated at week 5 (RMSE = 9.4 days; boxes visualize the median; lower, and upper quartiles, whiskers indicate non-outlier minimum and maximum values; dots represent outlier values). **(H)** Representation of the most important features for age prediction (colors: purple = NW features, green = SC features). **(I)** Subsampling results for the age prediction indicate altered performance for the RF regression analysis, if only smaller subsets of neurons were used (i.e., the average RMSE increased for fewer neurons).

### WT and A53T DA neuron cultures show distinct functional phenotypes

WT and A53T DA neuron cultures showed marked differences across all feature classes: Extracellular AP waveform shapes (Figure 3A, upper panel, waveform features in italics) differed between WT and A53T, as several waveform features yielded highly accurate classification results (e.g., area under the curve of peak 2; AUCP2, accuracy: 0.89 ± 0.18, p<0.05). Single-cell features also differed consistently between genotypes at the activity level, particularly those metrics that describe the temporal regularity of DA neuron firing (e.g., single-cell regularity frequency; SCRF, accuracy: 0.86 ± 0.14, p<0.001, Figure 3B). Network features differed even more between the two genotypes (Figure 3A, lower panel) and showed differences that became apparent already early during development: while A53T cultures demonstrated high levels of correlated spontaneous activity one week after plating, no bursts were detectable in WT cultures at that time. This translated into differences in several other network-related metrics throughout development, most prominently, the temporal regularity of network oscillations, here termed *network regularity frequency* (NRF). The NRF was highly consistent among cultures (accuracy: 0.97 ± 0.07, p<0.001) and displayed a characteristic developmental trajectory for control and mutant cultures (Figure 3B, third panel): While the NRF increased continuously in A53T cultures, it decreased after the emergence of synchronous activity at week 2 in WT cultures.

Overall, many metrics across all feature groups displayed strong phenotypic differences and high feature importance values at week 1 and 3, which was often paralleled by a cross-over around the second week (e.g., IBIM, Figure 3B, bottom panel). In Figure 3C, we depict how well each feature class differentiated the two genotypes in the principal component (PC) space. Principal component analysis was performed on single-cell features (top panel), network-level features (middle panel), as well as on the combination of both (lower panel). In all three cases, the first three PCs visually separated WT and A53T cultures. To quantify these differences, k-means clustering (k=2) was performed on the PCs explaining 95% of the variance. Cluster purity, a measure of the separability of clusters, increased from single-cell features (0.59, 16 PCs) to network features (0.90, 17 PCs) and reached maximum values for a combination of both (1.00, 19 PCs). These results highlight that network features were the more consistent predictors of the respective phenotype, but also, that single-cell features contributed additional important information.

### Phenotypic differences are robust to subsampling

Next, we quantified differences in the functional phenotypes of WT and A53T cultures at individual developmental time points (i.e., weeks 1-5, Figure 3D). Comparing RF classification results with feature values, computed for each week, showed a marked peak in prediction accuracy at week 3; this resembled the development in predictor importance values after the cross-over after the second week *in vitro* (Figure 3A, dot sizes correspond to feature importance values). Overall, clear phenotype differences across all developmental time points were evident, and RF classifications achieved accuracy values well above 0.8. Note, this result held true even when each feature group was considered individually (Figure 3 - figure supplement 2). A perfect classification accuracy was possible when features from all time points were combined, which indicates that insights on the developmental trajectories can further improve the performance.

We further quantified the robustness of the RF genotype prediction by performing a subsampling analysis. Here, the number of putative neurons per culture for the classification task were varied from 16 to 512 neurons (neurons were selected randomly; accuracy values are averaged over 25 iterations) (Figure 3E). While predictions based on both feature classes (i.e., single-cell and network level) remained highly accurate throughout all subsampling steps (>0.8 for single-cell; >0.9 for network), we found that predictions based on single-cell features decreased slightly in accuracy at lower neuron numbers. The robustness of the k-means clustering was assessed by calculating the cluster purity across all subsampling iterations for both feature classes: While clustering based on single-cell features was hardly affected and achieved similar cluster purity values across all subsampling iterations, cluster purity values for network features dropped by more than 0.3 (Figure 3F).

### Single-cell waveform and network features allow for robust age prediction

Finally, we probed how accurately electrophysiological features could predict the age of WT/A53T cultures and performed age prediction using RF regression models (Figure 3G). Culture ages could be accurately predicted until week 4 (all root mean square errors (RMSE) < 4.5 days) but were consistently underestimated at week 5 (RMSE = 9.4 days). This finding likely indicates a plateau in the developmental trajectory after 4 weeks *in vitro*. Features describing the AP waveform (e.g., the area under the curve of peak 1 (AUCP1)) and network-level activity (e.g., the intra-burst firing rate (INTRABF)) were most predictive (Figure 3H). We also evaluated the robustness of the age regression (subsampling analysis) and found that the performance dropped when the number of neurons was reduced (Figure 3I). Taken together, our results highlight the importance of sampling from a sufficiently large number of neurons for functional phenotyping and the potential of RF regression analysis to assess the development of neuronal cultures *in vitro*.

### LNA application effectively downregulates α-synuclein expression in DA neurons

As reducing the expression of α-syn has been proposed as a therapeutic option to slow the progression of PD (Dehay et al., 2015; Fields et al., 2019), we applied a locked nucleic acid (LNA) probe to target the α-syn mRNA (Petersen and Wengel, 2003). In order to validate the efficacy of the LNA treatment, we performed a control experiment to measure α-syn levels in untreated, non-targeted LNA (ntLNA)-treated, and *SNCA*-targeting LNA-treated cultures at DIV 21 using a Homogeneous Time Resolved Fluorescence (HTRF) assay (N=3 cultures per condition x 3 technical replicates) (Degorce et al., 2009). Two-factor ANOVA revealed a significant effect of the LNA treatment (F(2,11) = 885.3, p<0.001) on total α-syn levels, but no significant difference between both genotypes (F(1,11) = 0.05, p=0.820). Post-hoc multiple comparisons tests showed a significant decrease of α-syn levels in LNA-treated WT (p<0.001, Tukey’s test) and A53T cultures (p<0.001), compared to the untreated and ntLNA-treated controls (Figure 4C-D; **Supplemental table 7**). Next, we performed automated immunocytochemical analysis (quantification of average intensities) to assess the subcellular localization of α-syn and phosphorylated α-syn (p-syn) in TH^+^ neurons at DIV 21. In accordance with the HTRF results, we observed a pronounced reduction of both α-syn (Figure 4A) and p-syn (Figure 4B) levels after LNA application. Two-factor ANOVA revealed a significant treatment effect on somatic α-syn (F(2,102) = 691.9, p<0.001) and p-syn levels (F(2,102) = 180.1, p<0.001, N=3 cultures per condition, 54 fields per culture; Figure 4C-D; **Supplemental table 8-9**). Post-hoc multiple comparisons tests showed a significant decrease only in the LNA-treated conditions of both genotypes (both p<0.001, Tukey’s test). Moreover, the genotype had a significant effect on somatic p-syn levels (F(1,102) = 11.2, p=0.001, Two-factor ANOVA), but not on somatic α-syn levels (F(1,102) = 2.5, p=0.116), indicating higher levels of α-syn accumulation in A53T cultures. Quantification of α-syn levels in the neurites of TH^+^ neurons also revealed a significant treatment effect (F(2,102) = 182.6, p<0.001, Two-factor ANOVA), a significant effect of the genotype (F(1,102) = 29.0, p<0.001), and a significant interaction between the two factors (F(2,102) = 3.9, p=0.023). Additionally, quantification of p-syn levels in neurites demonstrated a significant reduction of p-syn in LNA-treated cultures (F(2,102) = 142.4, p<0.001), but no significant effect of the genotype (F(1,102) = 0.4, p=0.530; **Supplemental table 10-11**). As mentioned above, we found differences in the TH^+^/MAP2^+^ ratio between the cell lines (**Supplemental Table 3-4**). Taken together, these data suggest a lack of α-syn in the neurites of A53T DA neurons, which might be the result of an increase in α-syn phosphorylation and accumulation in the soma.

**Figure 4.**
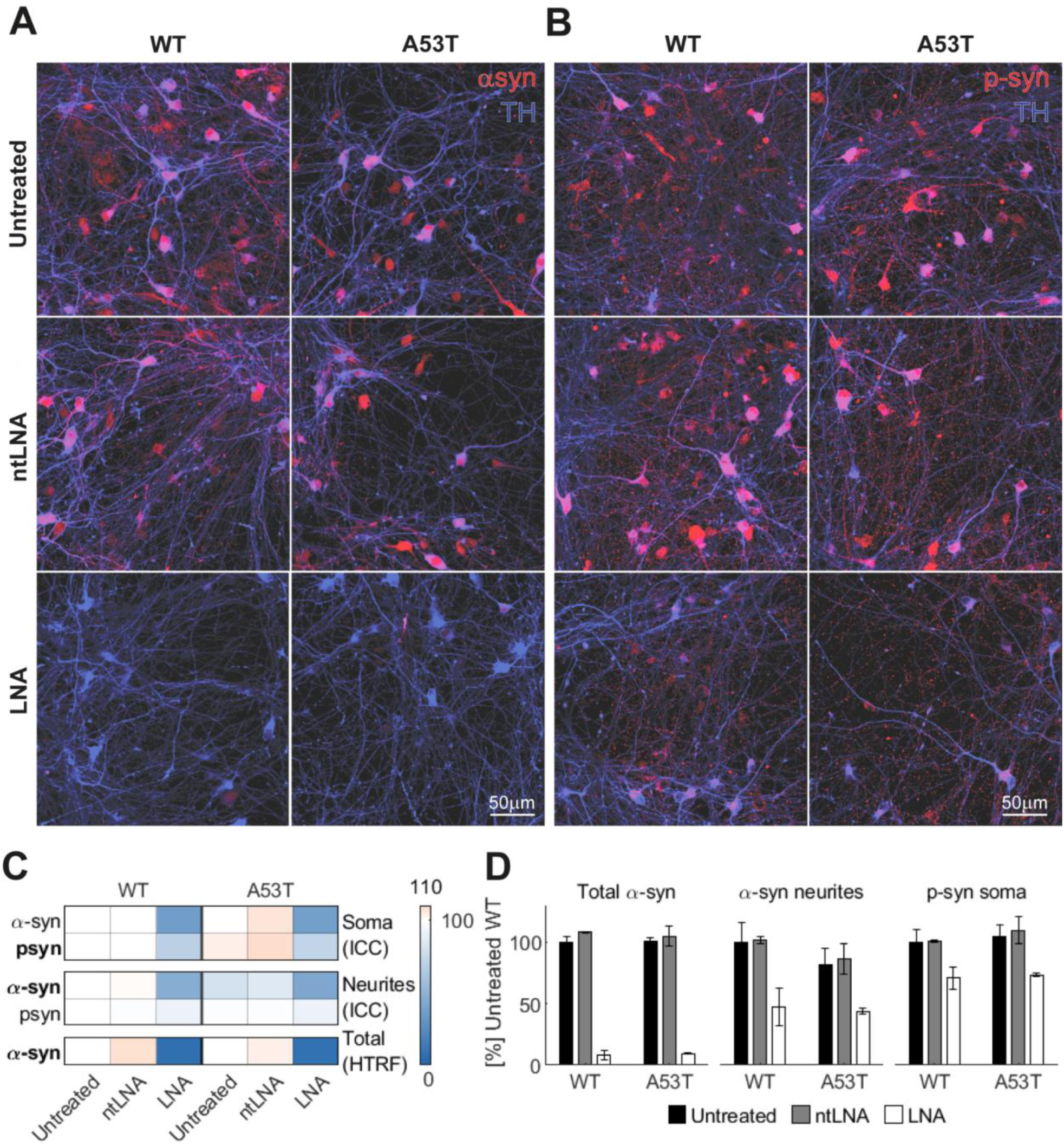
Effective downregulation of α-synuclein expression through LNA administration. **(A)** Immunocytochemical (ICC) stainings for α-syn expression in WT/A53T untreated and non-targeted LNA (ntLNA)/SNCA-targeting LNA-treated cultures at DIV21 (N=3 per condition). Both genotypes displayed similar patterns in their α-syn expression (α-syn: red, TH: blue), which was most prominent in the soma, but also clearly visible in neurites (left panels). ntLNA-treatment did not result in any visible alterations of α-syn levels (middle panels), while LNA-administration resulted in an almost complete absence of the α-syn signal (lower panels). **(B)** Phosphorylated α-synuclein (p-syn) expression displayed a similar pattern across the different conditions (p-syn: red, TH: blue). **(C)** Summary table of changes in somatic and neuritic α-syn, p-syn and total α-syn (quantified by a Homogeneous Time Resolved Fluorescence (HTRF) assay). α-syn/p-syn levels were quantified automatically from ICC stainings and are color coded in reference to the untreated WT condition. LNA treatment reduced expression levels significantly (all p<0.001, Tukey’s test), confirming the efficacy of the applied LNA to reduce both α-syn and p-syn expression *in vitro*. Importantly, ntLNA treatment did not alter expression levels significantly (all p>0.05, two-factor ANOVA). **(D)** While the LNA reduced total α-syn levels significantly, there was no significant difference between the genotypes (p=0.820, two-factor ANOVA, left panel). Neuritic α-syn levels, however, were significantly decreased in A53T DA neurons (p<0.001, two-factor ANOVA, middle panel), whereas somatic p-syn levels were significantly increased (p=0.001, two-factor ANOVA, right panel).

### Downregulation of α-synuclein strongly alters electrophysiological features and development

Next, we probed the effect of an LNA-induced reduction of α-syn expression on the functional phenotypes of WT/A53T DA neuronal cultures. This analysis was based only on activity-related single-cell and network metrics, i.e., features that had the highest predictive power in distinguishing between both genotypes (see RF accuracies, Figure 3A). Since electrophysiological features of ntLNA-treated cultures resembled results obtained for untreated cultures (see Figure 5 - figure supplement 1, **Supplemental tables 12-13**), we combined untreated and ntLNA-treated cultures for further analyses to increase statistical power. Results indicated that LNA-mediated α-syn downregulation led to distinct developmental trajectories for both genotypes (Figure 5). Network and single-cell activity metrics mostly showed similar trends in both genotypes, and treatment-induced differences were only significant in WT cultures (Figure 5A, see asterisks for significant features, LMM; **Supplemental tables 14-15**). At the network-level, LNA treatment of DA cultures resulted in shorter, but more frequent bursts of smaller peak amplitude (Figure 5B). While the NRF decreased continuously throughout development in WT control cultures, LNA treatment reversed this trend (Figure 5B, upper panel); A53T cultures displayed a similar development (LNA treatment increased the NRF). Similarly, the mean burst duration (MBD) increased consistently for control WT cultures, whereas LNA-treated WT cultures resembled A53T cultures, which displayed a decrease in the MBD after week 2 (Figure 5B, lower panel). Consequently, k-means clustering (k=2) on the PCA results grouped LNA-treated cultures with control A53T cultures, while control WT cultures formed a separate cluster (Figure 5C, lower panel). PCA performed only on network features resulted in an even more pronounced separation and additionally differentiate between control A53T and LNA-treated cultures (Figure 5C, upper panel).

**Figure 5.**
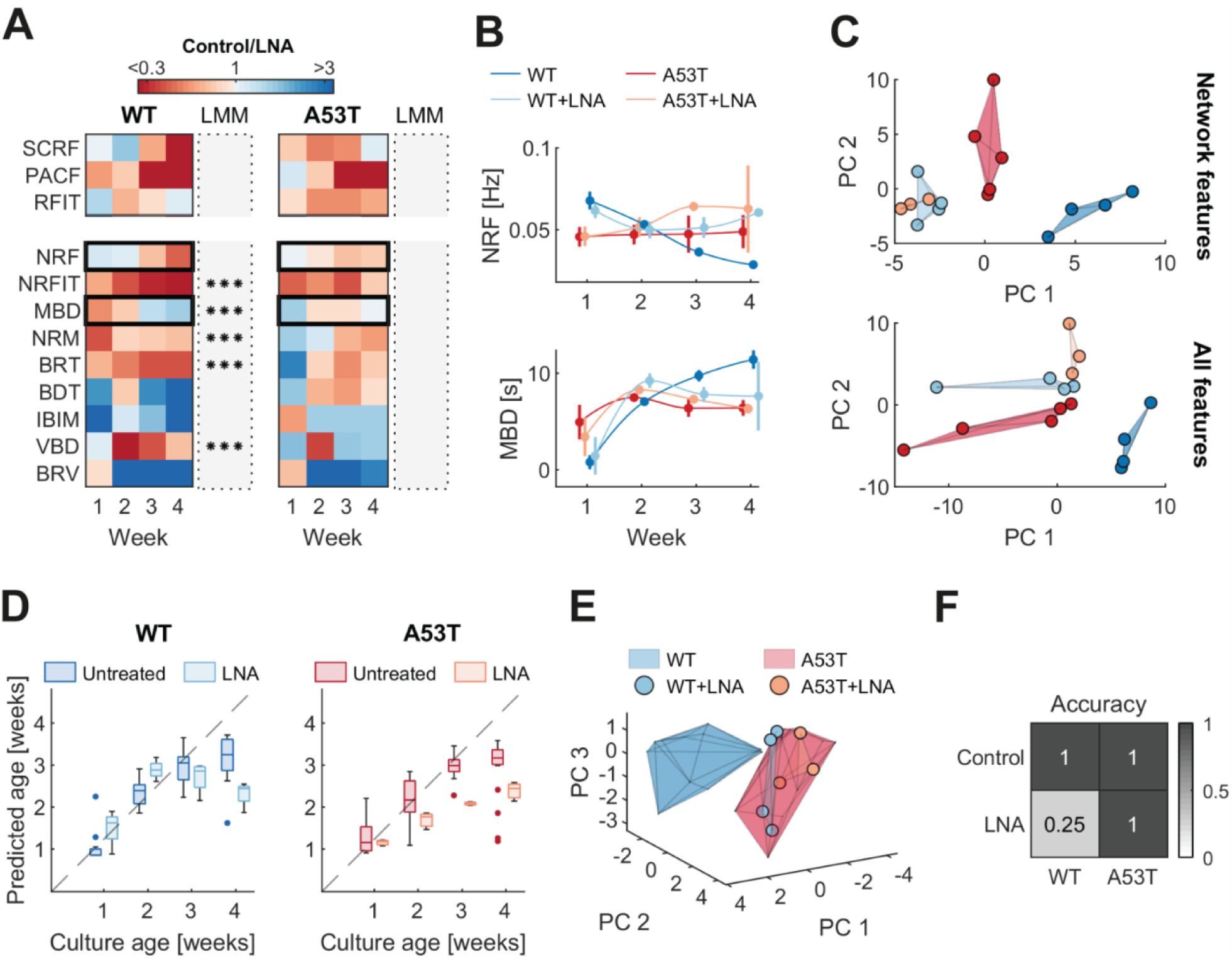
Downregulation of α-synuclein alters electrophysiological phenotypes and culture development. **(A)** Differences in α-syn levels translate into distinct electrophysiological phenotypes. Heatmaps show the relative feature values of activity-based single-cell (top panels) and network features (bottom panels) with the highest classification accuracies (see RF accuracies, Figure 3a). The treatment affected the activity of both genotypes similarly: At week 4, most metrics showed the same relative feature value between treated and control cultures in both genotypes, but only alterations in WT cultures were significant (significance of the LNA-treatment effect), and the treatment-time interaction was estimated by linear mixed-effect models (LMMs); see asterisks, ***p<0.001; **Supplemental tables 14-15**). **(B)** The effect of the LNA treatment on the developmental trajectory was clearly reflected in the network regularity frequency (NRF, upper panel), where LNA-treated cultures diverged from their respective controls at week 3. The mean burst duration (MBD, lower panel) was specifically affected in LNA-treated WT cultures, which followed the developmental trajectory of A53T cultures. **(C)** Visualization of the first two PCs showed a co-localization of treated WT and A53T cultures with control A53T cultures, while WT controls were clearly separated (lower panel). Using the first two PCs of network features led to an additional separation of LNA-treated and control A53T cultures (upper panel). **(D)** RF age regression analysis indicated a clear effect of the LNA treatment on DA culture maturation. Both LNA-treated WT and A53T cultures were estimated to be younger than the respective controls. **(E)** LNA-treated WT cultures became more similar to A53T cultures, as indicated by a clustering analysis including the main dataset (untreated cultures). **(F)** Accuracy matrix of the RF prediction. The use of the previously trained RF classifier yielded perfect classification accuracies for control cultures (N_WT_=4, N_A53T_=5) and LNA-treated A53T cultures (N=3), while all but one of the LNA-treated WT cultures were misclassified (25%, N=4).

Finally, we assessed how α-syn downregulation affected the RF age prediction (Figure 5D). LNA-treated WT cultures were initially estimated to be older than their controls, but after week 3, their age was increasingly underestimated (Figure 5D, left panel). This time point coincided with the divergence of the developmental trajectories for several features (e.g., NRF, Figure 5B, upper panel). An even stronger effect was observed for A53T cultures, where the age of LNA-treated cultures was underestimated throughout the whole development (Figure 5D, right panel). Taken together, these data likely indicate a developmental delay induced by the strong LNA-mediated reduction of α-syn. Figure 5E shows the effect of LNA treatment on functional phenotypes in the PC space: LNA-treated cultures clearly co-localized with control A53T cultures, highlighting the shift towards the mutant phenotype as a result of the α-syn downregulation (week 3 and 4). Similarly, k-means clustering (k=2) grouped all but one LNA-treated culture with the A53T cluster (in red). Correspondingly, LNA-treated cultures were predominantly classified as A53T by the previously trained RF model (Figure 3D, all features), irrespective of their actual genotype (N_WT_=4, accuracy=25%; N_A53T_=3, accuracy=100%), while control cultures of both genotypes were all classified correctly as controls (N_WT_=4, N_A53T_=5).

## Discussion

The primary goal of this study was to investigate the impact of the *SNCA*-A53T mutation on the electrophysiological phenotype of human induced pluripotent stem cell (iPSC)-derived dopaminergic (DA) neurons cultured on high-density microelectrode arrays (HD-MEAs). To this end, we developed *DeePhys*, a new method that combines automated electrophysiological feature extraction from spike sorted HD-MEA recordings with a robust machine learning workflow (Figure 1). The *DeePhys* approach allowed us to infer and quantitatively evaluate functional phenotypic differences between isogenic mutant and control DA neuron lines across development and treatment conditions. We find that both cell lines can be reliably classified according to their electrophysiological phenotypes, with network-level metrics representing the more informative features. Locked nucleic acid (LNA)-mediated downregulation of α-synuclein (α-syn) had specific effects on α-syn and phosphorylated α-syn levels as well as on electrophysiological features and impacted the overall developmental trajectories of both cell lines. We expect that *DeePhys* and the approach presented here has great potential to be applied to a wide range of other human cellular models of neurological diseases *in vitro*.

### Large-scale electrophysiological characterization of dopaminergic neurons

In line with previous studies (Hartfield et al., 2014; Tepper et al., 1990; Watmuff et al., 2012), we found a rapid increase of spontaneous electrical activity in both DA cell lines during development and an emergence of pronounced network bursts after about two weeks *in vitro* (Figure 3). Intrinsic burst firing has been previously described as an important electrical feature of DA neurons in acute slices of neonatal rodents (Dufour et al., 2014; Ferrari et al., 2012), which precedes irregular activity in juvenile animals and mature tonic and phasic activity in later stages of development (Paladini & Roeper, 2014). The burst activity that we observed is also in agreement with previous reports of iPSC-derived DA neuron physiology, studied by single-cell patch-clamping and calcium imaging (Hartfield et al., 2014; Kriks et al., 2011; Renner et al., 2020; Ryan et al., 2013; Zygogianni et al., 2019). HD-MEAs, however, offer distinct advantages over other electrophysiological methods, such as higher throughput and higher temporal resolution, and thus enable a more systematic functional characterization. Our results demonstrate that human iPSC-derived DA neuron-astrocyte co-cultures, maintained over several weeks *in vitro* on HD-MEAs, can be used to systematically study the electrophysiological development of neuronal networks at scale (>130.000 neurons in total). Moreover, we show that spike-sorted HD-MEA network recordings provide a wealth of information for a detailed functional assessment of DA neurons across scales (>30 single-cell and network features).

### The *SNCA*-A53T mutation alters the electrophysiological phenotype of dopaminergic neurons

The A53T point mutation is one of the best studied *SNCA* mutations causing familial Parkinson’s disease (PD) (Polymeropoulos et al., 1997). A key finding of our study is that DA neurons carrying the A53T mutation can be clearly distinguished from isogenic DA neuron cultures using single-cell and network-level electrophysiological features. A distinct A53T phenotype was also previously reported *in vivo*, where age- and brain-region-specific effects of the α-syn mutation on DA neurons of the substantia nigra *pars compacta* (SNc) have been reported in a transgenic A53T mouse model (Gispert et al., 2003). A related study found an increase in SNc DA neuron AP firing in 7-9 months old animals (Subramaniam et al., 2014) and proposed a link between DA neuron hyperactivity, mutant α-syn, and dysfunctional A-type Kv4.3 channels. A hyperactivity phenotype in some SNc DA neurons may precede a reduction in DA neuron firing (Janezic et al., 2013) and the development of altered dopamine release and synaptic plasticity described at later stages of mutant rodent development (Kurz et al., 2010).

Similarly, studies found multi-facetted alterations in patient-derived iPSC DA neurons, including impaired mitochondrial function (Ryan et al., 2013), bioenergetic dysfunctions (Zambon et al., 2019), perturbations in cholesterol metabolism and the proteasomal pathway (Fernandes et al., 2020), and altered expression of genes involved in synaptic signaling and defective synaptic connectivity (Kouroupi et al., 2017). Functional characterization of iPSC-derived A53T DA neurons revealed an increased firing rate compared to control neurons (Zygogianni et al., 2019). In line with this finding, our mapping of DA neuron electrophysiological properties indicated a moderately reduced firing rate in WT DA cultures (Figure 2a). It is noteworthy, however, that low-activity units (<0.1 Hz) were not included in our analyses to assure good feature inference quality and to focus on neurons that can be reliably spike-sorted, which likely biases comparisons of absolute activity levels (Yger et al., 2018).

We found that changes related to the regularity of neuronal activity occurred in several features of both genotypes. A53T cultures displayed an earlier onset of spontaneous synchronized activity and smaller, but more frequent bursts later during development. The early emergence of synchronized population activity among A53T DA neurons was somewhat unexpected, as it would indicate rapid axonal outgrowth and synaptogenesis. However, upregulation of genes involved in neurite growth in A53T DA neurons has been recently reported (Fernandes et al., 2020). Also, the α-syn induced enhanced synaptogenic properties of astrocytes through TGF-β1 signaling could be relevant in explaining the observed early coactivity (Diniz et al., 2019).

The differences between WT and A53T burst dynamics, as observed in our study, could also provide a link to a previous study, which found that mutant α-syn interferes with vesicle recycling and the regulation of the recycling pool (Xu et al., 2016). Both processes are essential in maintaining synaptic activity over prolonged periods of time. Alterations in the functionality of α-syn due to the A53T mutation hence may alter the duration and frequency of network bursts. Additionally, mitochondrial defects and bioenergetic deficits, caused by the A53T mutation, may reduce the length of high-activity periods, as the energy demand of prolonged spiking cannot be met (Ryan et al., 2013; Zambon et al., 2019).

### Effects of reduced α-synuclein levels on spontaneous activity

Reduced levels of α-syn in A53T DA neurons could be another explanation for the observed A53T functional phenotype, as a recent transcriptomic analysis of human iPSC-derived DA neurons revealed that *SNCA* was downregulated by the introduction of the A53T mutation (Fernandes et al., 2020). Indeed, we observed reduced expression of α-syn specifically in the neurites of A53T DA neurons (Figure 3). We also found increased levels of phosphorylated α-syn in the soma of A53T DA neurons (Figure 3d), a post-translational modification that has been implicated in α-syn aggregation and neurodegeneration in patients suffering from synucleinopathies (Fujiwara et al., 2002; Okochi et al., 2000) and animal models of PD (Neumann et al., 2002; Takahashi et al., 2003). This result supports the hypothesis that the accumulation of α-syn sequesters the functional forms of α-syn and reduces its necessary function at the presynaptic terminal (Benskey et al., 2016), where it has been implicated in a variety of regulatory functions. For example, α-syn has been reported to be involved in the maintenance of the synaptic vesicle (SV) pool size through vesicle recycling and inhibition of intersynaptic vesicle-trafficking (Nemani et al., 2010; Scott & Roy, 2012; Sun et al., 2019). In primary hippocampal neurons, α-syn downregulation revealed a significant reduction in the distal pool of synaptic vesicles (Murphy et al., 2000). Additionally, α-syn clusters synaptic vesicles and attenuates the release of neurotransmitters (Wang et al., 2014). A reduction in α-syn levels may, therefore, restrict the SV pool size before bursts and the recycling rate during network bursts - which could result in shorter, low-frequency bursts. A similar effect was previously reported *in vivo*, where a lack of α-syn attenuated synaptic responses to prolonged repetitive stimulation as a consequence of SV depletion (Cabin et al., 2002).

The LNA-mediated downregulation of α-syn provided further support for this hypothesis. The strongly decreased α-syn levels in DA neurons after the LNA treatment (Figure 4) resulted in an electrophysiological phenotype that resembled the phenotype of A53T mutant cultures (Figure 5). This finding was also confirmed by applying a pre-trained RF model on LNA-treated cultures, which classified all but one LNA-treated culture as A53T (Figure 5f). However, most phenotypical features - such as an increased burst frequency and a reduced burst duration – were even more pronounced in the LNA condition, which might relate to the respective α-syn levels. This finding proved true regardless of the genotype, as WT and A53T cultures were affected similarly by the LNA-induced reduction in α-syn levels. Results from our RF age prediction further supports the notion that α-syn downregulation likely disrupts physiological neuron function and maturation (Figure 5d), as the age of LNA-treated cultures was systematically underestimated. This effect was more pronounced in A53T cultures than in WT cultures, which may be indicative of an additive process in A53T mutant cultures.

### Limitations of the study

Despite the great promises and attractiveness to use human-derived cellular models of neurological diseases *in vitro*, there are important caveats that need to be taken into account: First, we observed considerable inter-batch variability with regard to the overall activity but also other features. This was specifically evident when comparing the main data set (batch 1 and 2) to that of the LNA-treatment experiment (batch 3). While we would assign a large part of the variation to the different medium change protocols that were used for normal culturing and applying the LNA, future studies will need to more systematically investigate the effect of medium composition and medium exchange protocols on the development of DA cultures (Bardy et al., 2015).

Another important consideration is the use of healthy WT astrocytes in our HD-MEA co-culture system. While the use of healthy WT astrocytes allowed us to attribute phenotypic changes to alterations in neurons rather than astrocytes, it does not reflect the real *in vivo* pathological condition, where astrocytes also carry the mutation and potentially contribute to the disease. Recent *in vitro* studies have emphasized the importance of astrocyte-neuron interactions in the pathogenesis of PD (di Domenico et al., 2019). Lastly, it is important to note that human iPSC-derived cells, as generated with todays’ iPSC technology, can only partially represent the physiology of matured neurons and astrocytes (Cornacchia & Studer, 2017).

Finally, despite the reported maturational changes, human iPSC-derived DA neurons of the current study showed relatively small AP amplitudes, which rendered spike sorting challenging and potentially affected both waveform and activity features of the resulting templates. Considering this observation and the important aging component of PD pathogenesis, future studies should look into strategies to induce cellular aging in DA neurons and study their physiology or functional phenotypes at later stages of development.

### Advantages of the *DeePhys* workflow

Due to the commercial availability of a wide range of human iPSC-derived neuronal cell lines, genetical engineering, and HD-MEA recording technology, there is increasing demand for easy-to-use analytical tools for researchers coming from outside the electrophysiological domain. While spike sorting has become more accessible (Buccino et al., 2020; Magland et al., 2020), subsequent analyses often require extensive domain/data analysis knowledge. Here, we introduced *DeePhys*, which provides an intuitive analysis pipeline that extracts multi-parametric information from spike-sorted recordings in a largely customizable and easily interpretable way and supplements analyses with appropriate visualizations. *DeePhys* is easily scalable, requires only minimal manual intervention, and can be used as a screening tool to investigate different human cellular models of neurological diseases *in vitro*. Applying *DeePhys* to genetically modified and/or patient-derived neurons will facilitate the discovery of specific electrophysiological biomarkers, phenotype-driven evaluation of new treatment approaches, and more personalized treatments for PD (Trudler et al., 2021; Valadez-Barba et al., 2020).

### Conclusion

In sum, the application of *DeePhys* on iPSC-derived DA neurons suggests that α-syn is essential for the regulation of sustained synaptic activity in DA neurons, which may link to previous findings regarding the role of α-syn in the maintenance of the SV pool (Cabin et al., 2002; Murphy et al., 2000; Scott & Roy, 2012; Sun et al., 2019; Wang et al., 2014). Cultures of neurons carrying the A53T mutation showed altered single-cell waveforms as well as altered network development and dynamics with respect to the isogenic healthy cultures. A53T cultures displayed an earlier onset of bursting, yet shorter and more frequent bursts throughout development. Burst-related metrics were the most reliable predictors to differentiate between the WT and A53T genotypes, which evidences their potential as biomarkers. LNA-treatment resulted in alterations in network dynamics and single-cell activity that resembled those of the A53T phenotype, which might be correlated to the respective levels of functional α-syn. Age regression analysis indicated a developmental delay in the LNA-treated cultures, which highlights the importance of α-syn for physiological DA network development. Our results show that *DeePhys* provides an easy-to-use, scalable, quantitative analysis platform for functional phenotype screening and for addressing important biomedical questions in the development of potential treatments.

### Resource Availability

#### Lead contact

Further information and requests for resources and data should be directed to and will be fulfilled by the Lead Contact, Philipp Hornauer.

#### Data and Code Availability

The code of *DeePhys* and for the figures used in this manuscript is available at: https://github.com/hornauerp/EphysDopa.git.

The raw data sets supporting the current study have not been deposited in a public repository due to the excessive file size (> 3TB) but are available from the corresponding author upon reasonable request. The processed data set containing the extracted features is available on the git repository.

## Materials and Methods

### High-density microelectrode array recordings

Electrophysiological recordings were obtained using the CMOS-based high-density microelectrode arrays (HD-MEA) of the type “MaxOne” by MaxWell Biosystems (MaxWell Biosystems, Zurich, Switzerland). This HD-MEA features a total of 26’400 electrodes in a 120 × 220 electrode grid with a microelectrode center-to-center spacing of 17.5 μm, an overall sensing area of 3.85 × 2.10 mm^2^, and allows for simultaneous recordings from up to 1024 electrodes at a sampling rate of 20 kHz (Müller et al., 2015). Recordings were performed inside an incubator at 36°C and 5% CO_2_ using the MaxLab Live recording software (MaxWell Biosystems). Recordings started at day *in vitro* (DIV) 7 and were subsequently performed once a week over the course of 5 weeks. Each recording consisted of an activity scan to determine the electrode selection and a subsequent network recording. The activity scan consisted of 7 sparse electrode configurations (center-to-center spacing: 35 μm, every 2^nd^ electrode), that were recorded 2 minutes each. To fully capture network dynamics, electrodes displaying the highest firing rate were selected, and spontaneous network activity was recorded for 15 minutes.

### Cell culture and plating

#### Cell lines

Human iPSC-derived DA neurons carrying a heterozygous A53T mutation (cat. C1112, iCell DopaNeurons A53T), an isogenic control line (cat. C1087, iCell DopaNeurons) and astrocytes (cat. R1092, iCell Astrocytes) were purchased from FUJIFILM Cellular Dynamics International (FCDI, Madison, WI, United States). The A53T cell line was generated through nuclease-mediated single-nucleotide polymorphism alterations of the isogenic control line. Midbrain DA neuron differentiation from iPSCs was based on a protocol from the Lorenz Studer lab (Kriks et al., 2011). The vendor guarantees a purity of at least 70% for midbrain DA neurons and 95% for astrocytes.

#### Cell plating

The protocol used for cell plating was previously established in (Ronchi et al., 2021). Prior to cell plating, HD-MEAs were sterilized in 70% ethanol for 30 minutes and rinsed 3 times with sterile deionized (DI) water. To enhance cell adhesion, the electrode area was covered with 20 μL of 0.05 mg/mL poly-L-ornithine (PLO) solution (cat. A-004-C, Sigma-Aldrich, Saint Louis, MO, United States) and incubated at 37°C for 2 hours. Next, the PLO solution was aspirated, and the HD-MEA was rinsed 3 times with sterile DI water. Next, we added 10 μL of 80 μg/ml laminin (cat. L2020-1MG, Sigma-Aldrich) in plating medium (see below) directly on the electrode area and incubated the chips at 37°C for 30 minutes. The plating medium consisted of 95 mL of BrainPhys Neuronal Medium (cat. 05790, STEMCELL Technologies, Vancouver, Canada), 2 mL of iCell Neural Supplement B (cat. M1029, FCDI), 1 mL iCell Nervous System Supplement (cat. M1031, FCDI), 1 mL N-2 Supplement (100X, cat. 17502048, Gibco), 100 μL laminin (1 mg/mL, cat. L2020-1MG, Sigma-Aldrich) and 1 mL Penicillin-Streptomycin (100X, cat. 15140122, Gibco). In the meantime, the cryovials containing the DA neurons and astrocytes were thawed in a 37°C water bath for 3 minutes. The cells were then transferred to 50 mL centrifuge tubes, and 8 mL plating medium (at room temperature) were drop-wise added (numbers are indicated for 20 chips). Cell suspensions were centrifuged at 380 x g (1600 RPM) for 5 minutes, and the supernatant was aspirated. Cell pellets were then resuspended in plating medium and combined to achieve a final concentration of 10’000 DA neurons and 2000 astrocytes per μL. Finally, 100’000 DA neurons and 20’000 astrocytes were seeded on each HD-MEA by adding 10 μL of the prepared solution directly to the laminin droplet. Next, chips were incubated for 1 hour, and 1.5 mL of plating medium were carefully added. Chips were equipped with a lid, placed inside a 100 mm petri dish - to facilitate transport and to reduce the risk of contamination - and kept inside an incubator at 37°C and 5% CO_2_. Additionally, a 35-mm petri dish, filled with DI water, was placed inside the larger petri dish to counteract evaporation. One day after the plating, we replaced 50% of the medium and resumed the normal media change protocol (one third of the medium was exchanged twice a week). Cultures were allowed to equilibrate for 3 days after the medium change to prevent effects on the recordings.

### Data analysis

#### Dataset

The dataset for the genotype comparison consisted of 18 WT and 19 A53T cultures, pooled across two batches with identical cell culture and recording protocols. No statistical method was used to predetermine sample size. Only cultures with recordings from all recording time points were included in the classification analysis, which reduced the number of available cultures for the classification analysis to 14 WT and 15 A53T cultures. Recordings were excluded if the culture detached from the HD-MEA, or if the HD-MEA displayed severe malfunctions. The data set of the LNA-treatment consisted of 10 WT and 10 A53T cultures for the statistical analysis, and of 8 WT and 8 A53T cultures for the classification analysis. All steps of the data analysis were performed on the high-performance computing cluster “Euler” of ETH Zurich. Feature extraction and statistical analysis were performed using custom-written code in MATLAB 2021a.

Linear mixed-effects models of the form *Y* ∼ 1 + *Genotype* \**Time*+(1|*Subject*) were applied to compute statistical significance, using the restricted maximum likelihood (REML) estimation as a fitting method. The resulting *p*-values were then adjusted for the number of comparisons (Bonferroni correction). Feature values were considered outliers and excluded if they exceeded three scaled median absolute deviations. A fraction of the data set used here originated from a previous study (Ronchi et al., 2021), which, however, did not include a more systematic analysis of waveform/network features of spike-sorted units.

#### Spike sorting

In order to systematically characterize single-cell and network features of the recorded DA neurons, we spike-sorted HD-MEA recordings. This step was necessary, as the low electrode pitch of the used HD-MEA increases the likelihood that electrodes pick up signals of multiple neurons or neuronal processes. Spike-sorting was performed using SpyKING Circus 0.8.5 (Yger et al., 2018). Spike-sorting parameters were tailored to the data set at hand and are provided in the Supplemental Material (**Supplemental table 5**). Spike sorting results underwent further quality control: only units that had a firing rate of at least 0.1 Hz and few refractory-period violations (below 2%) were included in further analyses.

#### Feature extraction

In the following, we provide an overview of the features that were used to characterize the development and functional state of the neuronal cultures, many of which have been used previously for similar tasks (Eisenman et al., 2015; Farkhooi et al., 2009; Jia et al., 2019; Stratton et al., 2012; Weir et al., 2014). These features were inferred from the *electrical footprints* (EFs) and the respective *reference waveforms* of spike-sorted units (Figure 2). The *reference waveform* of an EF was extracted from the electrode that recorded the EF maximum waveform amplitude.

##### Single-cell features

Single-cell features here refer to features that can be extracted from a single spike-sorted unit and do not include information from other units of the culture. This group of features can be further subdivided into features that are derived from the reference waveform of the unit (*waveform features*, Figure 2C-D) and features that are derived from the distribution and regularity of spiking activity of the unit (*activity-based features*). Single-cell features were extracted from all units of a culture individually and then averaged to obtain one representative value for the whole culture.

*Waveform features:*

1. *Amplitude (AMPL)* is the maximum absolute value 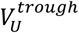 of the reference waveform of one unit *U* (Weir et al., 2014):

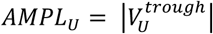
2. *Half width (HLFW)* was defined as the width of the trough at half the trough amplitude value 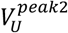 of one unit *U* (Weir et al., 2014).
3. *Asymmetry (ASYM)* was defined as the ratio of the difference and the sum of the peaks after 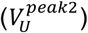 and before 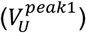 the trough of one unit *U* (Weir et al., 2014):

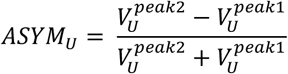
4. *Trough-to-peak ratio (T2PR)* was defined as the absolute value of the ratio of the trough 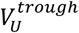 and the second peak 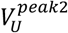 of one unit *U (Jia et al*., *2019):*

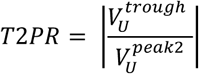
5. *Trough-to-peak delay (T2PD)* was defined as the time difference between the occurrence of the trough 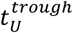 and the second peak 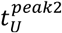 of one unit *U* (Weir et al., 2014):

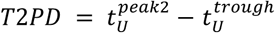
6. *Peak area under the curve (AUCP)* was defined as the integral of the waveform *WF*_*u*_ between the zero crossings before 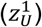 and after 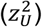 the respective peak 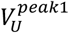 or 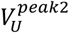 of one unit *U*:

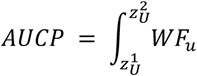
7. *Trough area under the curve (AUCT)* was defined as the integral of the waveform *WF*_*u*_ between the zero crossings before 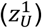 and after 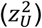 the trough 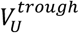 of one unit *U*:

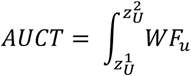
8. *Rise (RISE)* was defined as the slew rate from 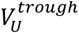 to 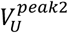 (10^th^ to 90^th^ percentile) of one unit *U* (Jia et al., 2019):

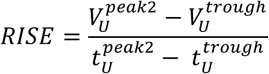
9. *Decay (DECAY)* was defined as the slew rate from 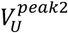 to the resting potential 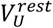 (90^th^ to 10^th^) of one unit *U* (Jia et al., 2019):

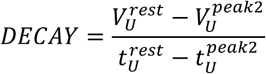

*Activity-based features:*
10. *Mean interspike interval (ISIM)* was defined as the average time between spiking events 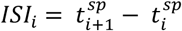 of one unit *U* over a defined number of spikes *N* (Weir et al., 2014):

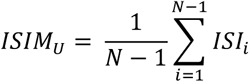
11. *Interspike interval variance (ISIV)* was defined as the variance of interspike intervals *ISI*_*i*_ of one unit *U* over a defined number *N* of ISIs (Weir et al., 2014):

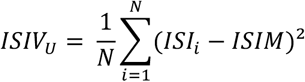
12. *Interspike interval coefficient of variation (ISICV)* was defined as the ratio of the standard deviation to the mean of the interspike intervals *ISI*_*i*_ of one unit *U* (Weir et al., 2014):

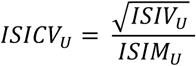
13. *Partial autocorrelation function (PACF)* was defined as the partial autocorrelation of lag 1 for all *ISI*_*t*_ of one unit *U* (Farkhooi et al., 2009):

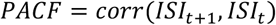
14. *Single-cell regularity frequency (SCRF)* was defined as the frequency with the highest magnitude of the Fourier-transformed activity of one unit *Act*_*U*_:

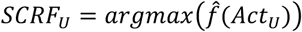
15. *Single-cell regularity magnitude (SCRM)* was defined as the magnitude of the peak frequency of the Fourier-transformed activity of one unit *Act*_*U*_:

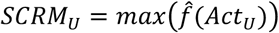
16. *Resonance fit (RFIT)* was defined as the exponential decay constant *d* of the fit through log10-transformed magnitudes *Y* of the regularity frequency harmonics *X* of one unit *U*. The exponential model to fit is of the form:

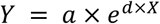

##### Network features

Synchronized spontaneous population activity represents a prominent feature of *in vitro* developing neuronal networks (Blankenship & Feller, 2010), and has been linked to essential neuronal functions, such as central pattern generation (Marder & Bucher, 2001) and information encoding (Kepecs & Lisman, 2003). Network burst patterns were reported to be remarkably variable in primary cortical cultures (Wagenaar et al., 2006). Additionally, compensatory mechanisms appear to be in play that ensure a high degree of robustness of this network behavior (Blankenship & Feller, 2010). As a result, characterizing synchronous network activity represents a promising approach to differentiate between phenotypes. To quantify these bursting dynamics, we performed algorithmic burst detection using the ISI_N_-threshold burst detector (Bakkum et al., 2013). Since we tracked cultures during development, the threshold was set in reference to the overall activity (number of spikes) of the culture (N = 0.1% × *N*_*spikes*_, ISI_N = 1.5s).

1. *Mean interburst interval (IBIM)* was defined as the average time from the end of one burst to the beginning of the next burst 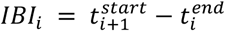 across all *N* bursts of one recording (Weir et al., 2015):

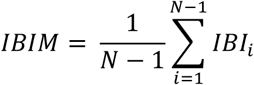
2. *Interburst interval variance (IBIV)* was defined as the variance across all *N* IBIs of one recording:

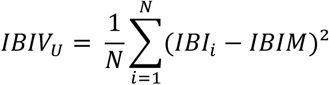
3. *Mean burst duration (MBD)* was defined as the average time from beginning 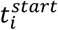 to the end 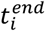 of a burst across all *N* bursts of one recording (Weir et al., 2014):

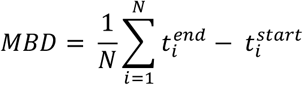
4. *Burst duration variance (VBD) was* defined as the variance across all *N* BDs of one recording:

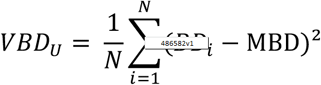
5. *Intra-burst firing rate (INTRABF)* was defined as the number of spikes 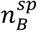 during the total bursting time *T*_*B*_ of one recording (Weir et al., 2014):

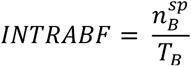
6. *Inter-burst firing rate (INTERBF)* was defined as the number of spikes 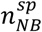 during the total non-bursting time *T*_*NB*_ of one recording:

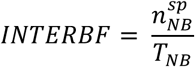
7. *Burst rise time (BRT)* was defined as the average time from the beginning 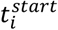 to the peak 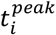 of a burst across all *N* bursts of a recording:

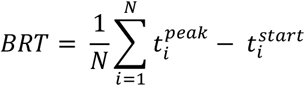
8. *Burst rise velocity (BRV)* was defined as the average slew rate from the coactivity at the beginning 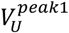 to the coactivity at the peak 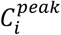 of a burst (10^th^ to 90^th^ percentile) across all *N* bursts of a recording:

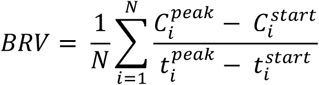
9. *Burst decay time (BDT)* was defined as the average time from the peak 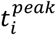 to the end 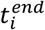 of a burst across all *N* bursts of a recording:

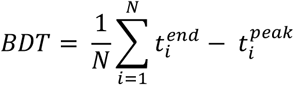
10. *Burst decay velocity (BDV)* was defined as the average slew rate from the coactivity at the peak 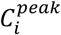 to the coactivity at the end 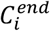 of a burst (90^th^ to 10^th^ percentile) across all *N* bursts of a recording:

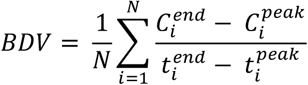
11. *Synchronicity (SYNC)* was defined as the averaged cross-correlation *CC* within a 10 ms interval of all units *U* of a recording, normalized so that the autocorrelations at zero lag equal 1:

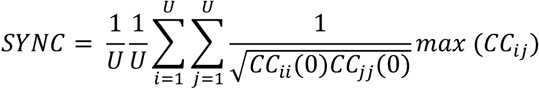
12. *Network regularity frequency (NRF)* was defined as the frequency with the highest magnitude of the Fourier-transformed activity of the whole network *Act*_*NW*_:

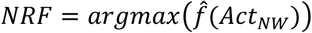
13. *Regularity magnitude (NRM)* was defined as the magnitude of the peak frequency of the Fourier-transformed activity of the whole network *Act*_*NW*_:

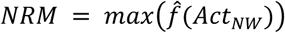
14. *Network resonance fit (NRFIT)* was defined as the exponential decay constant *d* of the fit through log10-transformed magnitudes of the regularity frequency harmonics of the whole network. The exponential model to fit was of the form:

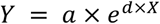

*Phenotype classification and age prediction*

##### Clustering

Input for phenotype clustering consisted of all features from each recording time point. Data was batch-wise transformed into z-scores to minimize inter-batch variability, and z-scores were then used to perform principal component analysis (PCA). Principal components were included until 95% of the variance was explained. K-means clustering was performed to divide the data points into two clusters (k=2), representing the two genotypes (WT/A53T). Cluster centroid positions were initialized 100 times using the k-means++ algorithm, and the solution with the lowest within-cluster sums of point-to-centroid distances was reported (squared Euclidean distance). The phenotype labels were kept hidden during clustering. Then, true labels were compared with the clustering assignment, and the clustering purity was computed as: 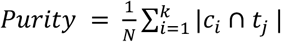,where N is the number of data points, k is the number of clusters, c_i_ is a cluster, and t_j_ is the classification with the maximum count for cluster c_i_.

##### Classification

Training data for classification was batch-wise transformed into z-scores to minimize inter-batch variability, and test data was transformed accordingly, using parameters derived only from the training data. Random Forest (RF) was selected for the analysis, as it provides the option to infer feature importance. Other common classifiers (support-vector machine, Naive Bayes, k-nearest neighbor) were also run as benchmarks (see **Figure 3 - figure supplement 1**). Classification analysis of WT and PD cultures was performed using RF classifiers trained on different feature classes (single-cell, network, combined). Hyperparameter optimization was performed on the training set using 5-fold nested cross-validation (CV) and 100 iterations of Bayesian optimization (for details see **Supplemental table 16**). Due to the high accuracy of the classification, we validated the results using a stratified 5-fold CV to obtain overall accuracies for different features despite the relatively low sample size (N=29). For that, the original data set was partitioned into 5 subsets of equal size, each containing approximately the same number of observations from both classes. One of those subsets was then retained to test the model that was trained on the remaining four subsets. This CV was repeated five times to ensure that every subset was used exactly once as validation data. The generalizability of this classification analysis was probed by predicting the genotype of cultures from another batch with a different medium change protocol (see Methods 4). Furthermore, we assessed the impact of the treatment on the genotype classification by using a model trained on untreated cultures of both genotypes.

##### Age prediction

The prediction of culture age was performed by training RF regression models on features from individual time points. As this task proved to be more difficult, we decided to maximize the number of samples in the training set and used leave-one-out CV to calculate prediction accuracies. The assessment of the differential impact of treatment on development of waveform and activity metrics was performed using RF regression models trained on all untreated cultures of the same genotype.

##### Subsampling

The robustness of the features/models was assessed by performing spatial subsampling on the spike-sorted data by randomly selecting 16, 32, 64, 128, 256 or 512 units and inferring the features from this subset. To mitigate the effect of the random selection, we performed each subsampling and subsequent analysis 25 times.

### Locked nucleic acid experiment

#### Locked nucleic acid-mediated downregulation of α-synuclein expression

We used anti-sense locked nucleic acid (LNA; sequence: TCAGACATCAACCAC) to reduce the α-synuclein expression and a non-targeted LNA (ntLNA) as control (sequence: AACACGTCTATACGC). Antisense LNA GapmeR (cat. 339511) were purchased from QIAGEN (Qiagen N.V., Venlo, Netherlands). From DIV 1 on, LNA or ntLNA were added to the medium to achieve a final concentration of 100 nM. The whole medium was exchanged twice a week to prevent accumulation of LNA in the medium and ensure reliable concentration.

#### Homogeneous Time Resolved Fluorescence analysis of α-synuclein levels

At DIV 21, the medium was aspirated, and plates were washed once with DPBS. The Homogeneous Time Resolved Fluorescence assay (Total alpha Synuclein cellular assay, cat. 6FNSYPEG, Cisbio Bioassays, Codolet, France) was performed according to the manufacturer’s instructions, and fluorescence emission at the acceptor (665nm) and donor wavelength (620nm) were measured in a microplate reader (PHERAstar FSX, BMG LABTECH, Ortenberg, Germany). Total protein concentration was determined using the Pierce™ BCA Protein Assay Kit (cat. 23225, ThermoFisher). The ratios of acceptor and donor emission signals were calculated for each individual well and normalized by the total protein concentration. For each condition, three biological and three technical replicates were measured.

### Immunocytochemistry and microscopy

#### Immunocytochemistry

Cells were fixed using 8% paraformaldehyde solution (cat. 15714S, Electron Microscopy Sciences) and blocked for 1 hour at room temperature (RT) in a blocking buffer containing 10% normal donkey serum (NDS) (cat. 017-000-001, Jackson ImmunoResearch, West Grove, USA), 1% bovine serum albumin (BSA) (cat. 05482, Sigma-Aldrich), and 0.2% Triton X (cat. 93443, Sigma-Aldrich) in PBS (cat. AM9625, ThermoFisher Scientific). Primary antibodies were diluted in blocking buffer (**Supplemental table 17**) and incubated overnight at 4°C. Samples were washed three times with 1% BSA in PBS and incubated with the secondary antibody diluted in blocking buffer for 1 hour at RT (**Supplemental table 17**). After three additional washes with PBS, DAPI was added for 2 min at RT (1:10000).

#### Image quantifications

Images were acquired using the Opera Phenix Plus High-Content Screening System (cat. HH14001000, PerkinElmer, Waltham, MA, USA), and the associated Harmony analysis software was used for quantification. Samples were analyzed by imaging 6 evenly spaced fields, each consisting of 3×3 images, resulting in 54 total images per sample (40x magnification). Images were acquired as z-stacks, flat-field corrected, and converted to a 2D image using maximum intensity projection. Somatic quantification of α-synuclein and phospho-α-synuclein was performed by finding TH^+^ (avg. intensity > 50) nuclei (DAPI mask) and averaging the intensity of the target channel (α-synuclein or phospho-α-synuclein) in the selected area. For neurite quantification, we applied the CSIRO Neurite Analysis 2 algorithm after selecting TH+ nuclei and averaged the intensity of the target channel in the detected neurites.

Statistical analysis was performed in GraphPad Prism 8 using an ordinary two-way ANOVA (factors: genotype and treatment) and Tukey’s test to compare all pairs of means, which accounts for multiple comparisons. Values were again considered to be outliers if they exceeded 3 median absolute deviations.

## Supporting information

Full Supplement

## Acknowledgments

This work was supported by the European Research Council Advanced Grant 694829 ‘neuroXscales’ and the corresponding proof-of-concept Grant 875609 “HD-Neu-Screen”, by the two Cantons of Basel through a Personalized Medicine project (PMB-01-18) granted by ETH Zurich, the Innosuisse Project 25933.2 PFLS-LS, the Swiss National Science Foundation under contract 205320_188910 / 1 and a Swiss Data Science Center project grant (C18-10).

## Author Contributions

Conceptualization: PH, MS, AH, MF and SR; Methodology: PH, MS, DR; Investigation: PH, GP, MS, NA, MF and SR; Software: PH, MS; Formal Analysis: PH; Writing - Original Draft: PH; Writing - Review & Editing: PH, MS, DR, AH, KB, CD, TK, RJ, and VT; Funding Acquisition: MS, AH, KB, VT; Resources: AH, VT and RJ; Supervision: MS, AH, VT and KB, Project Administration: MS, AH; Funding Acquisition: MS, AH, VT and KB.

