## Supplementary material for "Downregulating α-synuclein in iPSC-derived dopaminergic neurons mimics electrophysiological phenotype of the A53T mutation": Full Supplement

#### Figure supplement

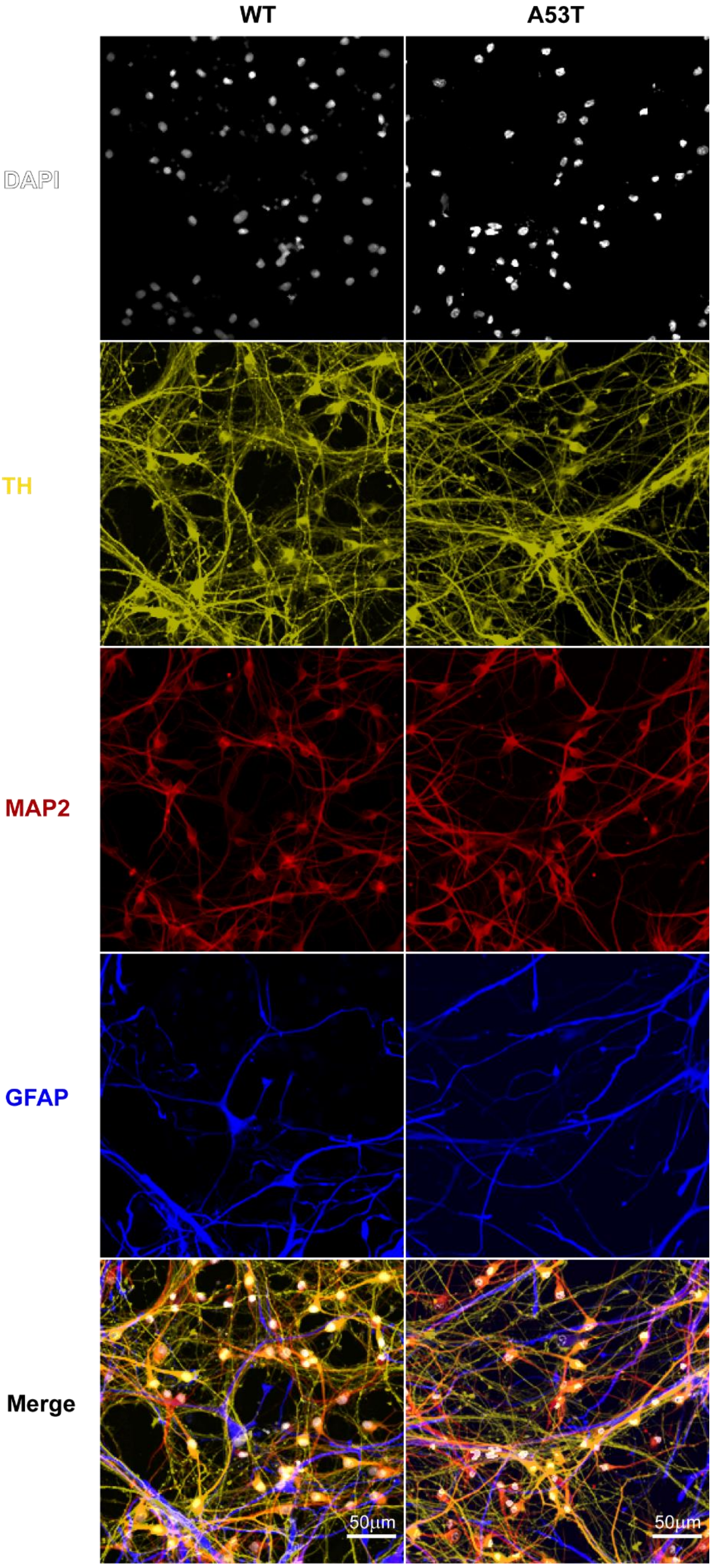

**Figure 2 - figure supplement 1. Similar network formation in WT and A53T neuron-astrocyte co-cultures.** MAP2+ (red)/TH+ (yellow) DA neurons formed dense networks across the whole HD-MEA for both genotypes. Astrocytic processes (GFAP, blue) were also observed to be evenly distributed, indicating a successful integration into the culture.

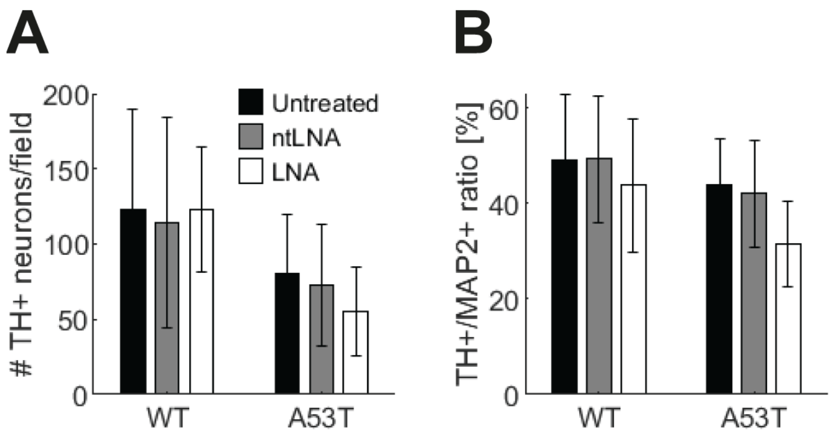

**Figure 2 - figure supplement 2. Reduced number of TH+ neurons in A53T cultures. (A)** Quantification of immunocytochemical stainings revealed a significant decrease of TH+ neurons for the A53T condition (for details see **Supplementary table 1**). **(B)** The ratio of TH+/MAP2+ was also significantly reduced in A53T cultures (for details see **Supplementary table 2**).

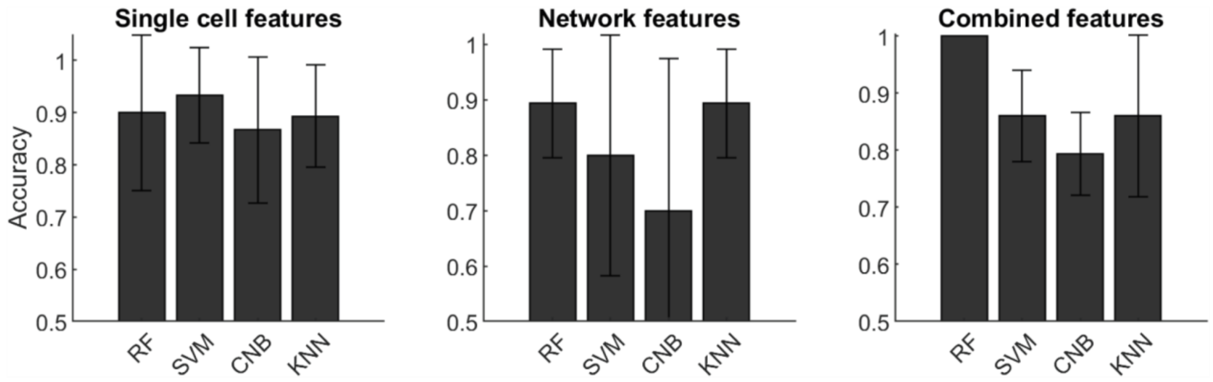

**[Figure 3](#) - figure supplement 1. High genotype classification accuracies across different classifiers.** The comparison of random forest (RF), support vector machine (SVM), Naive Bayes classifier (CNB), and k-nearest neighbor (KNN) showed comparable genotype classification accuracies for RF, SVM, and KNN while CNB performed worse across all feature classes. However, only RF achieved perfect accuracy when both feature classes were combined (right panel).

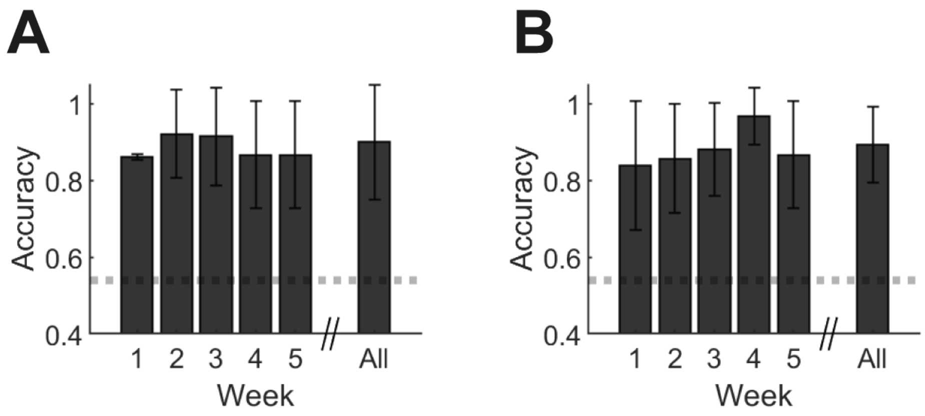

**[Figure 3](#) - figure supplement 2. Complementary classification data.** Probing the difference between both genotypes over time revealed differences between the feature groups: (a) Single-cell features displayed similar accuracy values across all weeks, while (b) network features displayed an increase until a peak value at week 4 with a decline thereafter.

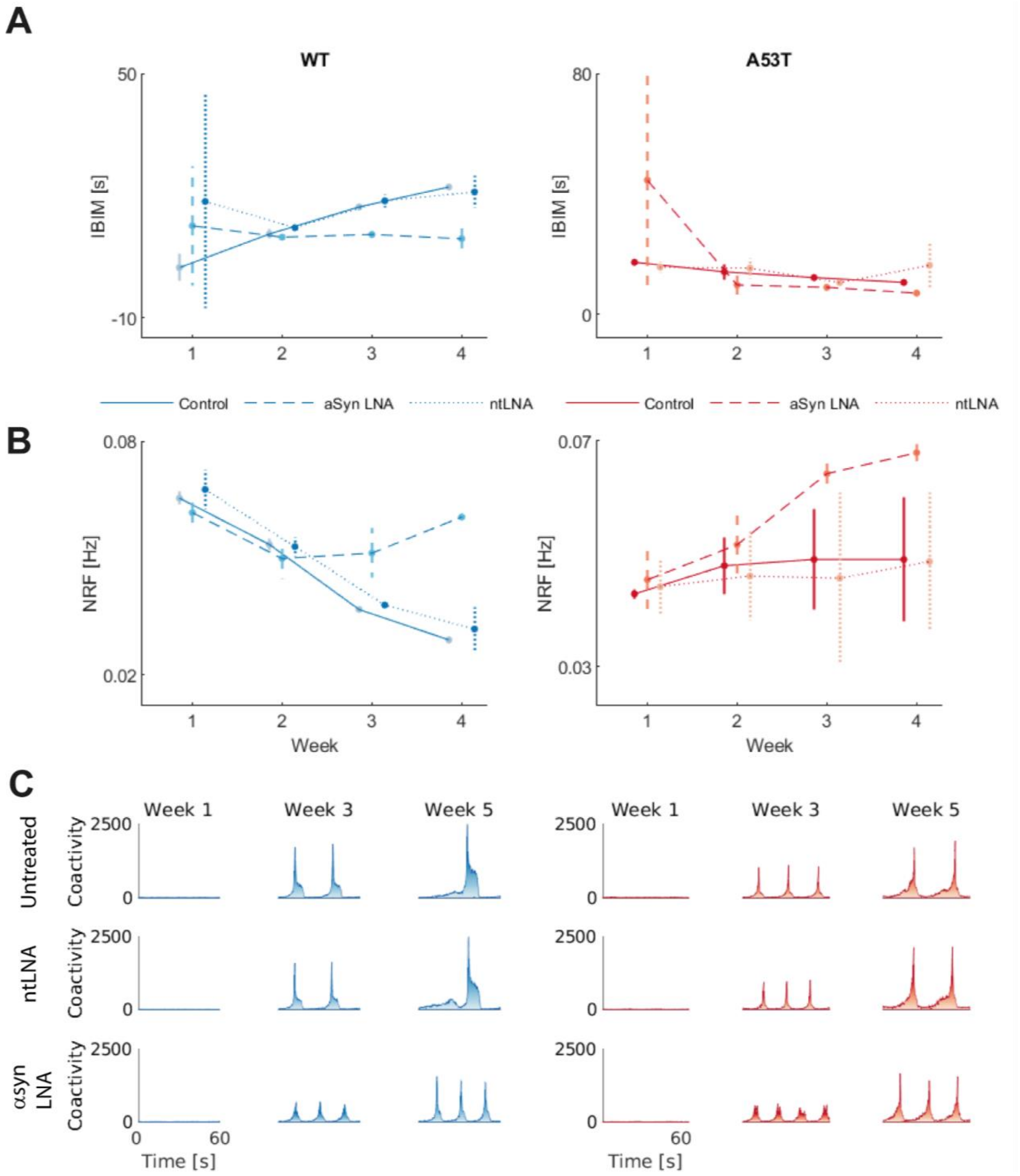

**Figure 5 - figure supplement 1. Similar developmental trajectories of untreated and non-targeted LNA-treated cultures.** Two of the most distinctive features of the two genotypes (mean inter-burst interval (IBIM), panel **A**; network regularity frequency (NRF), panel **B**) displayed similar developmental trajectories in cultures, treated with non-targeted LNA (ntLNA; dotted lines), and untreated control cultures (solid lines). The anti-α-synuclein LNA-treatment (αsynLNA; dashed lines) had a clear effect (i.e., longer interburst interval with higher network regularity frequency). **(C)** The observed network coactivity evidences this effect: While burst shapes and time intervals were very similar in untreated and ntLNA-treated cultures, αsynLNA-treated cultures displayed shorter bursts of smaller amplitude.

### Supplemental tables

| Feature | Time | A53T - Batch 1 | WT - Batch 1 | A53T - Batch 2 | WT - Batch 2 | A53T - Batch 3 | WT - Batch 3 | A53T LNA7 Batch 3 | WT LNA7 Batch 3 | A53T ntLNA Batch 3 | WT ntLNA Batch 3 |
|---|---|---|---|---|---|---|---|---|---|---|---|
| # templates | Week 1 | 927 ± 81 | 942 ± 26 | 897 ± 123 | 871 ± 95 | 1017 ± 77 | 962 ± 8 | 906 ± 44 | 984 ± 59 | 1025 ± 100 | 895 ± 18 |
| | Week 2 | 1104 ± 25 | 1046 ± 59 | 1281 ± 38 | 1035 ± 39 | 1262 ± 36 | 1115 ± 229 | 1289 ± 312 | 1389 ± 95 | 1247 ± 69 | 1203 ± 79 |
| | Week 3 | 1098 ± 61 | 1132 ± 64 | 1272 ± 71 | 1264 ± 130 | 1212 ± 197 | 1268 ± 68 | 1643 ± 8 | 1367 ± 65 | 1149 ± 159 | 1291 ± 6 |
| | Week 4 | 1207 ± 100 | 1129 ± 23 | 1336 ± 68 | 1357 ± 27 | 1451 ± 19 | 1495 ± 36 | 1810 ± 107 | 1193 ± 351 | 993 ± 441 | 1517 ± 41 |
| | Week 5 | 1128 ± 53 | 1051 ± 152 | 1167 ± 243 | 1019 ± 353 | | | | | | |
| # templates after QC | Week 1 | 402 ± 133 | 332 ± 54 | 282 ± 67 | 225 ± 20 | 364 ± 1 | 334 ± 79 | 343 ± 14 | 300 ± 120 | 353 ± 58 | 310 ± 29 |
| | Week 2 | 612 ± 89 | 577 ± 74 | 654 ± 157 | 640 ± 137 | 797 ± 14 | 735 ± 138 | 801 ± 166 | 768 ± 108 | 509 ± 240 | 707 ± 109 |
| | Week 3 | 564 ± 139 | 598 ± 34 | 709 ± 109 | 788 ± 112 | 516 ± 294 | 836 ± 18 | 1241 ± 109 | 557 ± 301 | 244 ± 51 | 641 ± 210 |
| | Week 4 | 644 ± 139 | 624 ± 75 | 564 ± 238 | 574 ± 213 | 924 ± 17 | 1004 ± 49 | 1297 ± 182 | 271 ± 65 | 193 ± 4 | 1047 ± 16 |
| | Week 5 | 609 ± 160 | 455 ± 170 | 352 ± 141 | 389 ± 244 | | | | | | |
| Firing rate [Hz] | Week 1 | 0.56 ± 0.23 | 0.58 ± 0.16 | 0.11 ± 0.02 | 0.15 ± 0.05 | 0.14 ± 0.01 | 0.12 ± 0.08 | 0.12 ± 0.02 | 0.11 ± 0.01 | 0.14 ± 0.02 | 0.095 ± 0.017 |
| | Week 2 | 0.82 ± 0.25 | 0.81 ± 0.24 | 0.36 ± 0.15 | 0.28 ± 0.12 | 0.53 ± 0.22 | 0.50 ± 0.39 | 0.24 ± 0.06 | 0.46 ± 0.07 | 0.70 ± 0.35 | 0.64 ± 0.14 |
| | Week 3 | 1.28 ± 0.27 | 1.33 ± 0.21 | 0.87 ± 0.19 | 0.57 ± 0.22 | 1.12 ± 0.04 | 1.62 ± 0.62 | 0.38 ± 0.19 | 0.82 ± 0.12 | 0.72 ± 0.22 | 1.41 ± 0.41 |
| | Week 4 | 1.35 ± 0.16 | 1.41 ± 0.32 | 0.71 ± 0.38 | 0.46 ± 0.07 | 1.13 ± 0.27 | 1.80 ± 0.02 | 0.57 ± 0.11 | 0.37 ± 0.21 | 0.58 ± 0.59 | 1.55 ± 0.20 |
| | Week 5 | 1.13 ± 0.26 | 1.23 ± 0.44 | 0.62 ± 0.24 | 0.36 ± 0.13 | | | | | | |
| Amplitude [µV] | Week 1 | 35.2 ± 0.9 | 34.0 ± 0.8 | 34.4 ± 0.4 | 33.9 ± 0.6 | 34.5 ± 0.6 | 33.8 ± 1.0 | 33.9 ± 0.1 | 34.4 ± 0.1 | 34.6 ± 0.0 | 34.1 ± 0.0 |
| | Week 2 | 36.5 ± 0.5 | 35.8 ± 0.5 | 36.2 ± 1.2 | 35.6 ± 0.7 | 35.9 ± 0.0 | 35.4 ± 0.5 | 36.1 ± 0.6 | 35.9 ± 0.4 | 35.9 ± 0.9 | 35.6 ± 0.1 |
| | Week 3 | 36.7 ± 0.6 | 36.5 ± 0.4 | 36.7 ± 0.4 | 35.9 ± 0.4 | 35.5 ± 0.8 | 36.6 ± 0.6 | 37.5 ± 0.4 | 36.2 ± 1.6 | 35.8 ± 0.0 | 36.0 ± 0.4 |
| | Week 4 | 37.4 ± 0.7 | 36.9 ± 0.6 | 35.2 ± 1.4 | 35.5 ± 0.5 | 35.7 ± 0.6 | 37.7 ± 0.6 | 38.0 ± 0.0 | 35.9 ± 2.4 | 35.9 ± 0.5 | 36.8 ± 0.4 |
| | Week 5 | 36.8 ± 0.8 | 36.3 ± 0.9 | 34.5 ± 0.7 | 34.4 ± 1.3 | | | | | | |

**Supplemental table 1.** Overview statistics about the datasets of different cell lines across all three experiments.

| Module/Function | Description |
|---|---|
| **Feature extraction** | |
| Class WholeNetwork | Instantiates an object that contains all metadata and network feature values |
| .getMetaData | Uses reference file to import metadata information |
| .getCoordinates | Extracts electrode coordinates |
| .getSpikeTimes | Extracts spike times and sampling rate |
| .getTemplateMatrix | Imports the template matrix resulting from the spike sorting |
| .generateTemplates | Generates Template objects to infer single cell metrics |
| .subsampleNW (optional) | Subsamples Template objects (randomly or most active ones) to infer features from a smaller number of neurons |
| .getActive | Returns Templates that passed the quality control |
| .getRegularity | Calculates network regularity frequency metrics |
| .getBurstTimes | Uses the ISIN burst detection algorithm to detect burst times (Bakkum et al., 2013) |
| .getBurstStatistics | Calculates burst-related metrics from the extracted burst start and end times |
| .getSynchronicity | Calculates synchronicity from normalized cross-correlograms between neurons |
| .getSingleCellFeatures | Infers single cell features from each Template object |
| .saveTMP | Only saves features and metadata to conserve storage space |
| Class Template | Instantiates Template objects that contain all single cell feature values |
| .passCriteria | Check if Template passes quality control |
| .getSpikes | Load single cell spiketimes |
| .calcFrequency | Calculates firing rate |
| .findElectrodes | Finds electrodes on which a Template was detected |
| .calcTemplateArea | Approximates the area of the Template |
| .inferFeatures | Infers all single cell features |
| wave_form_features | Infer waveform features |
| .calcRegularity | Calculate single cell regularity metrics |
| **Data exploration** | |
| Preprocessing | |
| generate_feature_matrices | Constructs matrices with mean network and single cell feature values and their respective standard deviations |
| preparePCAinput | Prepares data for subsequent PCA |
| Visualizations of feature values | |
| plot_features | Plots timelines of mean feature values |
| heatmap_plot | Plots heat map that depicts the relative feature value for two conditions/cell lines |
| plotPCA | PCA plot that color codes different cell |

| | |
|---|---|
| | lines/conditions and depicts cluster outlines in 3D |
| Visualizations of individual recordings | |
| WholeNetwork | |
| .plotWaveforms | Plots an overlay of all Template waveforms |
| .plotCoactivity | Plot aggregated network coactivity |
| .plotNetworkActivity | Plots a 2D map of all Templates and their corresponding firing rate |
| .plotSpikePerUnit | Plots a bar plot displaying the number of spikes per unit |
| .plotActivityRaster | Plots a raster plot for all Templates of the network |
| .plotOverview | Plots all aforementioned subplots together |
| Template | |
| .plotActiveElectrodes | Plots the location of all electrodes, the active electrodes of a Template and its reference electrode |
| .calcSpread(1) | Plots the electrical footprint of a Template with color-coded amplitude values |
| .plotWf | Plots the waveforms of all active electrodes of a Template |
| **Classification analysis** | |
| Processing | |
| filterObjectArray | Filters WholeNetworks by user-specified properties (Age, Mutation etc) |
| classify_genotypes | Main classification function, builds classifier based on user input |
| single_feature_classification | Iteratively builds classifiers for individual features and computes the average accuracy |
| predictAgeFunction | Main age prediction function |
| Visualization | |
| accuracy_barplot | Bar plot for accuracy across several weeks |
| plot_feature_importance | Heatmap with feature importances at individual time points and corresponding accuracy values of single feature classifications |
| predictAgeBoxplot | Plots actual vs predicted age in a box plot |
| **Treatment evaluation** | |
| evaluate_intervention | Classifies a treatment condition based on a pretrained model and evaluates |
| plot_treatment_evaluation | Plots a 2x2 heatmap that displays the accuracy of the evaluate_intervention output for each condition |

**Supplemental table 2.** Modules and functions of the DeePhys pipeline. Functions are only listed once, even if they are reused for other modules.

| Genotype | Treatment | Number of TH+ cells/field | | | | | | | | | | | | Mean ± SD |
|---|---|---|---|---|---|---|---|---|---|---|---|---|---|---|
| WT | Untreated | 37 | 203 | 177 | 48 | 220 | 172 | 73 | 245 | 159 | 75 | 176 | 203 | 122.8 ± 66.8 |
| | | 53 | 164 | 81 | 38 | 79 | 88 | 64 | 168 | 162 | 31 | 170 | 202 | |
| | | 49 | 146 | 206 | 59 | 150 | 223 | 47 | 95 | 158 | 29 | 51 | 121 | |
| | ntLNA | 60 | 112 | 135 | 40 | 145 | 187 | 38 | 163 | 109 | 82 | 162 | 113 | 114.2 ± 39.2 |
| | | 73 | 114 | 102 | 42 | 87 | 115 | 98 | 105 | 86 | 158 | 137 | 113 | |
| | | 146 | 162 | 87 | 204 | 149 | 109 | 134 | 97 | 144 | 87 | 102 | 115 | |
| | LNA | 33 | 72 | 61 | 41 | 255 | 191 | 126 | 161 | 130 | 94 | 165 | 140 | 122.5 ± 69.6 |
| | | 43 | 155 | 148 | 23 | 67 | 45 | 52 | 161 | 177 | 70 | 217 | 234 | |
| | | 55 | 162 | 231 | 37 | 203 | 209 | 54 | 156 | 203 | 25 | 101 | 114 | |
| A53T | Untreated | 51 | 42 | 24 | 101 | 59 | 18 | 113 | 92 | 24 | 100 | 128 | 39 | 80.1 ± 40.6 |
| | | 111 | 71 | 52 | 120 | 94 | 50 | 59 | 45 | 23 | 134 | 47 | 26 | |
| | | 130 | 70 | 42 | 102 | 118 | 72 | 122 | 83 | 103 | 131 | 186 | 100 | |
| | ntLNA | 64 | 32 | 2 | 66 | 59 | 28 | 105 | 62 | 21 | 178 | 90 | 24 | 72.7 ± 41.5 |
| | | 107 | 106 | 25 | 157 | 144 | 58 | 100 | 52 | 10 | 107 | 68 | 19 | |
| | | 82 | 84 | 56 | 77 | 124 | 41 | 108 | 100 | 49 | 96 | 60 | 55 | |
| | LNA | 35 | 53 | 15 | 87 | 50 | 42 | 88 | 115 | 54 | 139 | 60 | 50 | 55.1 ± 29.5 |
| | | 65 | 111 | 54 | 90 | 55 | 28 | 48 | 9 | 25 | 80 | 32 | 58 | |
| | | 51 | 54 | 66 | 87 | 45 | 67 | 43 | 34 | 21 | 17 | 25 | 30 | |

| ANOVA table | SS | DF | MS | F (DFn, DFd) | p value |
|---|---|---|---|---|---|
| Interaction | 7681.0 | 2 | 3840.5 | F (2, 210) = 1.5 | p=0.220 |
| Mutation | 138219.0 | 1 | 138219.0 | F (1, 210) = 54.9 | p<0.001 |
| Treatment | 5886.3 | 2 | 2943.1 | F (2, 210) = 1.17 | p=0.313 |
| Residual | 528962.8 | 210 | 2518.9 | | |

| Tukey's multiple comparisons test | Adjusted p value | Tukey's multiple comparisons test | Adjusted p value |
|---|---|---|---|
| WT:Untreated vs. WT:ntLNA | 0.978 | WT:ntLNA vs. A53T:LNA | <0.001 |
| WT:Untreated vs. WT:LNA | >0.999 | WT:LNA7 vs. A53T:Untreated | 0.005 |
| WT:Untreated vs. A53T:Untreated | 0.005 | WT:LNA vs. A53T:ntLNA | <0.001 |
| WT:Untreated vs. A53T:ntLNA | <0.001 | WT:LNA vs. A53T:LNA | <0.001 |
| WT:Untreated vs. A53T:LNA | <0.001 | A53T:Untreated vs. A53T:ntLNA | 0.989 |
| WT:ntLNA vs. WT:LNA | 0.982 | A53T:Untreated vs. A53T:LNA | 0.286 |
| WT:ntLNA vs. A53T:Untreated | 0.048 | A53T:ntLNA vs. A53T:LNA | 0.673 |
| WT:ntLNA vs. A53T:ntLNA | 0.007 | | |

**Supplemental table 3.** Number of $TH^+$ cells per imaged field. N=6 cultures per condition and 6 fields (3x3 images) per culture.

| Genotype | Treatment | TH+/MAP2+ ratio [%] | | | | | | | | | | | | Mean ± SD |
|---|---|---|---|---|---|---|---|---|---|---|---|---|---|---|
| WT | Untreated | 36.9 | 64.1 | 57.0 | 33.6 | 56.6 | 62.7 | 36.5 | 53.4 | 62.0 | 40.6 | 59.0 | 70.5 | 49.16 ± 13.94 |
| | | 34.8 | 41.3 | 58.9 | 21.3 | 30.7 | 41.2 | 30.1 | 65.9 | 60.8 | 33.8 | 70.1 | 57.9 | |
| | | 41.5 | 70.8 | 57.8 | 42.4 | 64.9 | 65.9 | 41.1 | 51.9 | 40.7 | 32.8 | 45.7 | 33.7 | |
| | ntLNA | 47.8 | 51.2 | 35.2 | 61.2 | 55.0 | 42.8 | 53.9 | 57.4 | 36.0 | 66.7 | 61.1 | 49.3 | 49.42 ± 9.62 |
| | | 57.8 | 42.2 | 58.8 | 50.0 | 47.7 | 43.2 | 39.5 | 56.6 | 59.7 | 30.3 | 58.5 | 63.0 | |
| | | 32.8 | 56.8 | 52.7 | 37.6 | 58.3 | 43.3 | 42.0 | 52.1 | 45.1 | 33.6 | 45.1 | 55.0 | |
| | LNA | 27.2 | 59.0 | 54.1 | 38.7 | 62.9 | 58.5 | 32.9 | 50.8 | 58.0 | 30.1 | 56.5 | 54.9 | 43.81 ± 13.29 |
| | | 34.6 | 49.8 | 56.7 | 23.4 | 34.4 | 40.6 | 22.6 | 41.6 | 34.1 | 30.6 | 68.0 | 58.8 | |
| | | 51.4 | 46.8 | 43.3 | 40.0 | 51.1 | 51.1 | 30.7 | 54.0 | 56.1 | 18.5 | 30.9 | 25.1 | |
| A53T | Untreated | 39.3 | 27.6 | 28.0 | 54.0 | 27.6 | 28.0 | 59.4 | 40.7 | 38.2 | 47.4 | 50.2 | 39.8 | 43.93 ± 11.33 |
| | | 51.9 | 44.4 | 52.3 | 58.0 | 69.4 | 52.9 | 38.9 | 30.7 | 33.8 | 50.2 | 37.3 | 27.3 | |
| | | 51.4 | 41.8 | 26.4 | 53.3 | 61.8 | 38.2 | 56.6 | 40.8 | 38.5 | 59.1 | 45.6 | 40.6 | |
| | ntLNA | 59.9 | 42.6 | 23.3 | 50.0 | 44.2 | 25.0 | 45.8 | 42.9 | 38.6 | 48.7 | 58.2 | 39.0 | 42.10 ± 13.91 |
| | | 54.8 | 56.2 | 38.6 | 41.6 | 47.2 | 42.3 | 33.5 | 28.3 | 4.44 | 35.1 | 39.3 | 23.1 | |
| | | 50.5 | 34.2 | 23.1 | 62.2 | 58.8 | 24.2 | 53.5 | 57.3 | 27.5 | 64.6 | 59.5 | 37.4 | |
| | LNA | 30.4 | 16.1 | 21.6 | 37.9 | 19.2 | 35.4 | 29.8 | 27.6 | 31.7 | 40.5 | 29.6 | 37.0 | 31.65 ± 8.99 |
| | | 32.6 | 25.6 | 17.6 | 15.3 | 24.3 | 22.0 | 31.3 | 28.2 | 21.7 | 40.5 | 33.3 | 34.7 | |
| | | 39.1 | 46.6 | 30.9 | 54.1 | 38.7 | 31.4 | 31.3 | 49.8 | 36.2 | 38.5 | 34.8 | 24.1 | |

| ANOVA table | SS | DF | MS | F (DFn, DFd) | p value |
|---|---|---|---|---|---|
| Interaction | 0.046 | 2 | 0.023 | F (2, 210) = 1.6 | p=0.208 |
| Mutation | 0.366 | 1 | 0.366 | F (1, 210) = 25.4 | p<0.001 |
| Treatment | 0.343 | 2 | 0.171 | F (2, 210) = 11.9 | p<0.001 |
| Residual | 3.031 | 210 | 0.014 | | |

| Tukey's multiple comparisons test | Adjusted p value | Tukey's multiple comparisons test | Adjusted p value |
|---|---|---|---|
| WT:Untreated vs. WT:ntLNA | >0.999 | WT:ntLNA vs. A53T:LNA | <0.001 |
| WT:Untreated vs. WT:LNA | 0.607 | WT:LNA7 vs. A53T:Untreated | >0.999 |
| WT:Untreated vs. A53T:Untreated | 0.643 | WT:LNA vs. A53T:ntLNA | 0.999 |
| WT:Untreated vs. A53T:ntLNA | 0.184 | WT:LNA vs. A53T:LNA | <0.001 |
| WT:Untreated vs. A53T:LNA | <0.001 | A53T:Untreated vs. A53T:ntLNA | 0.999 |
| WT:ntLNA vs. WT:LNA | 0.530 | A53T:Untreated vs. A53T:LNA | <0.001 |
| WT:ntLNA vs. A53T:Untreated | 0.566 | A53T:ntLNA vs. A53T:LNA | 0.004 |
| WT:ntLNA vs. A53T:ntLNA | 0.146 | | |

**Supplemental table 4.** Ratio of $TH^+/MAP2^+$ cells per imaged field. N=6 cultures per condition and 6 fields (3x3 images) per culture.

| Detection | | | | | |
|---|---|---|---|---|---|
| radius | auto | matched-filter | False | peaks | negative |
| N_t | 3 | alignment | True | isolation | False |
| spike_thresh | 7 | | | | |
| **Filtering** | | | | | |
| cut_off | 300, auto | filter | True | remove_median | True |
| **Triggers** | | | | | |
| trig_unit | ms | clean_artefact | False | ignore_times | False |
| **Whitening** | | | | | |
| chunk_size | 10 | safety_time | auto | output_dim | 5 |
| temporal | False | spatial | True | nb_elts | 0.8 |
| max_elts | 1000 | | | | |
| **Clustering** | | | | | |
| extraction | median-raw | safety_space | True | merging_param | default |
| safety_time | auto | max_elts | 10000 | cc_merge | 0.975 |
| nb_elts | 0.8 | nclus_min | 0.002 | dispersion | (5, 5) |
| max_clusters | 10 | nb_repeats | 3 | noise_thr | 0.8 |
| smart_search | True | smart_select | False | remove_mixture | True |
| merging_method | distance | cc_mixtures | 0.75 | | |
| **Fitting** | | | | | |
| chunk_size | 1 | amp_limits | (0.3, 5) | amp_auto | True |
| max_chunk | inf | collect_all | False | | |
| **Merging** | | | | | |
| cc_overlap | 0.85 | cc_bin | 2 | correct_lag | True |
| default_lag | 5 | auto_mode | 0.75 | | |
| **Validating** | | | | | |
| nearest_elec | auto | max_iter | 200 | learning_rate | 0.001 |
| roc_sampling | 10 | test_size | 0.3 | radius_factor | 0.5 |
| juxta_dtype | uint16 | juxta_thresh | 6 | juxta_valley | False |
| filter | True | | | | |
| **Extracting** | | | | | |
| safety_time | 1 | max_elts | 1000 | output_dim | 5 |
| cc_merge | 0.975 | noise_thr | 0.8 | | |

**Supplemental table 5.** Spike sorting parameters for SpyKING Circus 0.8.5**.**

| AMPL [µV] | WT | A53T |
|---|---|---|
| Week 1 | 33.9 ± 0.7 | 35.0 ± 1.0 |
| Week 2 | 35.7 ± 0.6 | 36.3 ± 0.9 |
| Week 3 | 36.2 ± 0.5 | 36.6 ± 0.8 |
| Week 4 | 36.1 ± 1.1 | 36.2 ± 1.5 |
| Week 5 | 35.2 ± 1.5 | 35.8 ± 1.5 |
| Mixed model effects | p-Values | F statistics |
| Time WT (T-WT) | <0.0001 | F(4.00,58.80) = 24.63 |
| Time A53T (T-A53T) | 0.0003 | F(4.00,62.00) = 8.85 |
| Genotype (G) | >0.9999 | F(1.00,33.15) = 4.02 |
| Genotype x Time (GxT) | >0.9999 | F(4.00,120.52) = 1.79 |
| HLFW [ms] | WT | A53T |
| Week 1 | 2.72 ± 0.26 | 2.88 ± 0.27 |
| Week 2 | 3.37 ± 0.21 | 3.34 ± 0.12 |
| Week 3 | 3.69 ± 0.19 | 3.50 ± 0.15 |
| Week 4 | 3.69 ± 0.17 | 3.63 ± 0.03 |
| Week 5 | 3.51 ± 0.53 | 3.53 ± 0.27 |
| Mixed model effects | p-Values | F statistics |
| Time WT (T-WT) | <0.0001 | F(4.00,55.95) = 49.83 |
| Time A53T (T-A53T) | <0.0001 | F(4.00,53.75) = 38.45 |
| Genotype (G) | >0.9999 | F(1.00,33.01) = 0.01 |
| Genotype x Time (GxT) | 0.2952 | F(4.00,110.96) = 3.52 |
| ASYM | WT | A53T |
| Week 1 | -0.063 ± 0.016 | -0.089 ± 0.031 |
| Week 2 | -0.14 ± 0.04 | -0.15 ± 0.07 |
| Week 3 | -0.17 ± 0.07 | -0.18 ± 0.07 |
| Week 4 | -0.14 ± 0.06 | -0.21 ± 0.07 |
| Week 5 | -0.14 ± 0.07 | -0.22 ± 0.07 |
| Mixed model effects | p-Values | F statistics |
| Time WT (T-WT) | <0.0001 | F(4.00,53.13) = 12.88 |
| Time A53T (T-A53T) | <0.0001 | F(4.00,62.06) = 21.06 |
| Genotype (G) | >0.9999 | F(1.00,33.93) = 3.46 |
| Genotype x Time (GxT) | 0.3203 | F(4.00,115.33) = 3.47 |
| T2PR | WT | A53T |
| Week 1 | 14.6 ± 1.6 | 14.7 ± 1.8 |
| Week 2 | 12.8 ± 1.3 | 13.2 ± 1.3 |
| Week 3 | 11.6 ± 1.3 | 12.5 ± 0.9 |
| Week 4 | 12.0 ± 2.0 | 12.0 ± 1.5 |
| Week 5 | 12.2 ± 2.6 | 12.4 ± 1.6 |
| Mixed model effects | p-Values | F statistics |
| Time WT (T-WT) | <0.0001 | F(4.00,52.04) = 12.02 |
| Time A53T (T-A53T) | <0.0001 | F(4.00,59.40) = 11.04 |
| Genotype (G) | >0.9999 | F(1.00,34.57) = 0.39 |
| Genotype x Time (GxT) | >0.9999 | F(4.00,110.33) = 0.79 |
| T2PD [ms] | WT | A53T |
| Week 1 | 13.2 ± 0.7 | 13.2 ± 0.6 |
| Week 2 | 13.3 ± 0.3 | 12.9 ± 0.3 |
| Week 3 | 13.5 ± 0.4 | 13.1 ± 0.3 |
| Week 4 | 13.7 ± 0.5 | 13.5 ± 0.1 |
| Week 5 | 13.9 ± 0.7 | 13.6 ± 0.4 |
| Mixed model effects | p-Values | F statistics |
| Time WT (T-WT) | <0.0001 | F(4.00,57.80) = 11.86 |
| Time A53T (T-A53T) | <0.0001 | F(4.00,54.23) = 16.07 |
| Genotype (G) | >0.9999 | F(1.00,33.49) = 2.66 |
| Genotype x Time (GxT) | >0.9999 | F(4.00,113.61) = 2.71 |

| AUCP1 | WT | A53T |
|---|---|---|
| Week 1 | 1.01 ± 0.22 | 1.12 ± 0.16 |
| Week 2 | 1.45 ± 0.14 | 1.33 ± 0.18 |
| Week 3 | 1.61 ± 0.11 | 1.49 ± 0.09 |
| Week 4 | 1.66 ± 0.08 | 1.64 ± 0.07 |
| Week 5 | 1.73 ± 0.17 | 1.77 ± 0.08 |
| Mixed model effects | p-Values | F statistics |
| Time WT (T-WT) | <0.0001 | F(4.00,54.75) = 73.24 |
| Time A53T (T-A53T) | <0.0001 | F(4.00,76.00) = 58.67 |
| Genotype (G) | >0.9999 | F(1.00,32.60) = 0.63 |
| Genotype x Time (GxT) | 0.0148 | F(4.00,114.72) = 5.44 |
| AUCP2 | WT | A53T |
| Week 1 | 5.96 ± 1.26 | 6.20 ± 1.20 |
| Week 2 | 7.99 ± 0.80 | 7.26 ± 0.87 |
| Week 3 | 8.85 ± 0.88 | 7.96 ± 0.66 |
| Week 4 | 9.10 ± 0.99 | 8.37 ± 0.73 |
| Week 5 | 7.98 ± 2.98 | 8.69 ± 1.16 |
| Mixed model effects | p-Values | F statistics |
| Time WT (T-WT) | <0.0001 | F(4.00,58.48) = 19.09 |
| Time A53T (T-A53T) | <0.0001 | F(4.00,61.00) = 42.46 |
| Genotype (G) | >0.9999 | F(1.00,35.63) = 0.39 |
| Genotype x Time (GxT) | 0.0333 | F(4.00,119.69) = 4.90 |
| AUCT | WT | A53T |
| Week 1 | 21.6 ± 1.9 | 22.5 ± 1.5 |
| Week 2 | 24.5 ± 1.2 | 23.7 ± 0.4 |
| Week 3 | 25.7 ± 1.1 | 24.7 ± 0.6 |
| Week 4 | 25.8 ± 1.0 | 25.3 ± 0.3 |
| Week 5 | 25.8 ± 1.7 | 25.1 ± 1.1 |
| Mixed model effects | p-Values | F statistics |
| Time WT (T-WT) | <0.0001 | F(4.00,54.98) = 65.07 |
| Time A53T (T-A53T) | <0.0001 | F(4.00,53.86) = 29.60 |
| Genotype (G) | >0.9999 | F(1.00,35.20) = 0.60 |
| Genotype x Time (GxT) | 0.0029 | F(4.00,109.52) = 6.54 |
| RISE [µV/ms] | WT | A53T |
| Week 1 | 1.08 ± 0.09 | 1.07 ± 0.09 |
| Week 2 | 0.95 ± 0.03 | 0.99 ± 0.03 |
| Week 3 | 0.93 ± 0.04 | 0.96 ± 0.03 |
| Week 4 | 0.93 ± 0.05 | 0.93 ± 0.02 |
| Week 5 | 0.91 ± 0.06 | 0.94 ± 0.02 |
| Mixed model effects | p-Values | F statistics |
| Time WT (T-WT) | <0.0001 | F(4.00,47.52) = 33.96 |
| Time A53T (T-A53T) | <0.0001 | F(4.00,53.27) = 23.11 |
| Genotype (G) | >0.9999 | F(1.00,31.36) = 0.79 |
| Genotype x Time (GxT) | >0.9999 | F(4.00,100.46) = 2.56 |
| DECAY [µV/ms] | WT | A53T |
| Week 1 | -0.12 ± 0.01 | -0.12 ± 0.02 |
| Week 2 | -0.080 ± 0.014 | -0.073 ± 0.007 |
| Week 3 | -0.064 ± 0.006 | -0.062 ± 0.005 |
| Week 4 | -0.069 ± 0.011 | -0.060 ± 0.004 |
| Week 5 | -0.072 ± 0.016 | -0.070 ± 0.011 |
| Mixed model effects | p-Values | F statistics |
| Time WT (T-WT) | <0.0001 | F(4.00,48.15) = 102.91 |
| Time A53T (T-A53T) | <0.0001 | F(4.00,52.96) = 61.87 |
| Genotype (G) | >0.9999 | F(1.00,28.56) = 2.45 |
| Genotype x Time (GxT) | >0.9999 | F(4.00,101.62) = 0.44 |

| ISIM [s] | WT | A53T |
|---|---|---|
| Week 1 | 180 ± 64 | 160 ± 74 |
| Week 2 | 85.8 ± 45.2 | 54.7 ± 21.1 |
| Week 3 | 32.6 ± 20.0 | 27.6 ± 9.6 |
| Week 4 | 41.2 ± 29.9 | 26.8 ± 14.0 |
| Week 5 | 101 ± 92 | 76.7 ± 61.0 |
| Mixed model effects | p-Values | F statistics |
| Time WT (T-WT) | <0.0001 | F(4.00,54.61) = 33.56 |
| Time A53T (T-A53T) | <0.0001 | F(4.00,55.55) = 31.85 |
| Genotype (G) | >0.9999 | F(1.00,32.72) = 2.96 |
| Genotype x Time (GxT) | >0.9999 | F(4.00,110.86) = 0.17 |
| ISIV | WT | A53T |
| Week 1 | 2338±1087 | 2001±1241 |
| Week 2 | 832 ± 587 | 413 ± 230 |
| Week 3 | 224 ± 190 | 169 ± 127 |
| Week 4 | 192 ± 109 | 121 ± 80 |
| Week 5 | 453 ± 428 | 751 ± 798 |
| Mixed model effects | p-Values | F statistics |
| Time WT (T-WT) | <0.0001 | F(4.00,46.95) = 34.84 |
| Time A53T (T-A53T) | <0.0001 | F(4.00,50.90) = 24.70 |
| Genotype (G) | >0.9999 | F(1.00,25.86) = 1.03 |
| Genotype x Time (GxT) | >0.9999 | F(4.00,97.98) = 0.68 |
| ISICV | WT | A53T |
| Week 1 | 1.09 ± 0.04 | 1.12 ± 0.08 |
| Week 2 | 1.44 ± 0.25 | 1.57 ± 0.31 |
| Week 3 | 2.12 ± 0.35 | 2.03 ± 0.27 |
| Week 4 | 2.06 ± 0.45 | 1.98 ± 0.19 |
| Week 5 | 1.86 ± 0.56 | 1.67 ± 0.23 |
| Mixed model effects | p-Values | F statistics |
| Time WT (T-WT) | <0.0001 | F(4.00,59.66) = 36.57 |
| Time A53T (T-A53T) | <0.0001 | F(4.00,55.69) = 43.52 |
| Genotype (G) | >0.9999 | F(1.00,33.28) = 0.39 |
| Genotype x Time (GxT) | >0.9999 | F(4.00,118.59) = 2.38 |
| PACF | WT | A53T |
| Week 1 | -0.022 ± 0.052 | -0.040 ± 0.032 |
| Week 2 | -0.050 ± 0.028 | -0.044 ± 0.035 |
| Week 3 | -0.0044 ± 0.0581 | -0.023 ± 0.028 |
| Week 4 | 0.013 ± 0.067 | -0.022 ± 0.031 |
| Week 5 | 0.019 ± 0.065 | -0.0041 ± 0.0262 |
| Mixed model effects | p-Values | F statistics |
| Time WT (T-WT) | 0.0318 | F(4.00,57.25) = 5.32 |
| Time A53T (T-A53T) | 0.0755 | F(4.00,57.56) = 4.68 |
| Genotype (G) | >0.9999 | F(1.00,33.40) = 3.00 |
| Genotype x Time (GxT) | >0.9999 | F(4.00,115.49) = 1.08 |
| SCRF [Hz] | WT | A53T |
| Week 1 | 2.02 ± 0.13 | 1.53 ± 0.41 |
| Week 2 | 0.77 ± 0.39 | 0.58 ± 0.27 |
| Week 3 | 0.25 ± 0.10 | 0.46 ± 0.15 |
| Week 4 | 0.27 ± 0.08 | 0.52 ± 0.16 |
| Week 5 | 0.77 ± 0.63 | 0.90 ± 0.32 |
| Mixed model effects | p-Values | F statistics |
| Time WT (T-WT) | <0.0001 | F(4.00,48.67) = 77.20 |
| Time A53T (T-A53T) | <0.0001 | F(4.00,54.27) = 48.51 |
| Genotype (G) | >0.9999 | F(1.00,24.79) = 0.21 |
| Genotype x Time (GxT) | 0.0001 | F(4.00,103.20) = 8.90 |

| SCRM | WT | A53T |
|---|---|---|
| Week 1 | 44.9 ± 16.5 | 49.7 ± 17.4 |
| Week 2 | 127 ± 58 | 165 ± 80 |
| Week 3 | 400 ± 175 | 236 ± 99 |
| Week 4 | 355 ± 185 | 223 ± 52 |
| Week 5 | 319 ± 236 | 166 ± 59 |
| Mixed model effects | p-Values | F statistics |
| Time WT (T-WT) | <0.0001 | F(4.00,58.09) = 26.64 |
| Time A53T (T-A53T) | <0.0001 | F(4.00,59.97) = 23.41 |
| Genotype (G) | 0.2258 | F(1.00,35.91) = 8.10 |
| Genotype x Time (GxT) | <0.0001 | F(4.00,118.62) = 8.81 |
| RFIT | WT | A53T |
| Week 1 | -0.0050 ± 0.0024 | -0.0045 ± 0.0022 |
| Week 2 | -0.0070 ± 0.0012 | -0.0067 ± 0.0026 |
| Week 3 | -0.0090 ± 0.0020 | -0.0059 ± 0.0024 |
| Week 4 | -0.0076 ± 0.0012 | -0.0058 ± 0.0028 |
| Week 5 | -0.0079 ± 0.0021 | -0.0049 ± 0.0018 |
| Mixed model effects | p-Values | F statistics |
| Time WT (T-WT) | <0.0001 | F(4.00,69.00) = 9.94 |
| Time A53T (T-A53T) | >0.9999 | F(4.00,79.00) = 2.42 |
| Genotype (G) | <0.0001 | F(1.00,148.00) = 25.61 |
| Genotype x Time (GxT) | 0.6395 | F(4.00,148.00) = 2.99 |
| IBIV | WT | A53T |
| Week 1 | 0 | 1.45 ± 2.24 |
| Week 2 | 1.34 ± 1.13 | 0.78 ± 0.60 |
| Week 3 | 1.01 ± 0.63 | 5.11 ± 2.77 |
| Week 4 | 3.17 ± 3.12 | 4.70 ± 2.67 |
| Week 5 | 4.81 ± 4.97 | 4.85 ± 2.88 |
| Mixed model effects | p-Values | F statistics |
| Time WT (T-WT) | 0.0046 | F(4.00,51.57) = 6.95 |
| Time A53T (T-A53T) | <0.0001 | F(4.00,52.75) = 10.62 |
| Genotype (G) | 0.2201 | F(1.00,29.31) = 8.38 |
| Genotype x Time (GxT) | 0.1655 | F(4.00,104.65) = 3.91 |
| IBIM [s] | WT | A53T |
| Week 1 | 0 | 12.0 ± 7.8 |
| Week 2 | 10.8 ± 3.0 | 10.8 ± 4.1 |
| Week 3 | 14.4 ± 2.8 | 8.53 ± 1.87 |
| Week 4 | 15.4 ± 3.5 | 7.75 ± 1.21 |
| Week 5 | 16.8 ± 6.2 | 7.90 ± 1.69 |
| Mixed model effects | p-Values | F statistics |
| Time WT (T-WT) | <0.0001 | F(4.00,52.16) = 39.22 |
| Time A53T (T-A53T) | 0.0185 | F(4.00,57.86) = 5.72 |
| Genotype (G) | 0.5280 | F(1.00,28.21) = 6.43 |
| Genotype x Time (GxT) | <0.0001 | F(4.00,108.78) = 38.96 |
| MBD [s] | WT | A53T |
| Week 1 | 0 | 7.32 ± 6.01 |
| Week 2 | 8.43 ± 2.07 | 5.53 ± 1.29 |
| Week 3 | 6.82 ± 0.98 | 4.94 ± 0.73 |
| Week 4 | 7.84 ± 1.38 | 4.46 ± 0.26 |
| Week 5 | 6.70 ± 2.34 | 4.87 ± 0.48 |
| Mixed model effects | p-Values | F statistics |
| Time WT (T-WT) | <0.0001 | F(4.00,65.00) = 46.10 |
| Time A53T (T-A53T) | >0.9999 | F(4.00,72.00) = 2.32 |
| Genotype (G) | >0.9999 | F(1.00,137.00) = 1.74 |
| Genotype x Time (GxT) | <0.0001 | F(4.00,137.00) = 21.93 |

| | | |
|---|---|---|
| VBD | WT | A53T |
| Week 1 | 0 | 0.37 ± 0.38 |
| Week 2 | 1.25 ± 1.16 | 0.39 ± 0.45 |
| Week 3 | 0.50 ± 0.36 | 1.69 ± 2.27 |
| Week 4 | 1.34 ± 1.33 | 0.73 ± 0.89 |
| Week 5 | 0.59 ± 0.39 | 2.39 ± 1.49 |
| Mixed model effects | p-Values | F statistics |
| Time WT (T-WT) | 0.0466 | F(4.00,43.61) = 5.27 |
| Time A53T (T-A53T) | 0.0151 | F(4.00,44.32) = 6.17 |
| Genotype (G) | >0.9999 | F(1.00,28.65) = 2.79 |
| Genotype x Time (GxT) | 0.0007 | F(4.00,87.68) = 7.67 |
| INTRABF [Hz] | WT | A53T |
| Week 1 | 0 | 0.35 ± 0.19 |
| Week 2 | 0.83 ± 0.46 | 1.17 ± 0.65 |
| Week 3 | 2.08 ± 0.91 | 2.25 ± 0.47 |
| Week 4 | 2.13 ± 1.13 | 2.42 ± 0.69 |
| Week 5 | 1.86 ± 1.17 | 1.55 ± 0.58 |
| Mixed model effects | p-Values | F statistics |
| Time WT (T-WT) | <0.0001 | F(4.00,52.30) = 36.22 |
| Time A53T (T-A53T) | <0.0001 | F(4.00,57.56) = 47.03 |
| Genotype (G) | >0.9999 | F(1.00,34.95) = 1.18 |
| Genotype x Time (GxT) | 0.9090 | F(4.00,110.78) = 2.80 |
| INTERBF [Hz] | WT | A53T |
| Week 1 | 0 | 0.17 ± 0.14 |
| Week 2 | 0.32 ± 0.22 | 0.26 ± 0.19 |
| Week 3 | 0.39 ± 0.24 | 0.37 ± 0.24 |
| Week 4 | 0.49 ± 0.30 | 0.39 ± 0.23 |
| Week 5 | 0.41 ± 0.27 | 0.41 ± 0.26 |
| Mixed model effects | p-Values | F statistics |
| Time WT (T-WT) | <0.0001 | F(4.00,51.77) = 24.60 |
| Time A53T (T-A53T) | <0.0001 | F(4.00,58.15) = 15.66 |
| Genotype (G) | >0.9999 | F(1.00,34.19) = 0.07 |
| Genotype x Time (GxT) | 0.0002 | F(4.00,109.70) = 8.48 |
| BRT [s] | WT | A53T |
| Week 1 | 0 | 1.64 ± 1.21 |
| Week 2 | 2.63 ± 0.80 | 0.89 ± 0.38 |
| Week 3 | 1.02 ± 0.33 | 0.72 ± 0.22 |
| Week 4 | 0.84 ± 0.26 | 0.62 ± 0.18 |
| Week 5 | 0.49 ± 0.12 | 0.64 ± 0.21 |
| Mixed model effects | p-Values | F statistics |
| Time WT (T-WT) | <0.0001 | F(4.00,44.35) = 85.53 |
| Time A53T (T-A53T) | 0.0006 | F(4.00,74.00) = 8.08 |
| Genotype (G) | >0.9999 | F(1.00,18.13) = 1.15 |
| Genotype x Time (GxT) | <0.0001 | F(4.00,95.03) = 36.73 |
| BRV [coactivity/s] | WT | A53T |
| Week 1 | 0 | 14.6 ± 7.4 |
| Week 2 | 27.6 ± 20.0 | 238 ± 222 |
| Week 3 | 485 ± 370 | 648 ± 369 |
| Week 4 | 556 ± 398 | 660 ± 399 |
| Week 5 | 630 ± 580 | 562 ± 365 |
| Mixed model effects | p-Values | F statistics |
| Time WT (T-WT) | <0.0001 | F(4.00,49.58) = 17.03 |
| Time A53T (T-A53T) | <0.0001 | F(4.00,58.50) = 24.33 |
| Genotype (G) | >0.9999 | F(1.00,35.17) = 0.82 |
| Genotype x Time (GxT) | >0.9999 | F(4.00,107.99) = 1.33 |

| BDT [s] | WT | A53T |
|---|---|---|
| Week 1 | 0 | 1.10 ± 0.68 |
| Week 2 | 1.44 ± 0.25 | 1.03 ± 0.41 |
| Week 3 | 1.87 ± 0.67 | 1.13 ± 0.40 |
| Week 4 | 2.36 ± 0.72 | 1.05 ± 0.38 |
| Week 5 | 2.27 ± 1.30 | 0.86 ± 0.28 |
| Mixed model effects | p-Values | F statistics |
| Time WT (T-WT) | <0.0001 | F(4.00,53.41) = 22.67 |
| Time A53T (T-A53T) | >0.9999 | F(4.00,55.57) = 0.92 |
| Genotype (G) | 0.0053 | F(1.00,31.22) = 18.22 |
| Genotype x Time (GxT) | <0.0001 | F(4.00,114.62) = 23.05 |
| BDV [coactivity/s] | WT | A53T |
| Week 1 | 0 | -16.3 ± 8.6 |
| Week 2 | -49.7 ± 39.4 | -229 ± 203 |
| Week 3 | -247 ± 131 | -395 ± 218 |
| Week 4 | -180 ± 128 | -399 ± 256 |
| Week 5 | -118 ± 91 | -387 ± 269 |
| Mixed model effects | p-Values | F statistics |
| Time WT (T-WT) | <0.0001 | F(4.00,49.34) = 25.43 |
| Time A53T (T-A53T) | <0.0001 | F(4.00,59.69) = 21.85 |
| Genotype (G) | 0.0401 | F(1.00,35.18) = 12.23 |
| Genotype x Time (GxT) | 0.3505 | F(4.00,109.62) = 3.42 |
| SYNC | WT | A53T |
| Week 1 | 0.0029 ± 0.0007 | 0.0040 ± 0.0011 |
| Week 2 | 0.0053 ± 0.0011 | 0.0074 ± 0.0025 |
| Week 3 | 0.0083 ± 0.0023 | 0.0093 ± 0.0026 |
| Week 4 | 0.0082 ± 0.0026 | 0.0094 ± 0.0019 |
| Week 5 | 0.0073 ± 0.0030 | 0.0077 ± 0.0020 |
| Mixed model effects | p-Values | F statistics |
| Time WT (T-WT) | <0.0001 | F(4.00,59.20) = 50.11 |
| Time A53T (T-A53T) | <0.0001 | F(4.00,61.55) = 43.28 |
| Genotype (G) | >0.9999 | F(1.00,35.35) = 4.24 |
| Genotype x Time (GxT) | >0.9999 | F(4.00,120.78) = 2.01 |
| NRF [Hz] | WT | A53T |
| Week 1 | 0.0014 ± 0.0005 | 0.042 ± 0.006 |
| Week 2 | 0.054 ± 0.010 | 0.058 ± 0.010 |
| Week 3 | 0.047 ± 0.009 | 0.085 ± 0.019 |
| Week 4 | 0.043 ± 0.010 | 0.084 ± 0.011 |
| Week 5 | 0.045 ± 0.012 | 0.088 ± 0.015 |
| Mixed model effects | p-Values | F statistics |
| Time WT (T-WT) | <0.0001 | F(4.00,50.80) = 53.95 |
| Time A53T (T-A53T) | <0.0001 | F(4.00,57.86) = 38.38 |
| Genotype (G) | <0.0001 | F(1.00,32.78) = 221.50 |
| Genotype x Time (GxT) | <0.0001 | F(4.00,109.50) = 17.68 |
| NRM | WT | A53T |
| Week 1 | 69.8 ± 23.9 | 183 ± 71 |
| Week 2 | 392 ± 122 | 237 ± 129 |
| Week 3 | 355 ± 134 | 160 ± 65 |
| Week 4 | 256 ± 58 | 161 ± 58 |
| Week 5 | 207 ± 105 | 123 ± 45 |
| Mixed model effects | p-Values | F statistics |
| Time WT (T-WT) | <0.0001 | F(4.00,58.80) = 37.99 |
| Time A53T (T-A53T) | 0.0012 | F(4.00,61.42) = 7.75 |
| Genotype (G) | 0.0043 | F(1.00,32.07) = 18.73 |
| Genotype x Time (GxT) | <0.0001 | F(4.00,120.66) = 21.64 |

| NRFIT | WT | A53T |
|---|---|---|
| Week 1 | -0.0028 ± 0.0033 | -0.0044 ± 0.0019 |
| Week 2 | -0.0036 ± 0.0015 | -0.0016 ± 0.0003 |
| Week 3 | -0.0017 ± 0.0002 | -0.0014 ± 0.0001 |
| Week 4 | -0.0018 ± 0.0003 | -0.0013 ± 0.0002 |
| Week 5 | -0.0018 ± 0.0003 | -0.0014 ± 0.0002 |
| Mixed model effects | p-Values | F statistics |
| Time WT (T-WT) | 0.0001 | $F(4.00,42.84) = 10.77$ |
| Time A53T (T-A53T) | <0.0001 | $F(4.00,44.47) = 34.49$ |
| Genotype (G) | >0.9999 | $F(1.00,23.53) = 2.08$ |
| Genotype x Time (GxT) | <0.0001 | $F(4.00,87.13) = 24.20$ |

**Supplemental table 6.** Complementary data related to Figure 3. Reported are Mean ± SD of all features for each week and genotype and the results of linear mixed-effect models assessing developmental (T), genotype (G) and interaction effects (GxT). p-Values were adjusted for the number of comparisons (Bonferroni correction). Sample size for these analyses were 18 WT and 19 A53T cultures.

| Genotype | Treatment | HTRF [intensity] | | | | | | | | | Mean ± SD |
|---|---|---|---|---|---|---|---|---|---|---|---|
| WT | Untreated | 63.8 | 64.6 | 65.7 | 70.2 | 70.1 | 70.4 | 64.2 | 65.3 | 65.5 | 66.6 ± 2.8 |
| | ntLNA | 74.3 | 74.9 | 73.4 | 70.7 | 71.1 | 70.3 | 71.1 | 71.2 | 71.2 | 72.0 ± 1.7 |
| | LNA | 5.4 | 5.6 | 5.5 | 5.5 | 5.6 | 5.5 | 5.2 | 5.3 | 5.1 | 5.4 ± 0.2 |
| A53T | Untreated | 74.3 | 72.9 | 72.6 | 63.9 | 66 | 60.8 | 63.7 | 63.8 | 64.5 | 66.9 ± 4.9 |
| | ntLNA | 71.3 | 71.9 | 72.5 | 67.9 | 67.2 | 68.7 | 39.8* | 39.5* | 40.0* | 69.9 ± 2.3 |
| | LNA | 5.5 | 5.9 | 5.8 | 6.5 | 6.3 | 6.4 | 6.5 | 6.7 | 6.4 | 6.2 ± 0.4 |

| ANOVA table | SS | DF | MS | F (DFn, DFd) | p value |
|---|---|---|---|---|---|
| Interaction | 6.2 | 2 | 3.1 | F (2, 11) = 0.4 | p=0.705 |
| Mutation | 0.5 | 1 | 0.5 | F (1, 11) = 0.05 | p=0.820 |
| Treatment | 15282.7 | 2 | 7641.3 | F (2, 11) = 885.3 | p<0.001 |
| Residual | 94.9 | 11 | 8.6 | | |

| Tukey's multiple comparisons test | Adjusted p value | Tukey's multiple comparisons test | Adjusted p value |
|---|---|---|---|
| WT:Untreated vs. WT:ntLNA | >0.999 | WT:ntLNA vs. A53T:LNA | <0.001 |
| WT:Untreated vs. WT:LNA | 0.607 | WT:LNA7 vs. A53T:Untreated | >0.999 |
| WT:Untreated vs. A53T:Untreated | 0.643 | WT:LNA vs. A53T:ntLNA | 0.999 |
| WT:Untreated vs. A53T:ntLNA | 0.184 | WT:LNA vs. A53T:LNA | <0.001 |
| WT:Untreated vs. A53T:LNA | <0.001 | A53T:Untreated vs. A53T:ntLNA | 0.999 |
| WT:ntLNA vs. WT:LNA | 0.530 | A53T:Untreated vs. A53T:LNA | <0.001 |
| WT:ntLNA vs. A53T:Untreated | 0.566 | A53T:ntLNA vs. A53T:LNA | 0.004 |
| WT:ntLNA vs. A53T:ntLNA | 0.146 | | |

**Supplemental table 7.** Quantification of total α-synuclein levels by Homogeneous Time Resolved Fluorescence (HTRF) assay. For each condition, three biological replicates (different cultures) with three technical replicates each (grouped by thick black lines) were analyzed. The samples indicated with asterisks were considered outliers and excluded from the analysis

| Genotype | Treatment | Somatic α-synuclein levels [intensity] | | | | | | | | | Mean ± SD |
|---|---|---|---|---|---|---|---|---|---|---|---|
| WT | Untreated | 817.6 | 674.8 | 683.3 | 744.5 | 719.4 | 701.7 | 665 | 698.9 | 667.7 | 721.4 ± 57.6 |
| | | 802 | 726.2 | 754 | 660.3 | 621.8 | 691.5 | 808.7 | 753.2 | 794 | |
| | ntLNA | 791.9 | 712.4 | 824.3 | 584.2 | 648.6 | 615.6 | 635.1 | 734.2 | 683.4 | 723.1 ± 91.3 |
| | | 731.6 | 713.4 | 702.7 | 600.6 | 637.7 | 559.5 | 821 | 830.7 | 792.4 | |
| | LNA | 271 | 258.1 | 273.2 | 256 | 296.3 | 272.9 | 264.2 | 276 | 256.6 | 272.4 ± 13.9 |
| | | 284.6 | 299.8 | 265.9 | 270.5 | 273.2 | 249.6 | 269.6 | 272 | 293 | |
| A53T | Untreated | 781.4 | 786.2 | 720 | 690.3 | 806.5 | 859.1 | 689.8 | 733.6 | 690.9 | 721.5 ± 75.6 |
| | | 791.9 | 712.4 | 824.3 | 584.2 | 648.6 | 615.6 | 635.1 | 734.2 | 683.4 | |
| | ntLNA | 878.2 | 869.9 | 892.5 | 872.8 | 801.6 | 793.7 | 734.9 | 767.6 | 684 | 775.1 ± 66.9 |
| | | 733.4 | 764.7 | 738.6 | 701.3 | 675.2 | 721.6 | 793.7 | 756.1 | 771.1 | |
| | LNA | 291.5 | 252.7 | 257.6 | 273.9 | 298.6 | 283.8 | 269.3 | 288.6 | 287.4 | 275.9 ± 12.6 |
| | | 274.4 | 281.3 | 282 | 271.3 | 258.3 | 274.9 | 267.9 | 265.9 | 286.1 | |

| ANOVA table | SS | DF | MS | F (DFn, DFd) | p value |
|---|---|---|---|---|---|
| Interaction | 15148.8 | 2 | 7574.3 | F (2, 102) = 2.0 | p=0.134 |
| Mutation | 9292.6 | 1 | 9292.6 | F (1, 102) = 2.5 | p=0.116 |
| Treatment | 5117295.0 | 2 | 2558647.5 | F (2, 102) = 691.9 | p<0.001 |
| Residual | 377216.7 | 102 | 3698.2 | | |

| Tukey's multiple comparisons test | Adjusted p value | Tukey's multiple comparisons test | Adjusted p value |
|---|---|---|---|
| WT:Untreated vs. WT:ntLNA | >0.999 | WT:ntLNA vs. A53T:LNA | <0.001 |
| WT:Untreated vs. WT:LNA | <0.001 | WT:LNA7 vs. A53T:Untreated | <0.001 |
| WT:Untreated vs. A53T:Untreated | >0.999 | WT:LNA vs. A53T:ntLNA | <0.001 |
| WT:Untreated vs. A53T:ntLNA | 0.132 | WT:LNA vs. A53T:LNA | >0.999 |
| WT:Untreated vs. A53T:LNA | <0.001 | A53T:Untreated vs. A53T:ntLNA | 0.134 |
| WT:ntLNA vs. WT:LNA | <0.001 | A53T:Untreated vs. A53T:LNA | <0.001 |
| WT:ntLNA vs. A53T:Untreated | >0.999 | A53T:ntLNA vs. A53T:LNA | <0.001 |
| WT:ntLNA vs. A53T:ntLNA | 0.163 | | |

**Supplemental table 8.** Quantification of somatic α-synuclein levels by immunocytochemical staining analysis. For each condition, three cultures were analyzed by quantifying the intensity of six fields, each consisting of 3x3 images (N=54 images per condition).

| Genotype | Treatment | Somatic phosphorylated α-synuclein levels [intensity] | | | | | | | | | Mean ± SD |
|---|---|---|---|---|---|---|---|---|---|---|---|
| WT | Untreated | 281.3 | 197.1 | 210.3 | 258.5 | 231.5 | 216.1 | 197 | 207.9 | 201.7 | 225 ± 22.9 |
| | | 218.6 | 227.7 | 217.6 | 225.7 | 215.6 | 208.2 | 244.9 | 254.4 | 238.1 | |
| | ntLNA | 215.9 | 234.7 | 239.9 | 211.5 | 233.8 | 284 | 213 | 213.3 | 246.3 | 226.9 ±21.4 |
| | | 211.5 | 203.5 | 248.1 | 201.5 | 196 | 230.3 | 238.9 | 238.9 | 222.4 | |
| | LNA | 156.1 | 157.1 | 155.8 | 161.4 | 160.6 | 160.9 | 160.3 | 157.8 | 157.1 | 159.6 ± 2.4 |
| | | 163.7 | 161.1 | 160.7 | 160.6 | 157.2 | 157.7 | 161.2 | 164 | 159.6 | |
| A53T | Untreated | 239.6 | 247.8 | 265.4 | 233.2 | 245.9 | 290 | 207.7 | 207.5 | 271.3 | 235.1 ± 24.7 |
| | | 224.5 | 211.1 | 256.9 | 210.3 | 201.9 | 244 | 227.7 | 219.7 | 226.6 | |
| | ntLNA | 234.4 | 238.9 | 260.6 | 249.5 | 226.2 | 293.2 | 249 | 216 | 275.6 | 246.8 ± 20.4 |
| | | 238 | 232.4 | 245.3 | 227.2 | 227.5 | 236.3 | 253.5 | 275.5 | 264.1 | |
| | LNA | 159.1 | 175 | 160.4 | 163.2 | 171.1 | 166.2 | 163.9 | 167.3 | 163.8 | 165.2 ± 3.7 |
| | | 162.9 | 165.7 | 166.9 | 162.6 | 164.5 | 165.4 | 167.1 | 162.6 | 166.5 | |

| ANOVA table | SS | DF | MS | F (DFn, DFd) | p value |
|---|---|---|---|---|---|
| Interaction | 976.7 | 2 | 488.3 | F (2, 102) = 1.4 | p=0.241 |
| Mutation | 3791.4 | 1 | 3791.4 | F (1, 102) = 11.2 | p=0.001 |
| Treatment | 121986.8 | 2 | 60993.4 | F (2, 102) = 180.1 | p<0.001 |
| Residual | 34547.8 | 102 | 338.7 | | |

| Tukey's multiple comparisons test | Adjusted p value | Tukey's multiple comparisons test | Adjusted p value |
|---|---|---|---|
| WT:Untreated vs. WT:ntLNA | 0.999 | WT:ntLNA vs. A53T:LNA | <0.001 |
| WT:Untreated vs. WT:LNA | <0.001 | WT:LNA7 vs. A53T:Untreated | <0.001 |
| WT:Untreated vs. A53T:Untreated | 0.587 | WT:LNA vs. A53T:ntLNA | <0.001 |
| WT:Untreated vs. A53T:ntLNA | 0.008 | WT:LNA vs. A53T:LNA | 0.941 |
| WT:Untreated vs. A53T:LNA | <0.001 | A53T:Untreated vs. A53T:ntLNA | 0.396 |
| WT:ntLNA vs. WT:LNA | <0.001 | A53T:Untreated vs. A53T:LNA | <0.001 |
| WT:ntLNA vs. A53T:Untreated | 0.764 | A53T:ntLNA vs. A53T:LNA | <0.001 |
| WT:ntLNA vs. A53T:ntLNA | 0.018 | | |

**Supplemental table 9.** Quantification of somatic phosphorylated α-synuclein levels by immunocytochemical staining analysis. For each condition, three cultures were analyzed by quantifying the intensity of six fields, each consisting of 3x3 images (N=54 images per condition).

| Genotype | Treatment | Neuritic α-synuclein levels [intensity] | | | | | | | | | Mean ± SD |
|---|---|---|---|---|---|---|---|---|---|---|---|
| WT | Untreated | 286.1 | 419.8 | 389.7 | 284.2 | 425.7 | 380.5 | 290.7 | 429.2 | 395.9 | 348.6 ± 57.9 |
| | | 301.8 | 389.8 | 390.6 | 275.8 | 386.3 | 330.7 | 253.3 | 325.9 | 318.9 | |
| | ntLNA | 329.1 | 358.6 | 393.2 | 302.6 | 378.3 | 443.8 | 278.4 | 405.8 | 384.1 | 355.1 ± 48.6 |
| | | 322.8 | 402.2 | 363.6 | 296.6 | 377.3 | 388 | 259.8 | 344.4 | 362.8 | |
| | LNA | 160 | 164 | 161.9 | 158.4 | 182.6 | 175.4 | 168.4 | 178.8 | 175.8 | 165.4 ± 10.8 |
| | | 160.9 | 176.7 | 175.5 | 148.8 | 167.1 | 166.6 | 143 | 157.5 | 154.9 | |
| A53T | Untreated | 256.3 | 246.2 | 207.7 | 320.3 | 273.6 | 217.9 | 328 | 311.3 | 233 | 284.5 ± 42.6 |
| | | 343.9 | 331.9 | 246.7 | 325.7 | 297.9 | 272.5 | 325.3 | 309.7 | 273.5 | |
| | ntLNA | 307.1 | 254.1 | 191.5 | 315 | 303 | 241.5 | 332.9 | 306.7 | 242.9 | 301.1 ± 53.1 |
| | | 372.8 | 318.7 | 243.9 | 355.3 | 347.8 | 251.3 | 371.3 | 366.1 | 297.5 | |
| | LNA | 145.1 | 150.8 | 136.9 | 158.5 | 154.1 | 145.2 | 156.3 | 164.1 | 153.3 | 153.2 ± 7.1 |
| | | 163.4 | 155.8 | 149.4 | 158.5 | 161.3 | 152 | 157.3 | 149.7 | 146.1 | |

| ANOVA table | SS | DF | MS | F (DFn, DFd) | p value |
|---|---|---|---|---|---|
| Interaction | 13656.8 | 2 | 6828.4 | F (2, 102) = 3.9 | P=0.023 |
| Mutation | 50873.5 | 1 | 50873.5 | F (1, 102) = 29.0 | p<0.001 |
| Treatment | 640357.4 | 2 | 320178.7 | F (2, 102) = 182.1 | p<0.001 |
| Residual | 178891.9 | 102 | 1753.8 | | |

| Tukey's multiple comparisons test | Adjusted p value | Tukey's multiple comparisons test | Adjusted p value |
|---|---|---|---|
| WT:Untreated vs. WT:ntLNA | 0.997 | WT:ntLNA vs. A53T:LNA | <0.001 |
| WT:Untreated vs. WT:LNA | <0.001 | WT:LNA7 vs. A53T:Untreated | <0.001 |
| WT:Untreated vs. A53T:Untreated | <0.001 | WT:LNA vs. A53T:ntLNA | <0.001 |
| WT:Untreated vs. A53T:ntLNA | 0.012 | WT:LNA vs. A53T:LNA | 0.953 |
| WT:Untreated vs. A53T:LNA | <0.001 | A53T:Untreated vs. A53T:ntLNA | 0.843 |
| WT:ntLNA vs. WT:LNA | <0.001 | A53T:Untreated vs. A53T:LNA | <0.001 |
| WT:ntLNA vs. A53T:Untreated | <0.001 | A53T:ntLNA vs. A53T:LNA | <0.001 |
| WT:ntLNA vs. A53T:ntLNA | 0.003 | | |

**Supplemental table 10.** Quantification of neuritic α-synuclein levels by immunocytochemical staining analysis. For each condition, three cultures were analyzed by quantifying the intensity of six fields, each consisting of 3x3 images (N=54 images per condition).

| Genotype | Treatment | Neuritic phosphorylated α-synuclein levels [intensity] | | | | | | | | | Mean ± SD |
|---|---|---|---|---|---|---|---|---|---|---|---|
| WT | Untreated | 164.5 | 164 | 163.9 | 162.7 | 167.2 | 168.9 | 161.3 | 166.7 | 167.7 | 165.6 ± 3.3 |
| | | 165.5 | 169.8 | 171 | 161.1 | 163.8 | 166.6 | 159.5 | 166.5 | 170.3 | |
| | ntLNA | 161.2 | 162.4 | 163.3 | 165.1 | 168.1 | 171.6 | 162 | 169.2 | 165.8 | 165.1 ± 3.3 |
| | | 167.6 | 164 | 168.9 | 163 | 161.3 | 166 | 158.9 | 166.3 | 167.4 | |
| | LNA | 146.8 | 150 | 150.4 | 152.4 | 154.4 | 155.3 | 157.7 | 153.3 | 155.1 | 153.5 ± 2.9 |
| | | 152.9 | 157.3 | 154.9 | 150.9 | 153.8 | 153.8 | 152.8 | 158.6 | 153.1 | |
| A53T | Untreated | 160.6 | 160.3 | 155.3 | 168.9 | 163.6 | 161.9 | 166 | 163.3 | 160.6 | 164.6 ± 3.9 |
| | | 168.7 | 166.2 | 169.5 | 164.6 | 162.3 | 167.4 | 166.6 | 168.8 | 168.1 | |
| | ntLNA | 162.5 | 162.9 | 153.1 | 168.8 | 163.3 | 160.3 | 166.6 | 166 | 166.6 | 165.4 ± 4.2 |
| | | 168.7 | 169.4 | 164.4 | 167.8 | 168.8 | 163.9 | 169.7 | 169.3 | 164.2 | |
| | LNA | 149.2 | 148.2 | 146.9 | 153.4 | 152.3 | 150.7 | 152.5 | 157.8 | 155.8 | 153.1 ± 2.9 |
| | | 154.8 | 154.2 | 154.7 | 151.1 | 156.1 | 153.4 | 153.8 | 156.1 | 154 | |

| ANOVA table | SS | DF | MS | F (DFn, DFd) | p value |
|---|---|---|---|---|---|
| Interaction | 7.1 | 2 | 3.5 | F (2, 102) = 0.3 | p=0.743 |
| Mutation | 4.7 | 1 | 4.7 | F (1, 102) = 0.4 | p=0.530 |
| Treatment | 3385.5 | 2 | 1692.7 | F (2, 102) = 142.4 | p<0.001 |
| Residual | 1212.7 | 102 | 11.9 | | |

| Tukey's multiple comparisons test | Adjusted p value | Tukey's multiple comparisons test | Adjusted p value |
|---|---|---|---|
| WT:Untreated vs. WT:ntLNA | 0.998 | WT:ntLNA vs. A53T:LNA | <0.001 |
| WT:Untreated vs. WT:LNA | <0.001 | WT:LNA7 vs. A53T:Untreated | <0.001 |
| WT:Untreated vs. A53T:Untreated | 0.949 | WT:LNA vs. A53T:ntLNA | <0.001 |
| WT:Untreated vs. A53T:ntLNA | 0.999 | WT:LNA vs. A53T:LNA | 0.998 |
| WT:Untreated vs. A53T:LNA | <0.001 | A53T:Untreated vs. A53T:ntLNA | 0.986 |
| WT:ntLNA vs. WT:LNA | <0.001 | A53T:Untreated vs. A53T:LNA | <0.001 |
| WT:ntLNA vs. A53T:Untreated | 0.998 | A53T:ntLNA vs. A53T:LNA | <0.001 |
| WT:ntLNA vs. A53T:ntLNA | 0.999 | | |

**Supplemental table 11.** Quantification of neuritic phosphorylated α-synuclein levels by immunocytochemical staining analysis. For each condition, three cultures were analyzed by quantifying the intensity of six fields, each consisting of 3x3 images (N=54 images per condition).

| ISIM [s] | WT (Untreated) | WT (ntLNA) |
|---|---|---|
| Week 1 | 261 ± 14 | 273 ± 89 |
| Week 2 | 36.0 ± 0.9 | 83.2 ± 77.2 |
| Week 3 | 14.7 ± 0.4 | 17.9 ± 9.5 |
| Week 4 | 18.9 ± 4.5 | 14.0 ± 1.9 |
| Mixed model effects | p-Values | F statistics |
| Time WT (Untreated) | 0.4086 | F(3.00,2.78) = 29.53 |
| Time WT (ntLNA) | 0.0450 | F(3.00,2.05) = 625.45 |
| LNA-Treatment (L) | >0.9999 | F(1.00,3.26) = 0.13 |
| LNA x Time (LxT) | >0.9999 | F(3.00,5.57) = 1.27 |
| ISIV | WT (Untreated) | WT (ntLNA) |
| Week 1 | 3849±59 | 3619±1350 |
| Week 2 | 154 ± 75 | 854 ± 1007 |
| Week 3 | 53.8 ± 2.5 | 105 ± 105 |
| Week 4 | 96.4 ± 14.2 | 51.0 ± 18.2 |
| Mixed model effects | p-Values | F statistics |
| Time WT (Untreated) | 0.2430 | F(3.00,2.87) = 40.05 |
| Time WT (ntLNA) | <0.0001 | F(3.00,2.99) = 31863.73 |
| LNA-Treatment (L) | >0.9999 | F(1.00,3.48) = 0.25 |
| LNA x Time (LxT) | >0.9999 | F(3.00,5.64) = 1.01 |
| ISICV | WT (Untreated) | WT (ntLNA) |
| Week 1 | 1.05 ± 0.02 | 1.04 ± 0.08 |
| Week 2 | 2.07 ± 0.34 | 1.63 ± 0.58 |
| Week 3 | 3.27 ± 0.58 | 3.39 ± 0.60 |
| Week 4 | 3.45 ± 0.06 | 3.81 ± 0.06 |
| Mixed model effects | p-Values | F statistics |
| Time WT (Untreated) | 0.0706 | F(3.00,3.94) = 38.76 |
| Time WT (ntLNA) | 0.0113 | F(3.00,7.00) = 26.22 |
| LNA-Treatment (L) | >0.9999 | F(1.00,12.00) = 0.00 |
| LNA x Time (LxT) | >0.9999 | F(3.00,12.00) = 0.94 |
| PACF | WT (Untreated) | WT (ntLNA) |
| Week 1 | -0.14 ± 0.14 | -0.046 ± 0.007 |
| Week 2 | -0.054 ± 0.002 | 0.020 ± 0.158 |
| Week 3 | 0.031 ± 0.013 | 0.024 ± 0.031 |
| Week 4 | 0.036 ± 0.007 | 0.040 ± 0.008 |
| Mixed model effects | p-Values | F statistics |
| Time WT (Untreated) | >0.9999 | F(3.00,3.00) = 0.62 |
| Time WT (ntLNA) | >0.9999 | F(3.00,6.00) = 2.99 |
| LNA-Treatment (L) | >0.9999 | F(1.00,3.41) = 1.15 |
| LNA x Time (LxT) | >0.9999 | F(3.00,8.21) = 0.47 |
| SCRF [Hz] | WT (Untreated) | WT (ntLNA) |
| Week 1 | 2.19 ± 0.06 | 2.00 ± 0.26 |
| Week 2 | 0.32 ± 0.08 | 0.66 ± 0.52 |
| Week 3 | 0.12 ± 0.01 | 0.12 ± 0.04 |
| Week 4 | 0.17 ± 0.08 | 0.085 ± 0.004 |
| Mixed model effects | p-Values | F statistics |
| Time WT (Untreated) | 0.2167 | F(3.00,3.18) = 33.30 |
| Time WT (ntLNA) | <0.0001 | F(3.00,5.00) = 447.25 |
| LNA-Treatment (L) | >0.9999 | F(1.00,2.36) = 0.00 |
| LNA x Time (LxT) | >0.9999 | F(3.00,6.33) = 1.58 |
| SCRM | WT (Untreated) | WT (ntLNA) |
| Week 1 | 25.7 ± 1.9 | 27.1 ± 10.9 |
| Week 2 | 310 ± 19 | 216 ± 209 |
| Week 3 | 1098±64 | 979 ± 436 |
| Week 4 | 623 ± 174 | 1112±147 |

| Mixed model effects | p-Values | F statistics |
|---|---|---|
| Time WT (Untreated) | 0.2077 | F(3.00,3.84) = 22.82 |
| Time WT (ntLNA) | 0.0052 | F(3.00,5.00) = 70.91 |
| LNA-Treatment (L) | >0.9999 | F(1.00,4.39) = 0.44 |
| LNA x Time (LxT) | >0.9999 | F(3.00,7.88) = 4.39 |
| RFIT | WT (Untreated) | WT (ntLNA) |
| Week 1 | -0.0024 ± 0.0051 | -0.0039 ± 0.0016 |
| Week 2 | -0.0079 ± 0.0001 | -0.0089 ± 0.0039 |
| Week 3 | -0.0076 ± 0.0005 | -0.0089 ± 0.0036 |
| Week 4 | -0.0098 ± 0.0005 | -0.011 ± 0.001 |
| Mixed model effects | p-Values | F statistics |
| Time WT (Untreated) | >0.9999 | F(3.00,3.86) = 3.99 |
| Time WT (ntLNA) | >0.9999 | F(3.00,4.40) = 3.33 |
| LNA-Treatment (L) | >0.9999 | F(1.00,3.49) = 0.60 |
| LNA x Time (LxT) | >0.9999 | F(3.00,8.36) = 0.02 |
| IBIV | WT (Untreated) | WT (ntLNA) |
| Week 1 | 6257±8294 | 0.58 ± 0.82 |
| Week 2 | 0.26 ± 0.06 | 0.46 ± 0.50 |
| Week 3 | 0.42 ± 0.21 | 0.19 ± 0.03 |
| Week 4 | 2.93 ± 3.37 | 1.73 ± 0.86 |
| Mixed model effects | p-Values | F statistics |
| Time WT (Untreated) | >0.9999 | F(3.00,4.00) = 2.21 |
| Time WT (ntLNA) | >0.9999 | F(3.00,4.86) = 1.20 |
| LNA-Treatment (L) | >0.9999 | F(1.00,3.72) = 0.78 |
| LNA x Time (LxT) | >0.9999 | F(3.00,8.44) = 0.83 |
| IBIM [s] | WT (Untreated) | WT (ntLNA) |
| Week 1 | 116 ± 154 | 2.20 ± 3.12 |
| Week 2 | 11.9 ± 0.2 | 10.4 ± 1.1 |
| Week 3 | 18.9 ± 1.6 | 17.2 ± 0.0 |
| Week 4 | 21.0 ± 4.0 | 22.4 ± 0.1 |
| Mixed model effects | p-Values | F statistics |
| Time WT (Untreated) | 0.0891 | F(3.00,2.15) = 250.95 |
| Time WT (ntLNA) | >0.9999 | F(3.00,4.83) = 0.87 |
| LNA-Treatment (L) | >0.9999 | F(1.00,3.73) = 0.72 |
| LNA x Time (LxT) | >0.9999 | F(3.00,8.40) = 0.79 |
| MBD [s] | WT (Untreated) | WT (ntLNA) |
| Week 1 | 1.27 ± 0.12 | 0.71 ± 1.01 |
| Week 2 | 7.01 ± 0.35 | 8.46 ± 1.68 |
| Week 3 | 9.06 ± 1.15 | 9.95 ± 0.10 |
| Week 4 | 10.8 ± 1.0 | 12.1 ± 0.4 |
| Mixed model effects | p-Values | F statistics |
| Time WT (Untreated) | 0.0429 | F(3.00,4.00) = 48.32 |
| Time WT (ntLNA) | 0.0125 | F(3.00,4.68) = 58.06 |
| LNA-Treatment (L) | >0.9999 | F(1.00,10.00) = 3.13 |
| LNA x Time (LxT) | >0.9999 | F(3.00,10.00) = 1.07 |
| VBD | WT (Untreated) | WT (ntLNA) |
| Week 1 | 0.19 ± 0.24 | 0.36 ± 0.51 |
| Week 2 | 0.097 ± 0.017 | 1.56 ± 1.94 |
| Week 3 | 0.19 ± 0.10 | 0.17 ± 0.03 |
| Week 4 | 0.29 ± 0.08 | 1.95 ± 2.28 |
| Mixed model effects | p-Values | F statistics |
| Time WT (Untreated) | >0.9999 | F(3.00,4.00) = 0.67 |
| Time WT (ntLNA) | >0.9999 | F(3.00,2.94) = 1.60 |
| LNA-Treatment (L) | >0.9999 | F(1.00,9.00) = 2.68 |
| LNA x Time (LxT) | >0.9999 | F(3.00,9.00) = 0.74 |

| | | |
|---|---|---|
| INTRABF [Hz] | WT (Untreated) | WT (ntLNA) |
| Week 1 | 0.20 ± 0.01 | 0.20 ± 0.02 |
| Week 2 | 1.60 ± 0.65 | 0.97 ± 0.88 |
| Week 3 | 3.73 ± 0.99 | 3.98 ± 1.57 |
| Week 4 | 3.88 ± 0.32 | 4.40 ± 0.03 |
| Mixed model effects | p-Values | F statistics |
| Time WT (Untreated) | 0.8139 | F(3.00,2.85) = 16.73 |
| Time WT (ntLNA) | 0.1139 | F(3.00,6.00) = 14.74 |
| LNA-Treatment (L) | >0.9999 | F(1.00,3.30) = 0.01 |
| LNA x Time (LxT) | >0.9999 | F(3.00,7.91) = 0.45 |
| INTERBF [Hz] | WT (Untreated) | WT (ntLNA) |
| Week 1 | 0.064 ± 0.058 | 0.084 ± 0.077 |
| Week 2 | 0.16 ± 0.02 | 0.16 ± 0.10 |
| Week 3 | 0.29 ± 0.07 | 0.26 ± 0.09 |
| Week 4 | 0.34 ± 0.00 | 0.37 ± 0.04 |
| Mixed model effects | p-Values | F statistics |
| Time WT (Untreated) | 0.1412 | F(3.00,3.06) = 49.12 |
| Time WT (ntLNA) | 0.0369 | F(3.00,7.00) = 17.92 |
| LNA-Treatment (L) | >0.9999 | F(1.00,3.57) = 0.20 |
| LNA x Time (LxT) | >0.9999 | F(3.00,8.20) = 0.28 |
| BRT [s] | WT (Untreated) | WT (ntLNA) |
| Week 1 | 0.23 ± 0.21 | 0.15 ± 0.21 |
| Week 2 | 1.36 ± 0.07 | 3.01 ± 1.08 |
| Week 3 | 1.15 ± 0.05 | 1.33 ± 0.18 |
| Week 4 | 1.43 ± 0.05 | 1.19 ± 0.07 |
| Mixed model effects | p-Values | F statistics |
| Time WT (Untreated) | 0.9669 | F(3.00,4.00) = 8.93 |
| Time WT (ntLNA) | 0.0036 | F(3.00,6.00) = 51.52 |
| LNA-Treatment (L) | >0.9999 | F(1.00,10.00) = 4.65 |
| LNA x Time (LxT) | 0.5046 | F(3.00,10.00) = 5.66 |
| BRV [coactivity/s] | WT (Untreated) | WT (ntLNA) |
| Week 1 | 20.2 ± 0.8 | 12.9 ± 14.6 |
| Week 2 | 199 ± 100 | 65.2 ± 76.2 |
| Week 3 | 738 ± 18 | 601 ± 81 |
| Week 4 | 643 ± 72 | 908 ± 1 |
| Mixed model effects | p-Values | F statistics |
| Time WT (Untreated) | 0.0019 | F(3.00,3.84) = 281.50 |
| Time WT (ntLNA) | 0.0380 | F(3.00,2.98) = 129.44 |
| LNA-Treatment (L) | >0.9999 | F(1.00,3.64) = 0.03 |
| LNA x Time (LxT) | 0.0303 | F(3.00,6.43) = 21.67 |
| BDT [s] | WT (Untreated) | WT (ntLNA) |
| Week 1 | 0.23 ± 0.22 | 0.12 ± 0.16 |
| Week 2 | 1.46 ± 0.14 | 1.46 ± 0.10 |
| Week 3 | 4.57 ± 1.65 | 5.74 ± 0.30 |
| Week 4 | 6.00 ± 0.69 | 7.39 ± 0.61 |
| Mixed model effects | p-Values | F statistics |
| Time WT (Untreated) | 0.0029 | F(3.00,4.00) = 191.63 |
| Time WT (ntLNA) | 0.0705 | F(3.00,5.01) = 23.62 |
| LNA-Treatment (L) | >0.9999 | F(1.00,2.43) = 2.30 |
| LNA x Time (LxT) | >0.9999 | F(3.00,7.71) = 1.39 |
| BDV [coactivity/s] | WT (Untreated) | WT (ntLNA) |
| Week 1 | -14.5 ± 15.6 | -13.9 ± 14.9 |
| Week 2 | -207 ± 81 | -103 ± 105 |
| Week 3 | -150 ± 29 | -146 ± 48 |

| | | |
|---|---|---|
| Week 4 | -159 ± 8 | -173 ± 33 |
| Mixed model effects | p-Values | F statistics |
| Time WT (Untreated) | >0.9999 | F(3.00,3.73) = 8.65 |
| Time WT (ntLNA) | 0.2489 | F(3.00,7.00) = 9.28 |
| LNA-Treatment (L) | >0.9999 | F(1.00,4.10) = 0.78 |
| LNA x Time (LxT) | >0.9999 | F(3.00,9.39) = 1.73 |
| SYNC | WT (Untreated) | WT (ntLNA) |
| Week 1 | 0.0020 ± 0.0000 | 0.0020 ± 0.0007 |
| Week 2 | 0.0077 ± 0.0014 | 0.0054 ± 0.0013 |
| Week 3 | 0.0096 ± 0.0006 | 0.0095 ± 0.0013 |
| Week 4 | 0.0084 ± 0.0002 | 0.0093 ± 0.0002 |
| Mixed model effects | p-Values | F statistics |
| Time WT (Untreated) | 0.0568 | F(3.00,3.19) = 78.40 |
| Time WT (ntLNA) | 0.0013 | F(3.00,7.00) = 50.41 |
| LNA-Treatment (L) | >0.9999 | F(1.00,3.36) = 0.56 |
| LNA x Time (LxT) | >0.9999 | F(3.00,8.93) = 3.05 |
| NRF [Hz] | WT (Untreated) | WT (ntLNA) |
| Week 1 | 0.067 ± 0.006 | 0.065 ± 0.002 |
| Week 2 | 0.049 ± 0.008 | 0.053 ± 0.002 |
| Week 3 | 0.038 ± 0.000 | 0.037 ± 0.000 |
| Week 4 | 0.032 ± 0.006 | 0.029 ± 0.000 |
| Mixed model effects | p-Values | F statistics |
| Time WT (Untreated) | 0.0249 | F(3.00,2.33) = 526.08 |
| Time WT (ntLNA) | 0.0712 | F(3.00,6.00) = 17.62 |
| LNA-Treatment (L) | >0.9999 | F(1.00,10.00) = 0.04 |
| LNA x Time (LxT) | >0.9999 | F(3.00,10.00) = 0.70 |
| NRM | WT (Untreated) | WT (ntLNA) |
| Week 1 | 91.2 ± 33.0 | 70.6 ± 17.4 |
| Week 2 | 362 ± 10 | 486 ± 186 |
| Week 3 | 461 ± 114 | 566 ± 90 |
| Week 4 | 414 ± 102 | 509 ± 27 |
| Mixed model effects | p-Values | F statistics |
| Time WT (Untreated) | 0.6910 | F(3.00,3.00) = 17.15 |
| Time WT (ntLNA) | >0.9999 | F(3.00,2.31) = 17.18 |
| LNA-Treatment (L) | >0.9999 | F(1.00,9.00) = 1.75 |
| LNA x Time (LxT) | >0.9999 | F(3.00,9.00) = 0.77 |
| NRFIT | WT (Untreated) | WT (ntLNA) |
| Week 1 | -0.015 ± 0.012 | -0.025 ± 0.032 |
| Week 2 | -0.0018 ± 0.0000 | -0.0027 ± 0.0011 |
| Week 3 | -0.0017 ± 0.0002 | -0.0017 ± 0.0001 |
| Week 4 | -0.0015 ± 0.0001 | -0.0016 ± 0.0001 |
| Mixed model effects | p-Values | F statistics |
| Time WT (Untreated) | >0.9999 | F(3.00,4.00) = 0.99 |
| Time WT (ntLNA) | >0.9999 | F(3.00,5.00) = 2.90 |
| LNA-Treatment (L) | >0.9999 | F(1.00,9.00) = 0.23 |
| LNA x Time (LxT) | >0.9999 | F(3.00,9.00) = 0.17 |

**Supplemental table 12.** Statistical comparison of untreated and non-targeted LNA (ntLNA)-treated WT cultures. Reported are Mean ± SD of all features for each week and treatment condition and the results of linear mixed effect models assessing LNA-treatment (L) and LNA-treatment-by-time interaction (LxT). p-Values were adjusted for the number of comparisons (Bonferroni correction). Only one of the metrics showed a significant difference, indicating that non-specific effects of the LNA-treatment did not significantly alter the electrophysiological phenotype. Sample size for these analyses were 3 untreated and 3 non-targeted LNA-treated WT cultures.

| | | |
|---|---|---|
| ISIM [s] | A53T (Untreated) | A53T (ntLNA) |
| Week 1 | 191 ± 45 | 196 ± 0 |
| Week 2 | 42.3 ± 21.4 | 50.5 ± 8.0 |
| Week 3 | 86.9 ± 63.2 | 17.8 ± 1.7 |
| Week 4 | 216 ± 189 | 23.7 ± 9.5 |
| Mixed model effects | p-Values | F statistics |
| Time A53T (Untreated) | 0.0001 | F(3.00,5.00) = 321.65 |
| Time A53T (ntLNA) | >0.9999 | F(3.00,8.00) = 1.95 |
| LNA-Treatment (L) | >0.9999 | F(1.00,13.00) = 2.99 |
| LNA x Time (LxT) | >0.9999 | F(3.00,13.00) = 1.69 |
| ISIV | A53T (Untreated) | A53T (ntLNA) |
| Week 1 | 2080±948 | 2416±323 |
| Week 2 | 138 ± 20 | 290 ± 83 |
| Week 3 | 819 ± 718 | 58.8 ± 29.9 |
| Week 4 | 3351±2857 | 116 ± 94 |
| Mixed model effects | p-Values | F statistics |
| Time A53T (Untreated) | 0.0007 | F(3.00,6.00) = 91.29 |
| Time A53T (ntLNA) | >0.9999 | F(3.00,7.00) = 1.93 |
| LNA-Treatment (L) | >0.9999 | F(1.00,13.00) = 2.60 |
| LNA x Time (LxT) | >0.9999 | F(3.00,13.00) = 2.27 |
| ISICV | A53T (Untreated) | A53T (ntLNA) |
| Week 1 | 1.19 ± 0.10 | 1.13 ± 0.00 |
| Week 2 | 2.18 ± 0.68 | 1.81 ± 0.32 |
| Week 3 | 1.85 ± 0.26 | 2.40 ± 0.02 |
| Week 4 | 1.51 ± 0.54 | 2.15 ± 0.17 |
| Mixed model effects | p-Values | F statistics |
| Time A53T (Untreated) | 0.2888 | F(3.00,5.00) = 12.65 |
| Time A53T (ntLNA) | >0.9999 | F(3.00,8.00) = 2.64 |
| LNA-Treatment (L) | >0.9999 | F(1.00,13.00) = 1.21 |
| LNA x Time (LxT) | >0.9999 | F(3.00,13.00) = 2.14 |
| PACF | A53T (Untreated) | A53T (ntLNA) |
| Week 1 | -0.021 ± 0.003 | -0.055 ± 0.056 |
| Week 2 | 0.015 ± 0.009 | -0.086 ± 0.003 |
| Week 3 | -0.016 ± 0.000 | 0.019 ± 0.012 |
| Week 4 | 0.0066 ± 0.0763 | 0.017 ± 0.005 |
| Mixed model effects | p-Values | F statistics |
| Time A53T (Untreated) | >0.9999 | F(3.00,3.99) = 4.51 |
| Time A53T (ntLNA) | >0.9999 | F(3.00,5.00) = 0.28 |
| LNA-Treatment (L) | >0.9999 | F(1.00,10.00) = 1.25 |
| LNA x Time (LxT) | >0.9999 | F(3.00,10.00) = 2.00 |
| SCRF [Hz] | A53T (Untreated) | A53T (ntLNA) |
| Week 1 | 1.61 ± 0.27 | 1.76 ± 0.08 |
| Week 2 | 0.45 ± 0.02 | 0.49 ± 0.01 |
| Week 3 | 1.15 ± 0.12 | 0.18 ± 0.05 |
| Week 4 | 1.25 ± 0.99 | 0.31 ± 0.11 |
| Mixed model effects | p-Values | F statistics |
| Time A53T (Untreated) | 0.0003 | F(3.00,5.00) = 237.34 |
| Time A53T (ntLNA) | >0.9999 | F(3.00,6.00) = 1.53 |
| LNA-Treatment (L) | >0.9999 | F(1.00,11.00) = 4.22 |
| LNA x Time (LxT) | >0.9999 | F(3.00,11.00) = 2.21 |
| SCRM | A53T (Untreated) | A53T (ntLNA) |
| Week 1 | 39.6 ± 10.5 | 32.1 ± 0.3 |
| Week 2 | 375 ± 262 | 229 ± 92 |
| Week 3 | 159 ± 32 | 415 ± 290 |
| Week 4 | 209 ± 225 | 403 ± 207 |

| | | |
|---|---|---|
| Mixed model effects | p-Values | F statistics |
| Time A53T (Untreated) | >0.9999 | F(3.00,6.00) = 1.89 |
| Time A53T (ntLNA) | >0.9999 | F(3.00,7.00) = 1.68 |
| LNA-Treatment (L) | >0.9999 | F(1.00,13.00) = 0.76 |
| LNA x Time (LxT) | >0.9999 | F(3.00,13.00) = 1.26 |
| RFIT | A53T (Untreated) | A53T (ntLNA) |
| Week 1 | -0.0045 ± 0.0016 | -0.0047 ± 0.0022 |
| Week 2 | -0.0074 ± 0.0001 | -0.0084 ± 0.0011 |
| Week 3 | -0.0044 ± 0.0024 | -0.0066 ± 0.0002 |
| Week 4 | -0.011 ± 0.001 | -0.011 ± 0.003 |
| Mixed model effects | p-Values | F statistics |
| Time A53T (Untreated) | 0.8310 | F(3.00,7.00) = 5.80 |
| Time A53T (ntLNA) | 0.4951 | F(3.00,6.00) = 8.14 |
| LNA-Treatment (L) | >0.9999 | F(1.00,13.00) = 1.00 |
| LNA x Time (LxT) | >0.9999 | F(3.00,13.00) = 0.58 |
| IBIV | A53T (Untreated) | A53T (ntLNA) |
| Week 1 | 31.5 ± 9.4 | 14.8 ± 9.9 |
| Week 2 | 0.32 ± 0.26 | 0.21 ± 0.02 |
| Week 3 | 16.3 ± 16.9 | 0.082 ± 0.034 |
| Week 4 | 6.92 ± 9.66 | 5.40 ± 4.00 |
| Mixed model effects | p-Values | F statistics |
| Time A53T (Untreated) | >0.9999 | F(3.00,5.00) = 2.94 |
| Time A53T (ntLNA) | >0.9999 | F(3.00,4.10) = 3.41 |
| LNA-Treatment (L) | >0.9999 | F(1.00,11.00) = 3.91 |
| LNA x Time (LxT) | >0.9999 | F(3.00,11.00) = 1.09 |
| IBIM [s] | A53T (Untreated) | A53T (ntLNA) |
| Week 1 | 16.4 ± 5.6 | 6.53 ± 0.97 |
| Week 2 | 15.1 ± 3.5 | 13.7 ± 2.4 |
| Week 3 | 10.4 ± 0.0 | 12.3 ± 1.0 |
| Week 4 | 11.7 ± 2.9 | 10.8 ± 0.4 |
| Mixed model effects | p-Values | F statistics |
| Time A53T (Untreated) | >0.9999 | F(3.00,3.22) = 9.50 |
| Time A53T (ntLNA) | >0.9999 | F(3.00,6.00) = 1.18 |
| LNA-Treatment (L) | >0.9999 | F(1.00,12.00) = 3.50 |
| LNA x Time (LxT) | >0.9999 | F(3.00,12.00) = 3.30 |
| MBD [s] | A53T (Untreated) | A53T (ntLNA) |
| Week 1 | 5.53 ± 2.29 | 4.07 ± 0.14 |
| Week 2 | 7.01 ± 0.54 | 7.48 ± 0.05 |
| Week 3 | 6.51 ± 0.91 | 6.28 ± 1.12 |
| Week 4 | 6.01 ± 0.59 | 8.89 ± 2.30 |
| Mixed model effects | p-Values | F statistics |
| Time A53T (Untreated) | >0.9999 | F(3.00,4.09) = 7.16 |
| Time A53T (ntLNA) | >0.9999 | F(3.00,5.18) = 0.67 |
| LNA-Treatment (L) | >0.9999 | F(1.00,3.52) = 0.40 |
| LNA x Time (LxT) | >0.9999 | F(3.00,9.41) = 2.15 |
| VBD | A53T (Untreated) | A53T (ntLNA) |
| Week 1 | 7.46 ± 3.91 | 6.49 ± 2.91 |
| Week 2 | 0.22 ± 0.13 | 0.59 ± 0.03 |
| Week 3 | 4.41 ± 5.62 | 0.10 ± 0.03 |
| Week 4 | 0.48 ± 0.45 | 6.74 ± 9.20 |
| Mixed model effects | p-Values | F statistics |
| Time A53T (Untreated) | >0.9999 | F(3.00,4.00) = 1.13 |
| Time A53T (ntLNA) | >0.9999 | F(3.00,6.00) = 3.85 |
| LNA-Treatment (L) | >0.9999 | F(1.00,3.01) = 0.05 |
| LNA x Time (LxT) | >0.9999 | F(3.00,8.82) = 1.61 |

| | | |
|---|---|---|
| INTRABF [Hz] | A53T (Untreated) | A53T (ntLNA) |
| Week 1 | 0.24 ± 0.04 | 0.22 ± 0.04 |
| Week 2 | 1.95 ± 1.22 | 1.19 ± 0.66 |
| Week 3 | 1.56 ± 0.59 | 2.54 ± 0.18 |
| Week 4 | 1.28 ± 1.05 | 2.09 ± 0.74 |
| Mixed model effects | p-Values | F statistics |
| Time A53T (Untreated) | 0.2458 | F(3.00,6.00) = 10.89 |
| Time A53T (ntLNA) | >0.9999 | F(3.00,8.00) = 2.18 |
| LNA-Treatment (L) | >0.9999 | F(1.00,14.00) = 0.67 |
| LNA x Time (LxT) | >0.9999 | F(3.00,14.00) = 1.68 |
| INTERBF [Hz] | A53T (Untreated) | A53T (ntLNA) |
| Week 1 | 0.11 ± 0.00 | 0.13 ± 0.03 |
| Week 2 | 0.18 ± 0.01 | 0.18 ± 0.06 |
| Week 3 | 0.23 ± 0.05 | 0.31 ± 0.02 |
| Week 4 | 0.22 ± 0.19 | 0.38 ± 0.09 |
| Mixed model effects | p-Values | F statistics |
| Time A53T (Untreated) | 0.1610 | F(3.00,6.00) = 12.89 |
| Time A53T (ntLNA) | >0.9999 | F(3.00,3.91) = 1.05 |
| LNA-Treatment (L) | >0.9999 | F(1.00,2.71) = 1.70 |
| LNA x Time (LxT) | >0.9999 | F(3.00,7.75) = 1.18 |
| BRT [s] | A53T (Untreated) | A53T (ntLNA) |
| Week 1 | 2.13 ± 0.07 | 0.86 ± 0.25 |
| Week 2 | 1.96 ± 0.40 | 2.22 ± 0.03 |
| Week 3 | 1.82 ± 0.04 | 1.40 ± 0.04 |
| Week 4 | 1.38 ± 1.09 | 3.17 ± 0.56 |
| Mixed model effects | p-Values | F statistics |
| Time A53T (Untreated) | 0.4497 | F(3.00,2.71) = 29.28 |
| Time A53T (ntLNA) | >0.9999 | F(3.00,3.85) = 1.42 |
| LNA-Treatment (L) | >0.9999 | F(1.00,3.22) = 0.00 |
| LNA x Time (LxT) | 0.2966 | F(3.00,6.49) = 9.32 |
| BRV [coactivity/s] | A53T (Untreated) | A53T (ntLNA) |
| Week 1 | 11.8 ± 1.7 | 27.5 ± 2.1 |
| Week 2 | 50.4 ± 15.0 | 57.6 ± 2.1 |
| Week 3 | 48.9 ± 6.3 | 196 ± 158 |
| Week 4 | 37.5 ± 13.0 | 145 ± 2 |
| Mixed model effects | p-Values | F statistics |
| Time A53T (Untreated) | >0.9999 | F(3.00,3.67) = 1.46 |
| Time A53T (ntLNA) | >0.9999 | F(3.00,4.00) = 5.85 |
| LNA-Treatment (L) | >0.9999 | F(1.00,2.24) = 3.43 |
| LNA x Time (LxT) | >0.9999 | F(3.00,6.43) = 0.91 |
| BDT [s] | A53T (Untreated) | A53T (ntLNA) |
| Week 1 | 1.07 ± 0.52 | 0.78 ± 0.33 |
| Week 2 | 1.65 ± 0.11 | 1.56 ± 0.12 |
| Week 3 | 1.14 ± 0.01 | 1.50 ± 0.29 |
| Week 4 | 0.98 ± 0.62 | 1.52 ± 0.04 |
| Mixed model effects | p-Values | F statistics |
| Time A53T (Untreated) | 0.9969 | F(3.00,6.00) = 5.97 |
| Time A53T (ntLNA) | >0.9999 | F(3.00,7.00) = 1.44 |
| LNA-Treatment (L) | >0.9999 | F(1.00,13.00) = 0.66 |
| LNA x Time (LxT) | >0.9999 | F(3.00,13.00) = 1.50 |
| BDV [coactivity/s] | A53T (Untreated) | A53T (ntLNA) |
| Week 1 | -20.7 ± 7.7 | -22.5 ± 11.2 |
| Week 2 | -99.6 ± 53.5 | -86.7 ± 2.8 |
| Week 3 | -71.0 ± 15.7 | -187 ± 144 |

| | | |
|---|---|---|
| Week 4 | -34.4 ± 5.6 | -269 ± 10 |
| Mixed model effects | p-Values | F statistics |
| Time A53T (Untreated) | >0.9999 | F(3.00,6.00) = 4.14 |
| Time A53T (ntLNA) | >0.9999 | F(3.00,6.00) = 3.53 |
| LNA-Treatment (L) | 0.3951 | F(1.00,12.00) = 8.65 |
| LNA x Time (LxT) | >0.9999 | F(3.00,12.00) = 3.73 |
| SYNC | A53T (Untreated) | A53T (ntLNA) |
| Week 1 | 0.0035 ± 0.0001 | 0.0027 ± 0.0003 |
| Week 2 | 0.0074 ± 0.0018 | 0.0060 ± 0.0006 |
| Week 3 | 0.0064 ± 0.0000 | 0.0082 ± 0.0002 |
| Week 4 | 0.0045 ± 0.0025 | 0.0063 ± 0.0005 |
| Mixed model effects | p-Values | F statistics |
| Time A53T (Untreated) | 0.0017 | F(3.00,6.00) = 67.47 |
| Time A53T (ntLNA) | >0.9999 | F(3.00,6.00) = 2.46 |
| LNA-Treatment (L) | >0.9999 | F(1.00,12.00) = 0.40 |
| LNA x Time (LxT) | >0.9999 | F(3.00,12.00) = 2.12 |
| NRF [Hz] | A53T (Untreated) | A53T (ntLNA) |
| Week 1 | 0.044 ± 0.005 | 0.043 ± 0.001 |
| Week 2 | 0.046 ± 0.008 | 0.048 ± 0.005 |
| Week 3 | 0.046 ± 0.015 | 0.049 ± 0.009 |
| Week 4 | 0.049 ± 0.012 | 0.049 ± 0.011 |
| Mixed model effects | p-Values | F statistics |
| Time A53T (Untreated) | >0.9999 | F(3.00,4.00) = 0.30 |
| Time A53T (ntLNA) | >0.9999 | F(3.00,6.00) = 0.25 |
| LNA-Treatment (L) | >0.9999 | F(1.00,3.95) = 0.03 |
| LNA x Time (LxT) | >0.9999 | F(3.00,10.11) = 0.24 |
| NRM | A53T (Untreated) | A53T (ntLNA) |
| Week 1 | 269 ± 36 | 230 ± 68 |
| Week 2 | 387 ± 17 | 588 ± 76 |
| Week 3 | 129 ± 11 | 386 ± 208 |
| Week 4 | 275 ± 206 | 396 ± 263 |
| Mixed model effects | p-Values | F statistics |
| Time A53T (Untreated) | >0.9999 | F(3.00,7.00) = 2.55 |
| Time A53T (ntLNA) | >0.9999 | F(3.00,6.00) = 1.53 |
| LNA-Treatment (L) | >0.9999 | F(1.00,13.00) = 4.53 |
| LNA x Time (LxT) | >0.9999 | F(3.00,13.00) = 1.11 |
| NRFIT | A53T (Untreated) | A53T (ntLNA) |
| Week 1 | -0.010 ± 0.003 | -0.018 ± 0.001 |
| Week 2 | -0.0028 ± 0.0004 | -0.0031 ± 0.0001 |
| Week 3 | -0.0017 ± 0.0001 | -0.0021 ± 0.0002 |
| Week 4 | -0.0020 ± 0.0000 | -0.0025 ± 0.0001 |
| Mixed model effects | p-Values | F statistics |
| Time A53T (Untreated) | <0.0001 | F(3.00,6.00) = 594.95 |
| Time A53T (ntLNA) | 0.1918 | F(3.00,4.41) = 18.49 |
| LNA-Treatment (L) | 0.5481 | F(1.00,3.28) = 20.11 |
| LNA x Time (LxT) | 0.0220 | F(3.00,8.94) = 15.52 |

**Supplemental table 13.** Statistical comparison of untreated and non-targeted LNA-treated A53T cultures. Reported are Mean ± SD of all features for each week and treatment condition and the results of linear mixed effect models assessing LNA-treatment (L) and LNA-treatment-by-time interaction (LxT). p-Values were adjusted for the number of comparisons (Bonferroni correction). Only one of the investigated metrics (PACF) showed a significant difference, indicating that non-specific effects of the LNA-treatment did not significantly alter the electrophysiological phenotype. Sample size for these analyses were 3 untreated and 3 non-targeted LNA-treated A53T cultures.

| ISIM [s] | WT (Untreated) | WT (LNA) |
|---|---|---|
| Week 1 | 232 ± 23 | 266 ± 15 |
| Week 2 | 48.8 ± 8.2 | 31.4 ± 5.6 |
| Week 3 | 41.1 ± 8.6 | 16.3 ± 5.8 |
| Week 4 | 155 ± 111 | 16.5 ± 4.0 |
| Mixed model effects | p-Values | F statistics |
| Time WT (Untreated) | <0.0001 | F(3.00,12.00) = 778.86 |
| Time WT (LNA) | 0.0937 | F(3.00,11.00) = 8.79 |
| LNA-Treatment (L) | 0.7308 | F(1.00,23.00) = 5.95 |
| LNA x Time (LxT) | 0.1039 | F(3.00,23.00) = 6.12 |
| ISIV | WT (Untreated) | WT (LNA) |
| Week 1 | 2347±304 | 3609±896 |
| Week 2 | 224 ± 75 | 151 ± 61 |
| Week 3 | 236 ± 109 | 45.9 ± 13.7 |
| Week 4 | 2072±1888 | 73.7 ± 29.4 |
| Mixed model effects | p-Values | F statistics |
| Time WT (Untreated) | <0.0001 | F(3.00,7.06) = 138.89 |
| Time WT (LNA) | >0.9999 | F(3.00,10.00) = 4.15 |
| LNA-Treatment (L) | >0.9999 | F(1.00,23.00) = 0.71 |
| LNA x Time (LxT) | 0.1692 | F(3.00,23.00) = 5.51 |
| ISICV | WT (Untreated) | WT (LNA) |
| Week 1 | 1.04 ± 0.01 | 1.04 ± 0.05 |
| Week 2 | 1.56 ± 0.01 | 1.90 ± 0.45 |
| Week 3 | 1.67 ± 0.41 | 3.32 ± 0.51 |
| Week 4 | 1.34 ± 0.24 | 3.63 ± 0.21 |
| Mixed model effects | p-Values | F statistics |
| Time WT (Untreated) | <0.0001 | F(3.00,16.00) = 59.50 |
| Time WT (LNA) | 0.3898 | F(3.00,7.87) = 7.16 |
| LNA-Treatment (L) | <0.0001 | F(1.00,27.00) = 96.28 |
| LNA x Time (LxT) | <0.0001 | F(3.00,27.00) = 23.91 |
| PACF | WT (Untreated) | WT (LNA) |
| Week 1 | -0.058 ± 0.036 | -0.069 ± 0.042 |
| Week 2 | -0.066 ± 0.011 | -0.051 ± 0.036 |
| Week 3 | -0.019 ± 0.040 | 0.028 ± 0.018 |
| Week 4 | 0.059 ± 0.134 | 0.038 ± 0.006 |
| Mixed model effects | p-Values | F statistics |
| Time WT (Untreated) | 0.0059 | F(3.00,11.10) = 17.00 |
| Time WT (LNA) | >0.9999 | F(3.00,11.00) = 2.24 |
| LNA-Treatment (L) | >0.9999 | F(1.00,25.00) = 0.17 |
| LNA x Time (LxT) | >0.9999 | F(3.00,25.00) = 0.65 |
| SCRF [Hz] | WT (Untreated) | WT (LNA) |
| Week 1 | 1.99 ± 0.21 | 2.01 ± 0.24 |
| Week 2 | 0.44 ± 0.06 | 0.32 ± 0.07 |
| Week 3 | 0.39 ± 0.20 | 0.12 ± 0.03 |
| Week 4 | 1.01 ± 0.65 | 0.096 ± 0.019 |
| Mixed model effects | p-Values | F statistics |
| Time WT (Untreated) | <0.0001 | F(3.00,7.52) = 581.65 |
| Time WT (LNA) | 0.0114 | F(3.00,8.36) = 20.12 |
| LNA-Treatment (L) | 0.7927 | F(1.00,7.44) = 7.86 |
| LNA x Time (LxT) | 0.1764 | F(3.00,17.76) = 5.92 |
| SCRM | WT (Untreated) | WT (LNA) |
| Week 1 | 26.6 ± 0.5 | 25.6 ± 1.6 |
| Week 2 | 169 ± 38 | 246 ± 121 |
| Week 3 | 336 ± 223 | 917 ± 356 |
| Week 4 | 156 ± 113 | 868 ± 312 |

| | | |
|---|---|---|
| Mixed model effects | p-Values | F statistics |
| Time WT (Untreated) | 0.0122 | F(3.00,10.78) = 14.82 |
| Time WT (LNA) | 0.7118 | F(3.00,7.98) = 5.68 |
| LNA-Treatment (L) | 0.1403 | F(1.00,6.73) = 17.69 |
| LNA x Time (LxT) | 0.1118 | F(3.00,18.74) = 6.44 |
| RFIT | WT (Untreated) | WT (LNA) |
| Week 1 | -0.0034 ± 0.0015 | -0.0032 ± 0.0035 |
| Week 2 | -0.011 ± 0.002 | -0.0085 ± 0.0020 |
| Week 3 | -0.010 ± 0.004 | -0.0073 ± 0.0008 |
| Week 4 | -0.0079 ± 0.0033 | -0.0098 ± 0.0004 |
| Mixed model effects | p-Values | F statistics |
| Time WT (Untreated) | 0.1482 | F(3.00,11.16) = 7.70 |
| Time WT (LNA) | 0.9236 | F(3.00,7.17) = 5.46 |
| LNA-Treatment (L) | >0.9999 | F(1.00,6.41) = 0.69 |
| LNA x Time (LxT) | >0.9999 | F(3.00,18.36) = 1.32 |
| IBIV | WT (Untreated) | WT (LNA) |
| Week 1 | 7.50 ± 12.98 | 777 ± 1553 |
| Week 2 | 1.10 ± 1.06 | 0.50 ± 0.41 |
| Week 3 | 0.35 ± 0.38 | 0.25 ± 0.11 |
| Week 4 | 0.22 ± 0.22 | 2.33 ± 2.12 |
| Mixed model effects | p-Values | F statistics |
| Time WT (Untreated) | NaN | F(3.00,0.00) = 1.10 |
| Time WT (LNA) | >0.9999 | F(3.00,10.00) = 1.11 |
| LNA-Treatment (L) | NaN | F(1.00,0.00) = 0.84 |
| LNA x Time (LxT) | >0.9999 | F(3.00,3.47) = 0.84 |
| IBIM [s] | WT (Untreated) | WT (LNA) |
| Week 1 | 7.34 ± 9.28 | 31.2 ± 43.2 |
| Week 2 | 9.71 ± 0.16 | 11.1 ± 1.1 |
| Week 3 | 10.4 ± 0.4 | 17.6 ± 0.6 |
| Week 4 | 7.62 ± 1.56 | 22.9 ± 0.8 |
| Mixed model effects | p-Values | F statistics |
| Time WT (Untreated) | >0.9999 | F(3.00,10.12) = 0.52 |
| Time WT (LNA) | >0.9999 | F(3.00,7.04) = 0.26 |
| LNA-Treatment (L) | >0.9999 | F(1.00,6.02) = 2.71 |
| LNA x Time (LxT) | >0.9999 | F(3.00,16.48) = 0.51 |
| MBD [s] | WT (Untreated) | WT (LNA) |
| Week 1 | 1.44 ± 1.96 | 0.80 ± 0.73 |
| Week 2 | 9.24 ± 0.76 | 7.07 ± 0.31 |
| Week 3 | 7.83 ± 0.79 | 9.77 ± 0.59 |
| Week 4 | 7.64 ± 3.59 | 11.4 ± 1.0 |
| Mixed model effects | p-Values | F statistics |
| Time WT (Untreated) | <0.0001 | F(3.00,10.34) = 244.99 |
| Time WT (LNA) | 0.0326 | F(3.00,12.00) = 10.76 |
| LNA-Treatment (L) | >0.9999 | F(1.00,25.00) = 1.82 |
| LNA x Time (LxT) | 0.1077 | F(3.00,25.00) = 5.93 |
| VBD | WT (Untreated) | WT (LNA) |
| Week 1 | 0.24 ± 0.42 | 0.26 ± 0.32 |
| Week 2 | 0.48 ± 0.27 | 0.16 ± 0.08 |
| Week 3 | 0.41 ± 0.32 | 0.18 ± 0.06 |
| Week 4 | 0.41 ± 0.22 | 0.30 ± 0.06 |
| Mixed model effects | p-Values | F statistics |
| Time WT (Untreated) | >0.9999 | F(3.00,8.43) = 0.93 |
| Time WT (LNA) | >0.9999 | F(3.00,8.00) = 0.32 |
| LNA-Treatment (L) | >0.9999 | F(1.00,6.90) = 2.59 |
| LNA x Time (LxT) | >0.9999 | F(3.00,16.32) = 0.64 |

| | | |
|---|---|---|
| INTRABF [Hz] | WT (Untreated) | WT (LNA) |
| Week 1 | 0.085 ± 0.099 | 0.20 ± 0.01 |
| Week 2 | 0.80 ± 0.04 | 1.35 ± 0.72 |
| Week 3 | 1.27 ± 0.69 | 3.83 ± 1.06 |
| Week 4 | 0.65 ± 0.34 | 4.14 ± 0.35 |
| Mixed model effects | p-Values | F statistics |
| Time WT (Untreated) | <0.0001 | F(3.00,14.00) = 31.68 |
| Time WT (LNA) | 0.2867 | F(3.00,9.00) = 7.25 |
| LNA-Treatment (L) | 0.0025 | F(1.00,6.87) = 69.67 |
| LNA x Time (LxT) | 0.0006 | F(3.00,19.75) = 15.84 |
| INTERBF [Hz] | WT (Untreated) | WT (LNA) |
| Week 1 | 0.049 ± 0.058 | 0.074 ± 0.062 |
| Week 2 | 0.17 ± 0.04 | 0.16 ± 0.05 |
| Week 3 | 0.16 ± 0.05 | 0.28 ± 0.07 |
| Week 4 | 0.16 ± 0.01 | 0.34 ± 0.00 |
| Mixed model effects | p-Values | F statistics |
| Time WT (Untreated) | 0.0002 | F(3.00,9.98) = 40.95 |
| Time WT (LNA) | 0.4307 | F(3.00,10.00) = 5.96 |
| LNA-Treatment (L) | 0.2931 | F(1.00,6.81) = 12.96 |
| LNA x Time (LxT) | 0.0568 | F(3.00,17.45) = 7.66 |
| BRT [s] | WT (Untreated) | WT (LNA) |
| Week 1 | 0.28 ± 0.36 | 0.20 ± 0.18 |
| Week 2 | 4.07 ± 0.62 | 2.15 ± 0.99 |
| Week 3 | 2.79 ± 0.18 | 1.22 ± 0.14 |
| Week 4 | 2.98 ± 0.33 | 1.31 ± 0.15 |
| Mixed model effects | p-Values | F statistics |
| Time WT (Untreated) | 0.0120 | F(3.00,15.00) = 11.39 |
| Time WT (LNA) | 0.0003 | F(3.00,8.00) = 58.22 |
| LNA-Treatment (L) | <0.0001 | F(1.00,25.00) = 56.52 |
| LNA x Time (LxT) | 0.0848 | F(3.00,25.00) = 6.22 |
| BRV [coactivity/s] | WT (Untreated) | WT (LNA) |
| Week 1 | 14.9 ± 17.2 | 13.2 ± 11.8 |
| Week 2 | 23.2 ± 6.5 | 146 ± 109 |
| Week 3 | 56.4 ± 53.1 | 591 ± 194 |
| Week 4 | 16.5 ± 5.9 | 775 ± 159 |
| Mixed model effects | p-Values | F statistics |
| Time WT (Untreated) | <0.0001 | F(3.00,16.00) = 37.08 |
| Time WT (LNA) | >0.9999 | F(3.00,11.00) = 1.69 |
| LNA-Treatment (L) | <0.0001 | F(1.00,27.00) = 100.38 |
| LNA x Time (LxT) | <0.0001 | F(3.00,27.00) = 24.39 |
| BDT [s] | WT (Untreated) | WT (LNA) |
| Week 1 | 0.086 ± 0.148 | 0.18 ± 0.19 |
| Week 2 | 1.89 ± 0.06 | 1.46 ± 0.11 |
| Week 3 | 2.45 ± 0.09 | 5.60 ± 0.55 |
| Week 4 | 2.32 ± 0.17 | 6.70 ± 0.96 |
| Mixed model effects | p-Values | F statistics |
| Time WT (Untreated) | <0.0001 | F(3.00,10.93) = 180.33 |
| Time WT (LNA) | <0.0001 | F(3.00,6.89) = 389.22 |
| LNA-Treatment (L) | 0.0006 | F(1.00,7.21) = 100.41 |
| LNA x Time (LxT) | <0.0001 | F(3.00,18.09) = 67.28 |
| BDV [coactivity/s] | WT (Untreated) | WT (LNA) |
| Week 1 | -17.8 ± 21.8 | -14.2 ± 13.6 |
| Week 2 | -49.3 ± 8.0 | -165 ± 97 |
| Week 3 | -50.4 ± 34.3 | -149 ± 32 |

| | | |
|---|---|---|
| Week 4 | -14.9 ± 1.1 | -156 ± 8 |
| Mixed model effects | p-Values | F statistics |
| Time WT (Untreated) | 0.0315 | F(3.00,11.72) = 11.04 |
| Time WT (LNA) | >0.9999 | F(3.00,7.62) = 2.98 |
| LNA-Treatment (L) | 0.0295 | F(1.00,7.57) = 27.60 |
| LNA x Time (LxT) | 0.2447 | F(3.00,19.69) = 5.30 |
| SYNC | WT (Untreated) | WT (LNA) |
| Week 1 | 0.0022 ± 0.0001 | 0.0020 ± 0.0001 |
| Week 2 | 0.0052 ± 0.0003 | 0.0068 ± 0.0017 |
| Week 3 | 0.0050 ± 0.0015 | 0.0095 ± 0.0008 |
| Week 4 | 0.0039 ± 0.0011 | 0.0089 ± 0.0005 |
| Mixed model effects | p-Values | F statistics |
| Time WT (Untreated) | <0.0001 | F(3.00,14.00) = 45.44 |
| Time WT (LNA) | 0.1311 | F(3.00,9.00) = 9.26 |
| LNA-Treatment (L) | 0.0147 | F(1.00,5.79) = 50.42 |
| LNA x Time (LxT) | 0.0019 | F(3.00,18.50) = 13.68 |
| NRF [Hz] | WT (Untreated) | WT (LNA) |
| Week 1 | 0.062 ± 0.005 | 0.068 ± 0.005 |
| Week 2 | 0.050 ± 0.005 | 0.053 ± 0.002 |
| Week 3 | 0.051 ± 0.006 | 0.036 ± 0.002 |
| Week 4 | 0.060 ± 0.001 | 0.029 ± 0.001 |
| Mixed model effects | p-Values | F statistics |
| Time WT (Untreated) | <0.0001 | F(3.00,14.00) = 109.31 |
| Time WT (LNA) | 0.0167 | F(3.00,7.03) = 22.98 |
| LNA-Treatment (L) | 0.1102 | F(1.00,7.38) = 17.90 |
| LNA x Time (LxT) | <0.0001 | F(3.00,16.65) = 50.14 |
| NRM | WT (Untreated) | WT (LNA) |
| Week 1 | 184 ± 100 | 80.9 ± 24.6 |
| Week 2 | 448 ± 34 | 359 ± 9 |
| Week 3 | 627 ± 153 | 503 ± 108 |
| Week 4 | 685 ± 309 | 502 ± 24 |
| Mixed model effects | p-Values | F statistics |
| Time WT (Untreated) | 0.0002 | F(3.00,6.38) = 117.40 |
| Time WT (LNA) | 0.3986 | F(3.00,8.37) = 6.81 |
| LNA-Treatment (L) | >0.9999 | F(1.00,6.58) = 4.46 |
| LNA x Time (LxT) | >0.9999 | F(3.00,16.61) = 0.17 |
| NRFIT | WT (Untreated) | WT (LNA) |
| Week 1 | -0.032 ± 0.028 | -0.020 ± 0.021 |
| Week 2 | -0.0042 ± 0.0012 | -0.0019 ± 0.0001 |
| Week 3 | -0.0047 ± 0.0030 | -0.0017 ± 0.0001 |
| Week 4 | -0.0070 ± 0.0010 | -0.0015 ± 0.0001 |
| Mixed model effects | p-Values | F statistics |
| Time WT (Untreated) | >0.9999 | F(3.00,9.93) = 3.05 |
| Time WT (LNA) | >0.9999 | F(3.00,10.00) = 3.46 |
| LNA-Treatment (L) | >0.9999 | F(1.00,22.00) = 1.76 |
| LNA x Time (LxT) | >0.9999 | F(3.00,22.00) = 0.25 |

**Supplemental table 14.** Complementary data related to Figure 5. Reported are Mean ± SD of all features for each week and treatment condition and the results of linear mixed-effect models assessing LNA-treatment (L) and LNA-treatment-by-time interaction (LxT). p-Values were adjusted for the number of comparisons (Bonferroni correction). Sample size for these analyses were 6 untreated/non-targeted LNA-treated WT cultures and 4 LNA-treated WT cultures.

| ISIM [s] | A53T (Untreated) | A53T (LNA) |
|---|---|---|
| Week 1 | 250 ± 14 | 196 ± 6 |
| Week 2 | 68.7 ± 24.4 | 46.4 ± 15.1 |
| Week 3 | 75.3 ± 39.0 | 79.8 ± 72.7 |
| Week 4 | 46.3 ± 13.3 | 22.4 ± 7.1 |
| Mixed model effects | p-Values | F statistics |
| Time A53T (Untreated) | 0.0069 | F(3.00,15.00) = 12.70 |
| Time A53T (LNA) | 0.0027 | F(3.00,6.74) = 44.18 |
| LNA-Treatment (L) | >0.9999 | F(1.00,24.00) = 2.93 |
| LNA x Time (LxT) | >0.9999 | F(3.00,24.00) = 0.79 |
| ISIV | A53T (Untreated) | A53T (LNA) |
| Week 1 | 3068±997 | 2248±659 |
| Week 2 | 436 ± 157 | 276 ± 147 |
| Week 3 | 567 ± 454 | 815 ± 942 |
| Week 4 | 213 ± 113 | 113 ± 67 |
| Mixed model effects | p-Values | F statistics |
| Time A53T (Untreated) | 0.0043 | F(3.00,17.00) = 12.67 |
| Time A53T (LNA) | 0.0070 | F(3.00,10.00) = 18.32 |
| LNA-Treatment (L) | >0.9999 | F(1.00,27.00) = 0.90 |
| LNA x Time (LxT) | >0.9999 | F(3.00,27.00) = 1.23 |
| ISICV | A53T (Untreated) | A53T (LNA) |
| Week 1 | 1.12 ± 0.08 | 1.14 ± 0.03 |
| Week 2 | 1.23 ± 0.14 | 1.99 ± 0.52 |
| Week 3 | 1.38 ± 0.21 | 1.96 ± 0.41 |
| Week 4 | 1.47 ± 0.09 | 1.77 ± 0.52 |
| Mixed model effects | p-Values | F statistics |
| Time A53T (Untreated) | 0.5020 | F(3.00,18.00) = 4.52 |
| Time A53T (LNA) | 0.4893 | F(3.00,7.61) = 6.75 |
| LNA-Treatment (L) | 0.0620 | F(1.00,28.00) = 11.70 |
| LNA x Time (LxT) | >0.9999 | F(3.00,28.00) = 1.82 |
| PACF | A53T (Untreated) | A53T (LNA) |
| Week 1 | -0.038 ± 0.025 | -0.034 ± 0.028 |
| Week 2 | -0.055 ± 0.047 | -0.042 ± 0.055 |
| Week 3 | -0.074 ± 0.013 | -0.0013 ± 0.0191 |
| Week 4 | -0.069 ± 0.019 | 0.018 ± 0.005 |
| Mixed model effects | p-Values | F statistics |
| Time A53T (Untreated) | >0.9999 | F(3.00,15.00) = 2.54 |
| Time A53T (LNA) | >0.9999 | F(3.00,9.00) = 0.83 |
| LNA-Treatment (L) | 0.0560 | F(1.00,24.00) = 12.40 |
| LNA x Time (LxT) | >0.9999 | F(3.00,24.00) = 2.69 |
| SCRF [Hz] | A53T (Untreated) | A53T (LNA) |
| Week 1 | 1.98 ± 0.03 | 1.69 ± 0.20 |
| Week 2 | 0.94 ± 0.51 | 0.45 ± 0.05 |
| Week 3 | 0.83 ± 0.49 | 0.74 ± 0.62 |
| Week 4 | 0.44 ± 0.09 | 0.27 ± 0.11 |
| Mixed model effects | p-Values | F statistics |
| Time A53T (Untreated) | 0.0020 | F(3.00,16.00) = 15.13 |
| Time A53T (LNA) | 0.1871 | F(3.00,7.97) = 9.15 |
| LNA-Treatment (L) | >0.9999 | F(1.00,26.00) = 3.79 |
| LNA x Time (LxT) | >0.9999 | F(3.00,26.00) = 0.45 |
| SCRM | A53T (Untreated) | A53T (LNA) |
| Week 1 | 32.4 ± 7.4 | 36.9 ± 7.6 |
| Week 2 | 92.4 ± 64.3 | 302 ± 193 |
| Week 3 | 126 ± 91 | 345 ± 236 |
| Week 4 | 193 ± 37 | 287 ± 217 |

| | | |
|---|---|---|
| Mixed model effects | p-Values | F statistics |
| Time A53T (Untreated) | >0.9999 | F(3.00,19.00) = 3.36 |
| Time A53T (LNA) | 0.9957 | F(3.00,11.00) = 4.29 |
| LNA-Treatment (L) | 0.4676 | F(1.00,30.00) = 6.72 |
| LNA x Time (LxT) | >0.9999 | F(3.00,30.00) = 1.07 |
| RFIT | A53T (Untreated) | A53T (LNA) |
| Week 1 | -0.0052 ± 0.0004 | -0.0046 ± 0.0017 |
| Week 2 | -0.013 ± 0.002 | -0.0081 ± 0.0009 |
| Week 3 | -0.010 ± 0.004 | -0.0066 ± 0.0002 |
| Week 4 | -0.011 ± 0.002 | -0.0095 ± 0.0035 |
| Mixed model effects | p-Values | F statistics |
| Time A53T (Untreated) | 0.2488 | F(3.00,12.37) = 6.30 |
| Time A53T (LNA) | 0.6239 | F(3.00,10.00) = 5.26 |
| LNA-Treatment (L) | 0.1738 | F(1.00,27.00) = 9.14 |
| LNA x Time (LxT) | >0.9999 | F(3.00,27.00) = 1.39 |
| IBIV | A53T (Untreated) | A53T (LNA) |
| Week 1 | 97.7 ± 94.8 | 21.5 ± 12.4 |
| Week 2 | 2.16 ± 1.34 | 0.32 ± 0.20 |
| Week 3 | 1.99 ± 1.80 | 14.1 ± 16.9 |
| Week 4 | 0.89 ± 1.20 | 6.16 ± 6.10 |
| Mixed model effects | p-Values | F statistics |
| Time A53T (Untreated) | 0.7707 | F(3.00,12.24) = 4.50 |
| Time A53T (LNA) | >0.9999 | F(3.00,6.25) = 3.07 |
| LNA-Treatment (L) | >0.9999 | F(1.00,6.68) = 1.90 |
| LNA x Time (LxT) | 0.7201 | F(3.00,18.33) = 4.06 |
| IBIM [s] | A53T (Untreated) | A53T (LNA) |
| Week 1 | 22.4 ± 8.8 | 13.4 ± 6.5 |
| Week 2 | 9.58 ± 3.72 | 14.4 ± 2.8 |
| Week 3 | 8.11 ± 0.55 | 11.5 ± 1.1 |
| Week 4 | 7.72 ± 0.05 | 11.3 ± 1.7 |
| Mixed model effects | p-Values | F statistics |
| Time A53T (Untreated) | >0.9999 | F(3.00,13.74) = 0.84 |
| Time A53T (LNA) | 0.3491 | F(3.00,6.96) = 8.22 |
| LNA-Treatment (L) | >0.9999 | F(1.00,7.39) = 0.27 |
| LNA x Time (LxT) | 0.1743 | F(3.00,20.21) = 5.69 |
| MBD [s] | A53T (Untreated) | A53T (LNA) |
| Week 1 | 3.41 ± 2.00 | 4.94 ± 1.81 |
| Week 2 | 8.30 ± 0.12 | 7.50 ± 0.06 |
| Week 3 | 7.32 ± 0.44 | 6.39 ± 0.92 |
| Week 4 | 6.34 ± 0.51 | 6.42 ± 0.84 |
| Mixed model effects | p-Values | F statistics |
| Time A53T (Untreated) | 0.7940 | F(3.00,9.04) = 5.08 |
| Time A53T (LNA) | 0.4152 | F(3.00,5.02) = 10.63 |
| LNA-Treatment (L) | >0.9999 | F(1.00,5.71) = 0.01 |
| LNA x Time (LxT) | >0.9999 | F(3.00,14.12) = 2.56 |
| VBD | A53T (Untreated) | A53T (LNA) |
| Week 1 | 4.29 ± 4.01 | 7.07 ± 3.17 |
| Week 2 | 0.77 ± 0.37 | 0.33 ± 0.22 |
| Week 3 | 0.81 ± 0.14 | 1.25 ± 1.62 |
| Week 4 | 0.28 ± 0.33 | 0.42 ± 0.39 |
| Mixed model effects | p-Values | F statistics |
| Time A53T (Untreated) | 0.0013 | F(3.00,16.00) = 16.27 |
| Time A53T (LNA) | >0.9999 | F(3.00,8.00) = 1.95 |
| LNA-Treatment (L) | >0.9999 | F(1.00,24.00) = 0.92 |
| LNA x Time (LxT) | >0.9999 | F(3.00,24.00) = 1.01 |

| | | |
|---|---|---|
| INTRABF [Hz] | A53T (Untreated) | A53T (LNA) |
| Week 1 | 0.23 ± 0.05 | 0.23 ± 0.04 |
| Week 2 | 0.37 ± 0.14 | 1.57 ± 0.97 |
| Week 3 | 0.64 ± 0.38 | 1.75 ± 0.78 |
| Week 4 | 0.77 ± 0.02 | 1.61 ± 0.94 |
| Mixed model effects | p-Values | F statistics |
| Time A53T (Untreated) | 0.3067 | F(3.00,19.00) = 5.06 |
| Time A53T (LNA) | >0.9999 | F(3.00,6.46) = 4.71 |
| LNA-Treatment (L) | 0.0624 | F(1.00,28.00) = 11.68 |
| LNA x Time (LxT) | >0.9999 | F(3.00,28.00) = 1.64 |
| INTERBF [Hz] | A53T (Untreated) | A53T (LNA) |
| Week 1 | 0.10 ± 0.02 | 0.12 ± 0.02 |
| Week 2 | 0.12 ± 0.02 | 0.17 ± 0.04 |
| Week 3 | 0.15 ± 0.06 | 0.24 ± 0.07 |
| Week 4 | 0.21 ± 0.03 | 0.28 ± 0.17 |
| Mixed model effects | p-Values | F statistics |
| Time A53T (Untreated) | >0.9999 | F(3.00,12.96) = 3.61 |
| Time A53T (LNA) | 0.5083 | F(3.00,8.39) = 6.24 |
| LNA-Treatment (L) | >0.9999 | F(1.00,7.02) = 4.23 |
| LNA x Time (LxT) | >0.9999 | F(3.00,21.01) = 0.57 |
| BRT [s] | A53T (Untreated) | A53T (LNA) |
| Week 1 | 0.57 ± 0.36 | 1.38 ± 0.69 |
| Week 2 | 2.68 ± 0.10 | 2.27 ± 0.16 |
| Week 3 | 2.74 ± 0.76 | 1.59 ± 0.23 |
| Week 4 | 2.74 ± 1.02 | 2.09 ± 1.28 |
| Mixed model effects | p-Values | F statistics |
| Time A53T (Untreated) | >0.9999 | F(3.00,17.00) = 1.72 |
| Time A53T (LNA) | 0.2253 | F(3.00,9.00) = 7.83 |
| LNA-Treatment (L) | >0.9999 | F(1.00,26.00) = 1.94 |
| LNA x Time (LxT) | >0.9999 | F(3.00,26.00) = 2.82 |
| BRV [coactivity/s] | A53T (Untreated) | A53T (LNA) |
| Week 1 | 27.2 ± 10.5 | 20.1 ± 12.4 |
| Week 2 | 17.0 ± 3.9 | 50.0 ± 12.2 |
| Week 3 | 41.7 ± 25.9 | 164 ± 140 |
| Week 4 | 61.1 ± 24.0 | 144 ± 3 |
| Mixed model effects | p-Values | F statistics |
| Time A53T (Untreated) | 0.6885 | F(3.00,16.00) = 4.27 |
| Time A53T (LNA) | >0.9999 | F(3.00,10.00) = 3.24 |
| LNA-Treatment (L) | 0.5006 | F(1.00,26.00) = 6.69 |
| LNA x Time (LxT) | >0.9999 | F(3.00,26.00) = 1.88 |
| BDT [s] | A53T (Untreated) | A53T (LNA) |
| Week 1 | 0.60 ± 0.46 | 0.95 ± 0.43 |
| Week 2 | 2.37 ± 0.25 | 1.60 ± 0.12 |
| Week 3 | 2.23 ± 0.29 | 1.38 ± 0.26 |
| Week 4 | 1.71 ± 0.52 | 1.51 ± 0.03 |
| Mixed model effects | p-Values | F statistics |
| Time A53T (Untreated) | 0.2196 | F(3.00,16.00) = 5.83 |
| Time A53T (LNA) | 0.0133 | F(3.00,10.00) = 15.66 |
| LNA-Treatment (L) | 0.1211 | F(1.00,26.00) = 10.11 |
| LNA x Time (LxT) | 0.0530 | F(3.00,26.00) = 6.72 |
| BDV [coactivity/s] | A53T (Untreated) | A53T (LNA) |
| Week 1 | -30.7 ± 12.1 | -21.6 ± 8.6 |
| Week 2 | -21.2 ± 4.4 | -87.1 ± 38.8 |
| Week 3 | -46.0 ± 25.2 | -176 ± 139 |

| | | |
|---|---|---|
| Week 4 | -67.1 ± 0.2 | -282 ± 24 |
| Mixed model effects | p-Values | F statistics |
| Time A53T (Untreated) | 0.0316 | F(3.00,17.00) = 8.75 |
| Time A53T (LNA) | >0.9999 | F(3.00,6.51) = 4.68 |
| LNA-Treatment (L) | 0.0078 | F(1.00,26.00) = 18.04 |
| LNA x Time (LxT) | 0.7022 | F(3.00,26.00) = 3.80 |
| SYNC | A53T (Untreated) | A53T (LNA) |
| Week 1 | 0.0023 ± 0.0002 | 0.0029 ± 0.0005 |
| Week 2 | 0.0046 ± 0.0008 | 0.0062 ± 0.0006 |
| Week 3 | 0.0046 ± 0.0006 | 0.0069 ± 0.0015 |
| Week 4 | 0.0049 ± 0.0003 | 0.0061 ± 0.0006 |
| Mixed model effects | p-Values | F statistics |
| Time A53T (Untreated) | 0.0002 | F(3.00,17.00) = 20.97 |
| Time A53T (LNA) | 0.0263 | F(3.00,7.63) = 17.80 |
| LNA-Treatment (L) | 0.0008 | F(1.00,27.00) = 25.76 |
| LNA x Time (LxT) | >0.9999 | F(3.00,27.00) = 1.60 |
| NRF [Hz] | A53T (Untreated) | A53T (LNA) |
| Week 1 | 0.046 ± 0.006 | 0.046 ± 0.006 |
| Week 2 | 0.052 ± 0.006 | 0.047 ± 0.006 |
| Week 3 | 0.064 ± 0.003 | 0.047 ± 0.011 |
| Week 4 | 0.063 ± 0.027 | 0.049 ± 0.010 |
| Mixed model effects | p-Values | F statistics |
| Time A53T (Untreated) | >0.9999 | F(3.00,14.08) = 0.21 |
| Time A53T (LNA) | >0.9999 | F(3.00,9.00) = 1.46 |
| LNA-Treatment (L) | >0.9999 | F(1.00,7.80) = 3.29 |
| LNA x Time (LxT) | >0.9999 | F(3.00,20.78) = 1.35 |
| NRM | A53T (Untreated) | A53T (LNA) |
| Week 1 | 176 ± 50 | 250 ± 53 |
| Week 2 | 472 ± 70 | 526 ± 118 |
| Week 3 | 462 ± 287 | 327 ± 212 |
| Week 4 | 546 ± 252 | 323 ± 207 |
| Mixed model effects | p-Values | F statistics |
| Time A53T (Untreated) | >0.9999 | F(3.00,14.53) = 3.52 |
| Time A53T (LNA) | >0.9999 | F(3.00,10.00) = 2.51 |
| LNA-Treatment (L) | >0.9999 | F(1.00,29.00) = 0.95 |
| LNA x Time (LxT) | >0.9999 | F(3.00,29.00) = 1.45 |
| NRFIT | A53T (Untreated) | A53T (LNA) |
| Week 1 | -0.032 ± 0.020 | -0.015 ± 0.005 |
| Week 2 | -0.0052 ± 0.0018 | -0.0029 ± 0.0003 |
| Week 3 | -0.0042 ± 0.0026 | -0.0019 ± 0.0002 |
| Week 4 | -0.0032 ± 0.0014 | -0.0024 ± 0.0005 |
| Mixed model effects | p-Values | F statistics |
| Time A53T (Untreated) | <0.0001 | F(3.00,13.45) = 39.01 |
| Time A53T (LNA) | 0.3780 | F(3.00,7.58) = 7.45 |
| LNA-Treatment (L) | >0.9999 | F(1.00,6.55) = 4.93 |
| LNA x Time (LxT) | >0.9999 | F(3.00,20.11) = 3.05 |

**Supplemental table 15.** Complementary data related to Figure 5. Reported are Mean ± SD of all features for each week and genotype and the results of linear mixed-effect models assessing LNA-treatment (L) and LNA-treatment-by-time interaction (LxT). p-Values were adjusted for the number of comparisons (Bonferroni correction). Sample size for these analyses were 6 untreated/non-targeted LNA-treated A53T and 4 LNA-treated A53T cultures.

| Random Forest | |
|---|---|
| MinLeafSize | range(1,max(2, floor(NumObservations/2))) |
| MaxNumSplits | range(1, max(2,NumObservations-1)) |
| NumLearningCycles | range(10, 500) |
| SplitCriterion | gdi, deviance, twoing |
| NumVariablesToSample | range(1, max(2,NumPredictors)) |

| SVM | |
|---|---|
| BoxConstraint | range(0.1, 1000) |
| KernelFunction | gaussian, linear, polynomial |
| KernelScale | range(0.1, 1000) |
| PolynomialOrder | 2, 3, 4 |

| CNB | |
|---|---|
| DistributionNames | normal, kernel |
| Kernel | normal, box, epanechnikov, triangle |
| Width | range(MinPredictorDiff/4, max(MaxPredictorRange,MinPredictorDiff)) |

| KNN | |
|---|---|
| Distance | cityblock, chebychev, correlation, cosine, Euclidean, hamming, jaccard, mahalanobis, minkowski, seuclidean, spearman |
| Exponent | range(0.5,3) |
| DistanceWeight | equal, inverse, squaredinverse |
| NumNeighbors | range (1, max(2,round(NumObservations/2))) |

**Supplemental table 16.** Hyperparameter optimization options for the different classifiers used. The left column lists the hyperparameters that were optimized, the right column the respective values among which the optimization searched. Further information about the hyperparameters is available on the Matlab documentation.

| **Primary antibody** | **Dilution** | **Catalogue number** |
|---|---|---|
| mouse anti-TH | 1:500 | #MAB318, Sigma-Aldrich |
| chicken anti-MAP2 | 1:1000 | #CH22103, Neuromics (Edina, MN, USA) |
| rabbit anti-GFAP | 1:500 | #Z0334, Agilent (Santa Clara, CA, USA) |
| rabbit anti-phospho-α-synuclein | 1:100 | Britschgi lab, Roche (Basel, Switzerland) |
| rabbit anti-α-synuclein | 1:100 | #2628, Cell Signaling Technology (Danvers, MS, USA) |
| **Secondary antibody** | | |
| donkey anti-mouse 488 | 1:250 | #A-21202, ThermoFisher |
| goat anti-chicken 647 | 1:500 | #A-32933, ThermoFisher |
| donkey anti-rabbit 568 | 1:250 | #A-A10042, ThermoFisher |

**Supplemental table 17.** Primary and secondary antibodies.
